# Hippocampal subfield volumes are uniquely affected in PTSD and depression: International analysis of 31 cohorts from the PGC-ENIGMA PTSD Working Group

**DOI:** 10.1101/739094

**Authors:** Lauren E. Salminen, Philipp G. Sämann, Yuanchao Zheng, Emily L. Dennis, Emily K. Clarke-Rubright, Neda Jahanshad, Juan E. Iglesias, Christopher D. Whelan, Steven E. Bruce, Jasmeet P. Hayes, Soraya Seedat, Christopher L. Averill, Lee A. Baugh, Jessica Bomyea, Joanna Bright, Chanellé J. Buckle, Kyle Choi, Nicholas D. Davenport, Richard J. Davidson, Maria Densmore, Seth G. Disner, Stefan du Plessis, Jeremy A. Elman, Negar Fani, Gina L. Forster, Carol E. Franz, Jessie L. Frijling, Atilla Gonenc, Staci A. Gruber, Daniel W. Grupe, Jeffrey P. Guenette, Courtney C. Haswell, David Hofmann, Michael Hollifield, Babok Hosseini, Anna R. Hudson, Jonathan Ipser, Tanja Jovanovic, Amy Kennedy-Krage, Mitzy Kennis, Anthony King, Philipp Kinzel, Saskia B. J. Koch, Inga Koerte, Sheri M. Koopowitz, Mayuresh S. Korgaonkar, William S. Kremen, John Krystal, Lauren A. M. Lebois, Ifat Levy, Michael J. Lyons, Vincent A. Magnotta, Antje Manthey, Soichiro Nakahara, Laura Nawijn, Richard W. J. Neufeld, Jack B. Nitschke, Daniel C. M. O’Doherty, Robert H. Paul, Matthew Peverill, Faisal M. Rashid, Kerry J. Ressler, Annerine Roos, Christian Schmahl, Margaret A. Sheridan, Anika Sierk, Alan N. Simmons, Jeffrey S. Simons, Raluca M. Simons, Murray B. Stein, Jennifer S. Stevens, Benjamin Suarez-Jimenez, Jean Théberge, Kathleen Thomaes, Sophia I. Thomopoulos, Leigh L. van den Heuvel, Steven J. A. van der Werff, Theo G. M. van Erp, Sanne J. H. van Rooij, Mirjam van Zuiden, Tim Varkevisser, Robert R. J. M. Vermeiren, Tor D. Wager, Henrik Walter, Xin Wang, Sherry Winternitz, Jonathan D. Wolff, Kristen Wrocklage, Xi Zhu, Christopher R. K. Ching, Tiril P. Gurholt, Unn K. Haukvik, Ingrid Agartz, Chadi G. Abdallah, Richard Bryant, Judith K. Daniels, Michael DeBellis, Kelene A. Fercho, Elbert Geuze, Ilan Harpaz-Rotem, Julia I. Herzog, Milissa L. Kaufman, Jim Lagopoulos, Ruth A. Lanius, Katie A. McLaughlin, Sven C. Mueller, Yuval Neria, Miranda Olff, K. Luan Phan, Martha E. Shenton, Scott R. Sponheim, Dan J. Stein, Thomas Straube, Nic J. A. van der Wee, Dick J. Veltman, Paul M. Thompson, Rajendra A. Morey, Mark W. Logue, for the ENIGMA-PGC-PTSD Working Group

**Author notes:** Denotes equal contribution (Rajendra A. Morey, Mark W. Logue).

## Abstract

**Background:** PTSD and depression commonly co-occur and have been associated with smaller hippocampal volumes compared to healthy and trauma-exposed controls. However, the hippocampus is heterogeneous, with subregions that may be uniquely affected in individuals with PTSD and depression.

**Methods:** We used random effects regressions and a harmonized neuroimaging protocol based on FreeSurfer (v6.0) to identify sub-structural hippocampal markers of current PTSD (C-PTSD), depression, and the interaction of these conditions across 31 cohorts worldwide (N=3,115; *M*_age_=38.9±13.9 years). Secondary analyses tested these associations by sex and after modeling the simultaneous effects of remitted PTSD, childhood trauma, mild traumatic brain injury, and alcohol use disorder.

**Results:** A significant negative main effect of depression (n=800, vs. no depression, n=1456) was observed in the hippocampal tail (ß=−0.13) and CA1 (ß=−0.09) after adjusting for covariates and multiple testing (adjusted p’s (q)=0.028). A main effect of C-PTSD (n=1042 vs. control, n=1359) was not significant, but an interaction between C-PTSD and depression was significant in the CA1 (ß=−0.24, q=0.044). Pairwise comparisons revealed significantly smaller CA1 volumes in individuals with C-PTSD+Depression than controls (ß=−0.12, q=0.012), C-PTSD-only (ß=−0.17, q=0.001), and Depression-only (ß=−0.18, q=0.023). Follow-up analyses revealed sex effects in the hippocampal tail of depressed females, and an interaction effect of C-PTSD and depression in the fimbria of males.

**Conclusions:** Collectively our results suggest that depression is a stronger predictor of hippocampal volumetry than PTSD, particularly in the CA1, and provide compelling evidence of more pronounced hippocampal phenotypes in comorbid PTSD and depression compared to either condition alone.

## Introduction

Posttraumatic stress disorder (PTSD) is a serious mental health condition characterized by intrusive, avoidant, cognitive, and affective symptoms that emerge after a precipitating traumatic event and interfere with activities of daily living (1). Approximately 70% of the global population will experience one or more traumatic events at some point during their lifetime, but the prevalence of lifetime PTSD is only 8% worldwide (2). The mechanisms that underlie this discrepancy likely result from complex individual differences in genetic, environmental, and biopsychosocial factors that differentially influence PTSD susceptibility (3, 4).

PTSD has been linked to structural and functional brain changes via dysregulation of the hypothalamic-pituitary-adrenal (HPA) axis - a negative feedback loop that promotes adaptive responses to actual or perceived threat (5, 6). During acute stress, the hypothalamus rapidly activates the sympathetic nervous system to mobilize energy through the release of catecholamines (7, 8). Subsequently, the HPA axis is activated and corticotropin-releasing hormone (CRH) is released from the hypothalamus into the hypophyseal portal circulation. CRH potentiates the release of adrenocorticotropic hormone (ACTH) from the anterior pituitary, signaling glucocorticoid (GC) synthesis from the adrenals to the systemic circulation (8). GCs bind to low-affinity GC receptors (GRs) in the brain during stress, with the highest maximal binding capacity in the hippocampus (6). GRs stimulate rapid non-genomic self-defense processes (e.g., immune activity) as well as long-term changes in gene expression. Of critical importance, GRs activate GABAergic interneurons to mediate fast-feedback inhibition of the HPA axis once a stressor has ended to promote recovery from acute challenge (8–11).

Chronic stress, however, disrupts hippocampal feedback control of the hypothalamus, leading to prolonged HPA activation and epigenetic changes (8, 12, 13). Animal studies show that chronic stress is associated with dendritic atrophy and loss of spine density in distinct subregions of the hippocampus (12, 14, 15). As PTSD is associated with altered HPA axis regulation and GR signaling (16), this may explain more than two dozen studies that replicate smaller hippocampal volumes in PTSD (17–27), including a recent large-scale meta-analysis from our group (27). Hippocampal atrophy prior to the precipitating traumatic event may also increase vulnerability to PTSD (28–30). This is supported by neuroendocrine studies showing higher immune GR counts in pre-deployed soldiers who developed severe PTSD symptoms within 6 months post-deployment (31), and lower cortisol in the acute aftermath of trauma victims who developed PTSD symptoms compared to those who did not (32–34). Thus, low cortisol during traumatic stress recovery may result from pre-existing HPA axis dysregulation that primes hippocampal atrophy and subsequent development of PTSD.

Increasing evidence suggests that the effects of stress vary across subdivisions of the hippocampus due to their unique functional and cytoarchitectural properties (35). Indeed, previous studies reported smaller volumes in subregions of the *cornu ammonis* (CA), CA1 (36) and CA3 (37, 38), dentate gyrus (DG) (26, 37), subiculum (37), and hippocampal-amygdala-transition-area (HATA) (39, 40) of PTSD patients compared to controls. Hence, hippocampal volume differences in PTSD may be driven by effects in specific subregions that inform the signature of traumatic stress in the hippocampus and corresponding clinical symptoms. Concurrent clinical and sociodemographic factors may also exert independent or additive effects on substructural hippocampal volumes. In particular, depression occurs in 30-60% of PTSD patients and has been associated with reduced integrity of the same subfields implicated in PTSD (35, 41–44). Additional factors such as childhood trauma, alcohol use disorder (AUD), and traumatic brain injury (TBI) may alter the hippocampus but are variably represented across PTSD samples. To date, the relative influence of PTSD, depression, and other clinical variables on subfield volumetry has not been examined in a well-powered study.

Achieving sufficient power to analyze the shared and unique hippocampal correlates of PTSD, depression, and other intervening factors is challenging for prospective studies given the high comorbidity rates and high cost of MRI scans. Analyses of large multi-cohort samples using standardized pipelines offer greater reproducibility, are more generalizable, and are less likely to contribute to the growing number of contradictory reports that have plagued the literature (45). The present study leveraged these methods through a joint partnership between the Psychiatric Genetics Consortium (PGC; https://www.med.unc.edu/pgc/) and Enhancing NeuroImaging Genetics through Meta-Analysis (ENIGMA; http://enigma.ini.usc.edu) consortium to identify substructural hippocampal signatures of PTSD and depression through harmonized mega-analyses of 31 international cohorts.

## Methods

### Participants

Subject-level imaging and clinical data were obtained from 31 cohorts from seven countries (N=3115). Exclusion and inclusion criteria for each cohort are in **Supplementary Table 1**. We further excluded participants with any comorbid Axis I or II psychiatric disorder, except PTSD, depression, and AUD. Individuals with known moderate to severe TBI were also excluded. Sample characteristics are reported in **Table 1** and **Supplementary Table 2**. All participants provided written informed consent approved by the local Institutional Review Board of each cohort.

**Table 1.** Descriptive characteristics of the sample (with ethnicity data)

|  | N,<br>data available | Current PTSD | Control | Total |
| --- | --- | --- | --- | --- |
| N | -- | 1047 | 1361 | 2408 |
| Age (M, SD) | 2408 | 38.0 (13.0) *** | 41.1 (15.3) *** | 39.8 (14.4) |
| Sex (% Female) | 2408 | 42.3** | 35.8** | 38.6 |
| Ethnicity | 2408 | -- | -- | -- |
| • % White/Caucasian | -- | 66.8 | 67.2 | 67.0 |
| • % Black/African American | -- | 14.8 | 16.5 | 15.7 |
| • % Multiracial | -- | 12.9 | 12.3 | 12.5 |

<sup>1</sup> The hippocampal extended measurement is equivalent to that of whole hippocampal volume. This summed measurement is different than the measure of whole hippocampal volume in Logue et al. (27), which used the autosegmentation 'aseg' output from FreeSurfer 5.3 to calculate whole hippocampal volume.

|  |  |  |  |  |
| --- | --- | --- | --- | --- |
| • % Other | -- | 5.5 | 4.0 | 4.7 |
| % Depression | 2263 | 61.8*** | 11.0*** | 33.6 |
| % Military | 2369 | 44.7** | 50.7** | 48.1 |
| % mTBI | 785 | 68.1*** | 44.2*** | 51.5 |
| % AUD | 1616 | 18.5*** | 8.8*** | 13.0 |
| Childhood trauma | 940 | -- | -- | -- |
| • % 1 type | -- | 16.5*** | 18.9*** | 18.0 |
| • % 2+ types | -- | 64.0*** | 25.5*** | 40.9 |
| % Taking psychotropic medication | 1169 | 36.8*** | 9.4*** | 21.5 |
| % Trauma-exposed (controls) | 1353 | -- | 81.1 | 81.1 |
| % Lifetime PTSD (controls) | 1667 | -- | 24.6 | 24.6 |

*Note.* Proportions were calculated in participants with information on ethnicity, and without remitted PTSD to provide the most representative depiction of sociodemographic factors in the sample that was used for the majority of study analyses. Lifetime PTSD proportions were calculated in individuals with ethnicity data. Proportions were derived using chi square, age comparisons were derived using t-tests. AUD= alcohol use disorder, mTBI= mild traumatic brain injury. For detailed information by cohort see **Supplementary Table 3**. \* $p < 0.05$ , \*\* $p < 0.01$ , \*\*\* $p < 0.001$

### Assessment of PTSD and Depression

PTSD diagnoses were determined using one of seven instruments (**Supplementary Table 3**). Current PTSD (C-PTSD) was our primary index of PTSD. Current symptom severity was measured using total scores on the Clinician Administered PTSD Scale (CAPS), which were available in 50% of the sample. Current depression was assessed using a harmonized index of published thresholds (**Supplementary Table 3**). The Beck Depression Inventory-II (BDI) was the most frequently used instrument for depression assessment. Total BDI scores were used as our index of depression severity. Assessment and harmonization of other clinical variables are described in the Supplement.

### Neuroimaging Approach

Hippocampal subfield volumes were identically generated across cohorts using the ENIGMA hippocampal subfield extraction and quality control protocol (http://enigma.ini.usc.edu/ongoing/enigma-hippocampal-subfields/) based on FreeSurfer v.6 (FS6) (46). This protocol yields 12 bilateral subfields including the CA1, combined CA2/CA3 (referred to as CA3), CA4, granule cell layer of the DG (referred to as DG), HATA, presubiculum, subiculum, parasubiculum, molecular layer (MOL), fimbria, fissure, and tail (**Figure 1**). The fissure is a thin band of cerebrospinal fluid that was used for quality control, but not analyzed as a dependent variable. Additional details are presented in the Supplement.

**Figure 1.**
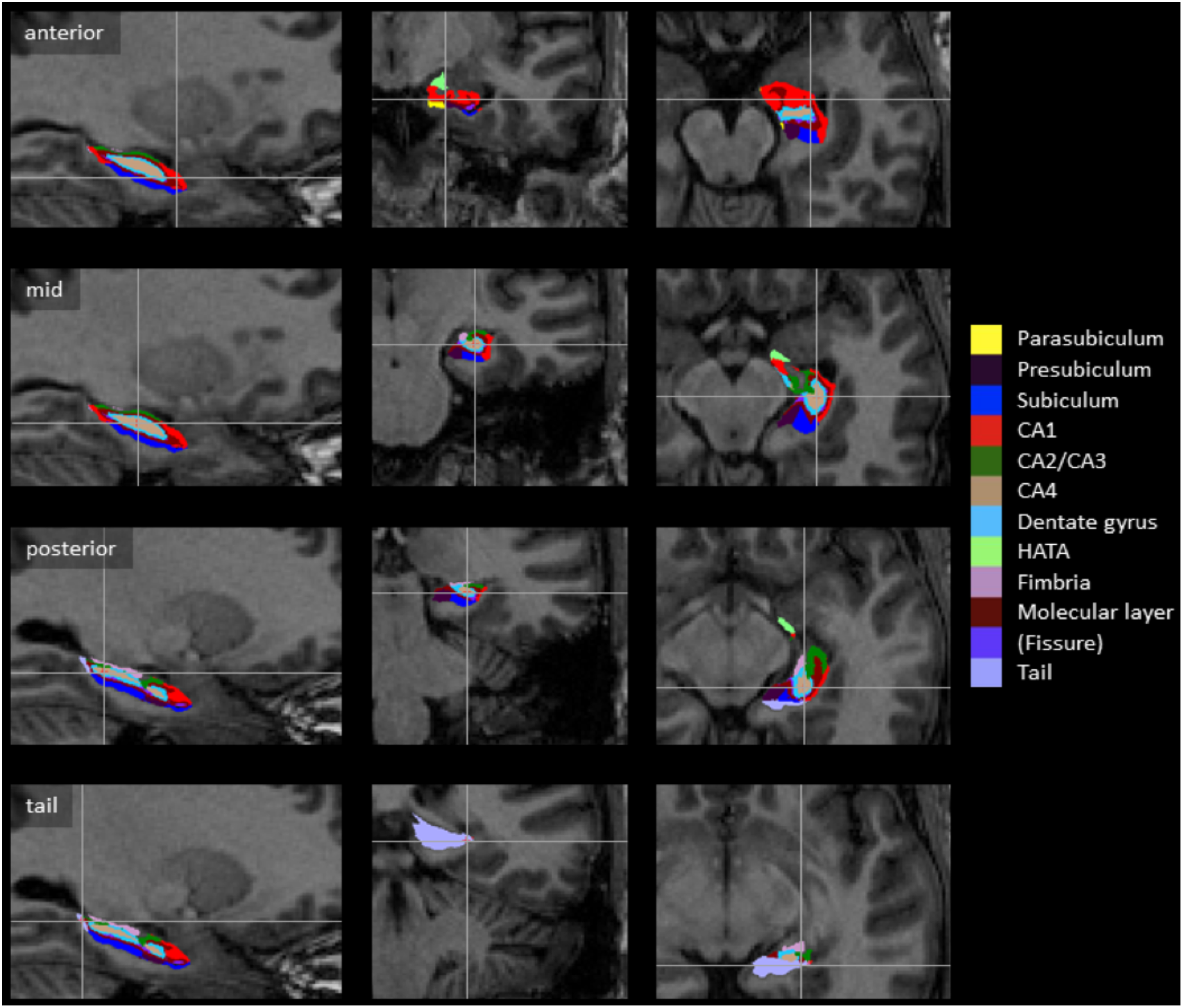
Example of automated FreeSurfer (v6.0) hippocampal subfield segmentation. Binarized subregions are overlaid on the bias corrected T1-weighted input image in native space. Rows shows different coronal positions (head, anterior body, posterior body and tail area) with color coded subregions; columns show sagittal, coronal and axial views with corresponding cross-hair (radiological view, i.e., subregions of a left hippocampus). Note the slim hippocampal fissure (dark purple), representing cerebrospinal fluid and not counted as parenchyma. Note anterior location of CA1 (red) and the hippocampal-amygdala-transition area (HATA; light green), both bordering the amygdala, and the central location of CA4, the superior position of CA2/CA3 and inferior position of the subiculum. Details regarding subfield descriptions are in the main text and Supplementary methods. Information on re-aggregation of subregions to composite subregions is described in **Supplementary Table 5.**

### Design

We used the lmerTest package in R to compute random-effects regressions for all analyses. The model included random effects for the study cohort and MRI scanner manufacturer, which was modeled as a nested variable within cohort for cohorts that used more than one scanner. Fixed effects included age, age^2^, sex, age*sex, age^2^*sex, ethnicity, and ICV (**Supplementary Table 5**). Civilian/military background was strongly correlated with sex (phi=0.631, p<0.001) and thus was not included as a covariate in our main models. Analyses were conducted using two-tailed tests; multiple comparisons were controlled within each set of analyses using the false discovery rate (FDR) procedure (47). We refer to statistical results with FDR-corrected p-values (q)>0.05, but q<0.1, as “trends”. A brief summary of analysis models is below, with detailed descriptions in the Supplement.

### Statistical Models

#### Main effects of PTSD and depression

The effect of C-PTSD on hippocampal subfield volumes was tested against all controls and trauma-exposed controls only. C-PTSD severity was analyzed in all participants with CAPS data and separately in the C-PTSD group. Additional analyses examined the effects of trauma exposure in the control group, and of L-PTSD and R-PTSD. We also tested the effects of C-PTSD after covarying for childhood trauma, mild TBI, AUD, and use of psychotropic medication.

Main effects of depression and depression severity were tested without covarying for C-PTSD to check for significant effects of depression despite PTSD-related variance in the target groups. The relationship between depression severity and hippocampal subfield volume was tested in all participants with BDI data.

#### Joint models of C-PTSD and depression

First, main effects of C-PTSD were analyzed after covarying for depression, and then C-PTSD*depression interaction terms were computed for each ROI. We examined the nature of these interactions by comparing pairwise differences in least square means between the four subgroups: 1) C-PTSD+Depression (n=621), 2) Depression-only (n=138), 3) C-PTSD-only (n=384), and 4) controls (n=1,120). A control was defined as a participant without PTSD or depression. To determine whether significant interactions were driven by symptom severity, we performed moderation analyses between the binary measure of depression and total CAPS score, and between the binary C-PTSD variable and total BDI score.

To detect any bias due to the use of a mega-analysis of heterogeneous groups, we duplicated the analysis for PTSD main effects, depression main effects, and the C-PTSD*Depression interaction using meta-analysis. The methods and results of these analyses are described in the Supplement.

#### Sensitivity analysis of composite subregions

To confirm the regional specificity of our findings, we performed post-hoc comparisons of combinations of subfields using recent methods outlined by Roddy et al. (44). We created the following nine composites based on structure-function synergies defined in the literature (39, 44, 46, 48): 1) hippocampal proper (HP; CA1-4); 2) hippocampal formation (HF; CA1-4/DG/subiculum/tail); 3) CA/DG (CA1-4/DG); 4) complete dentate (CA4/DG); 5) CA-only (CA1-3); 6) Sub-complex (subiculum, presubiculum, parasubiculum); 7) CA1/sub (CA1+subiculum); 8) CA/SubPresub/MOL (CA1-3, subiculum, presubiculum, MOL); and 9) hippocampal extended (sum of all subfields minus the fissure).^1^ Additional details are in the Supplement.

#### Effects by sociodemographic status

Main effects and interactions between C-PTSD and depression also were tested separately in males, females, military, and civilians, and also in ethnicity-unadjusted analyses for comparison to our main findings.

## Results

### Main effects of PTSD

There were no FDR-corrected significant differences in hippocampal subfield volumes between the C-PTSD group and all controls. Trends for smaller volumes were observed in the CA1 (ß=−0.07, p=0.024, q=0.088), tail (ß=−0.09, p=0.017, q=0.088), presubiculum (ß =−0.07, p=0.035, q=0.096), and MOL (ß=−0.08, p=0.016, q=0.088) in those with C-PTSD (**Figure 2**). Subfield volumes did not differ between C-PTSD and trauma-exposed control groups or correspond to C-PTSD symptom severity indexed with the CAPS (**Supplementary Tables 7-8**).

**Figure 2.**
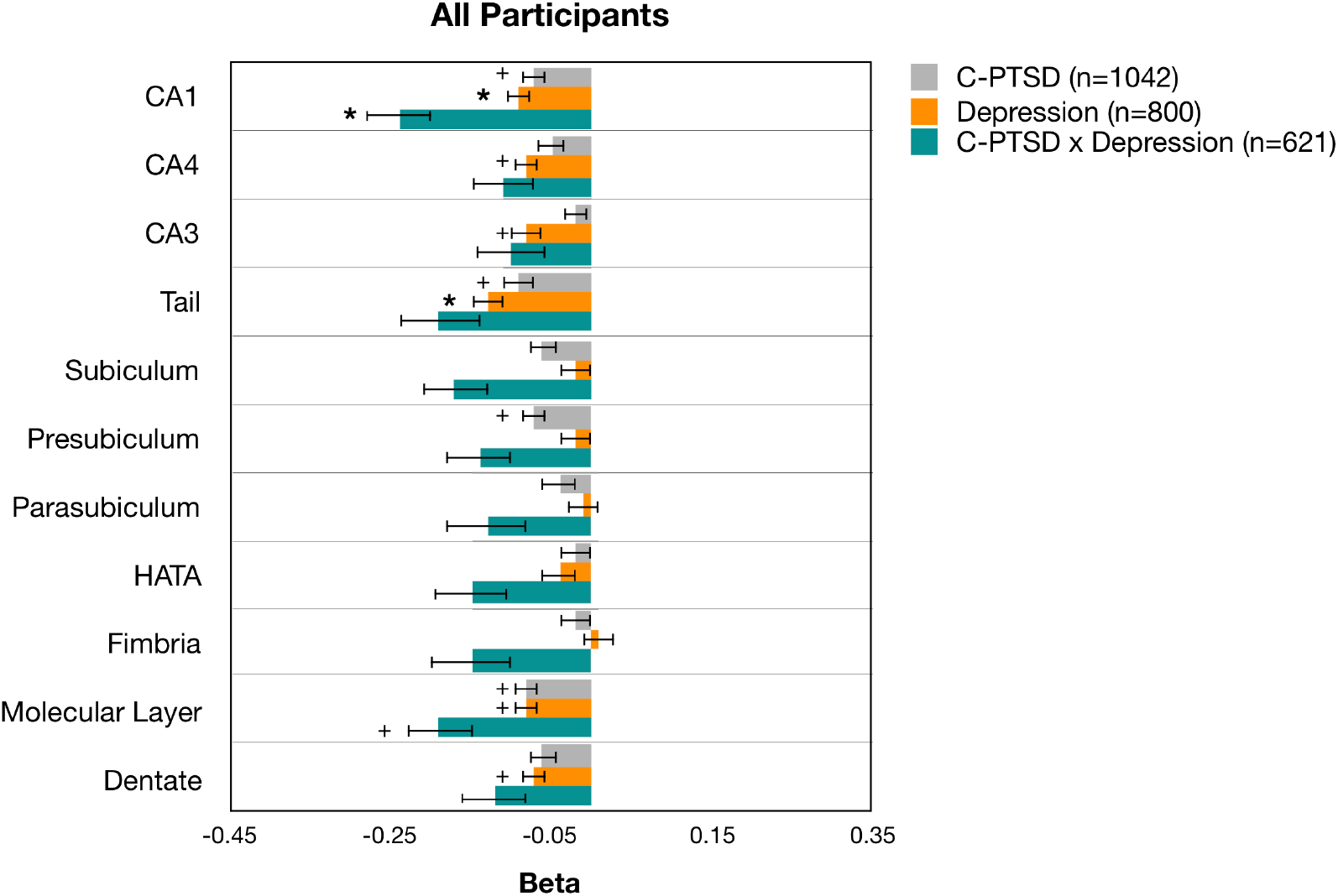
Effect sizes of C-PTSD, depression, and their interactions in all participants. Analyses of C-PTSD, Depression, and C-PTSD × Depression interactions were computed in separate models. The beta weights of those terms were plotted on the same graph for comparison. Analyses of C-PTSD did not covary for Depression, and analyses of Depression did not covary for C-PTSD. For all analyses, the sample sizes of C-PTSD and Depression controls were n=1359 and n=1456, respectively. Sample sizes for pairwise reference groups of the interaction term: C-PTSD-only, n=384; Depression-only, n=138; Control, n=1120. Significance and trend effects were determined using FDR-adjusted p-values (q). + = 0.05<q<0.10, *= 0.01<q<0.05, ** = q<0.01, ***=q<0.001.

Hippocampal subfields were not significantly affected by trauma exposure (in controls). Individuals with L-PTSD showed trends for smaller volumes in the CA1, subiculum, and MOL, and separate analyses showed a negative trend of R-PTSD in the subiculum (**Supplementary Tables 9-10, Supplementary Figure 1**). Main effects of C-PTSD were not significant in analyses adjusted for hippocampal volume rather than ICV. Other clinical covariates did not affect associations between PTSD and subfield volumes (**Supplementary Tables 11-15**).

### Main effects of depression

Individuals with depression (vs. no depression) had significantly smaller volumes in the CA1 (ß=−0.09, p=0.006, q=0.033) and tail (ß=−0.13, p=0.001, q=0.011), with trends for smaller volumes in the CA3, CA4, MOL, and DG (**Figure 2**). Higher BDI scores were significantly associated with smaller volumes in the tail (ß=−0.10, p=0.003, q=0.033), with trend effects in several other regions. There was no effect of BDI scores within the depressed group (**Supplementary Tables 16-17**).

### Joint models of PTSD and depression

Simultaneously modeling C-PTSD and depression revealed no significant effects of either condition on any subregion (**Supplementary Table 16B, C**). However, subsequent analyses including a C-PTSD*Depression interaction term revealed a significant interaction in the CA1(ß=−0.24, p=0.004, q=0.044; **Figure 2**, **Supplementary Tables 18-19**). Follow-up pairwise comparisons in the CA1 yielded significant negative effects of depression in individuals with C-PTSD (ß=−0.17, p=0.001, q=0.006), negative effects of C-PTSD in individuals with depression (ß=−0.18, p=0.009, q=0.018), and negative effects of C-PTSD+Depression compared to controls (ß=−0.12, p=0.003, q=0.009; **Figure 3A**). Depression or C-PTSD did not moderate symptom severity in any subregion (**Supplementary Table 20**).

**Figure 3A-B.**
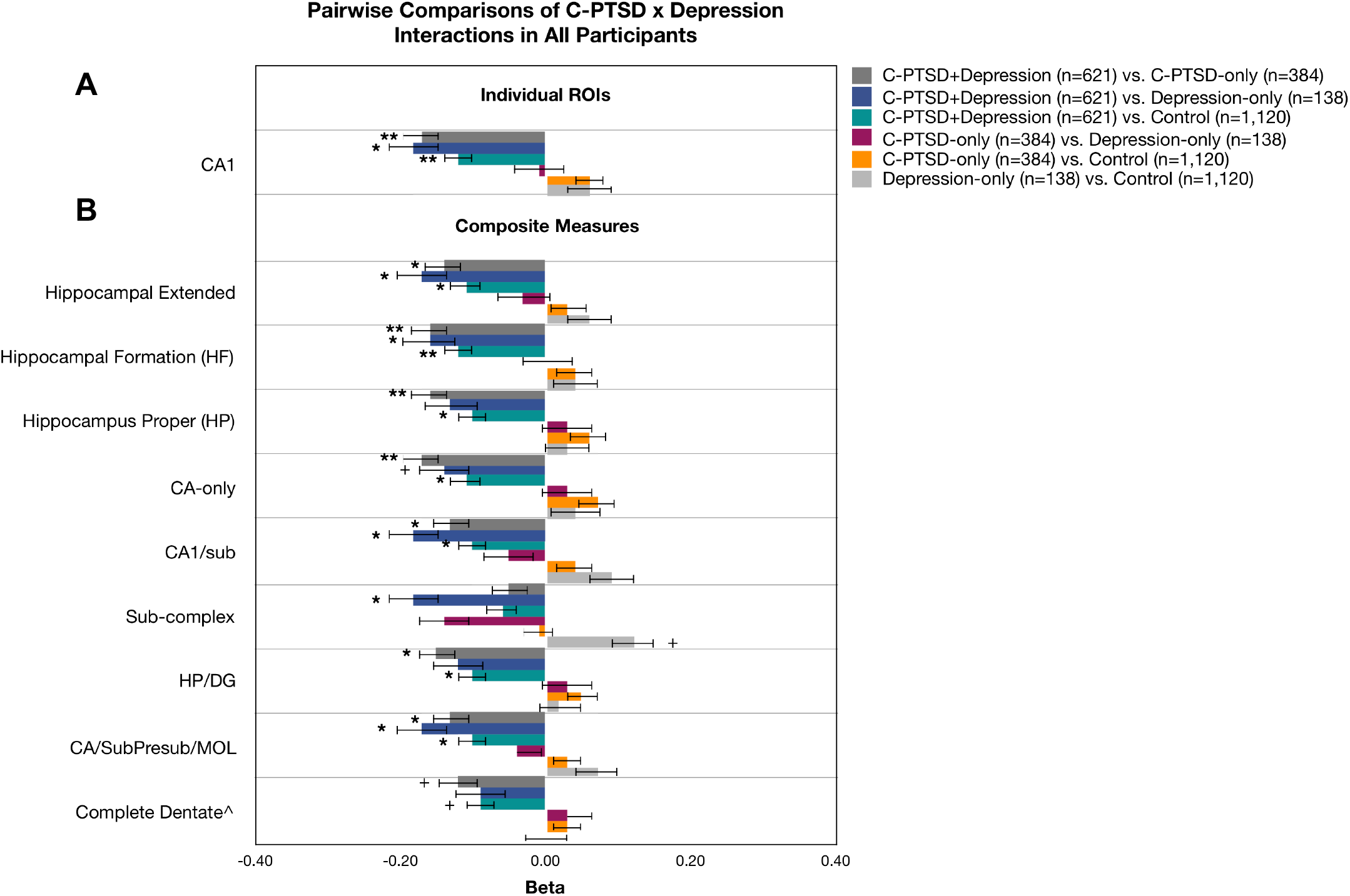
Effect sizes of pairwise comparisons of C-PTSD × Depression interactions. Beta weights are depicted for pairwise comparisons of C-PTSD × Depression interaction terms for analyses of the individual atlas-defined subregions (**A**) and aggregate composite measures (**B**). Separate FDR corrections were applied to each region. We focused on regions that survived FDR correction at the omnibus level; the CA1 is the only individual ROI that survived FDR correction at the omnibus level. Pairwise differences in non-significant ROIs are in **Supplementary Table 19**. ^ The complete dentate was the only composite measure that was not significant at the omnibus level (p=0.128); it is depicted for visual comparison to the other composites. Descriptions of composite measures are in **Supplementary Table 5**. Error bars reflect the standard error. Significance and trend effects were determined using FDR-adjusted p-values (q). + = 0.05<q<0.10, *= 0.01<q<0.05, ** = q<0.01, ***=q<0.001.

### Sensitivity analyses in composite subregions

Negative main effects of depression were significant in all composite regions except the sub-complex. Main effects of C-PTSD trended towards significance in all subfields, with stronger negative effects in composite measures that included the subiculum. C-PTSD*Depression interactions were significant in all composite regions except the complete dentate (**Table 2**). Pairwise comparisons showed a significant negative effect of C-PTSD+Depression in all composite subregions except the sub-complex and complete dentate compared to controls and C-PTSD-only (**Figure 3B**, **Supplementary Table 21**. The strongest effects of depression in the C-PTSD group, and of C-PTSD in the depressed group, were observed in CA-only (ß=−0.17, p=0.001, q=0.006) and CA1/sub (ß=−0.18, p=0.007, q=0.021), respectively. Effects of C-PTSD in the depressed group were only significant in composites that included the subiculum.

**Table 2.** Main Effects and Interactions of Depression and C-PTSD in Composite Measures

| A. Depression (n=801) vs. no depression (n=1459) |  |  |  |  |  |  |
| --- | --- | --- | --- | --- | --- | --- |
|  | Beta | SE | DF | t | p | q |
| CA1/sub | -0.07 | 0.03 | 2255.69 | -1.99 | 0.047* | 0.053<sup>^</sup> |
| Complete dentate | -0.08 | 0.03 | 2250.31 | -2.28 | 0.023* | **0.035\*** |
| CA-only | -0.09 | 0.03 | 2254.08 | -2.75 | 0.006** | **0.018\*** |
| HP | -0.09 | 0.03 | 2252.74 | -2.71 | 0.007** | **0.018\*** |
| HP/DG | -0.09 | 0.03 | 2251.90 | -2.64 | 0.008** | **0.018\*** |
| Sub-complex | -0.02 | 0.03 | 2248.01 | -0.52 | 0.602 | 0.602 |
| CA/SubPresub/MOL | -0.07 | 0.03 | 2255.29 | -2.09 | 0.037* | **0.048\*** |
| HF | -0.09 | 0.03 | 2255.08 | -2.85 | 0.004** | **0.018\*** |
| Hippo extended | -0.08 | 0.03 | 2259.50 | -2.42 | 0.016* | **0.029\*** |
| B. C-PTSD (n=1045) vs. No C-PTSD (n=1361) |  |  |  |  |  |  |
|  | Beta | SE | DF | t | p | q |
| CA1/sub | -0.07 | 0.03 | 2399.31 | -2.21 | 0.027* | 0.061<sup>^</sup> |
| Complete dentate | -0.06 | 0.03 | 2398.68 | -1.76 | 0.078<sup>^</sup> | 0.078<sup>^</sup> |
| CA-only | -0.06 | 0.03 | 2399.86 | -1.88 | 0.060<sup>^</sup> | 0.068<sup>^</sup> |
| HP | -0.06 | 0.03 | 2400.03 | -1.89 | 0.059<sup>^</sup> | 0.068<sup>^</sup> |
| HP/DG | -0.06 | 0.03 | 2399.87 | -1.89 | 0.059<sup>^</sup> | 0.068<sup>^</sup> |
| Sub-complex | -0.07 | 0.03 | 2364.96 | -2.04 | 0.041* | 0.068<sup>^</sup> |
| CA/SubPresub/MOL | -0.07 | 0.03 | 2400.32 | -2.25 | 0.025* | 0.061<sup>^</sup> |
| HF | -0.07 | 0.03 | 2400.48 | -2.29 | 0.022* | 0.061<sup>^</sup> |
| Hippo extended | -0.07 | 0.03 | 2404.85 | -2.36 | 0.018* | 0.061<sup>^</sup> |
| C. C-PTSD x Depression |  |  |  |  |  |  |

|  | Beta | SE | DF | t | p | q |
| --- | --- | --- | --- | --- | --- | --- |
| CA1/sub | -0.22 | 0.08 | 2243.04 | -2.78 | 0.006** | **0.022\*** |
| Complete dentate | -0.12 | 0.08 | 2234.24 | -1.52 | 0.128 | 0.128 |
| CA-only | -0.21 | 0.08 | 2238.29 | -2.59 | 0.010* | **0.022\*** |
| HP | -0.19 | 0.08 | 2236.52 | -2.38 | 0.017* | **0.026\*** |
| HP/DG | -0.18 | 0.08 | 2235.51 | -2.23 | 0.026* | **0.033\*** |
| Sub-complex | -0.17 | 0.08 | 2249.20 | -2.04 | 0.042* | **0.047\*** |
| CA/SubPresub/MOL | -0.20 | 0.08 | 2240.99 | -2.58 | 0.010* | **0.022\*** |
| HF | -0.20 | 0.08 | 2240.68 | -2.52 | 0.012* | **0.022\*** |
| Hippo extended | -0.20 | 0.08 | 2245.07 | -2.54 | 0.011* | **0.022\*** |

*Note.* Q-values depict FDR-corrected significance, values in bold ink passed the FDR threshold. $^{\wedge} = 0.05 < q < 0.10$ , $^* = 0.01 < q < 0.05$ , $^{**} = q < 0.01$ , $^{***} = q < 0.001$ . Descriptions: CA1/sub=CA1 + subiculum, Complete dentate=CA4+DG, HP (hippocampus proper)=CA1+CA2/3+CA4, CA-only= CA1+CA2/3, HP/DG= CA1+CA2/3+DG, Sub-complex=subiculum + presubiculum + parasubiculum, CA/SubPresub/MOL= CA1 + CA2/3 + subiculum + presubiculum + MOL, HF (hippocampal formation) = CA1 + CA2/3 + CA4 + DG + subiculum + tail, hippo extended= CA1 + CA2/3 + CA4 + DG + subiculum + presubiculum + parasubiculum + tail + MOL + HATA + fimbria.

### Effects by sociodemographic status

Detailed results for effects by sex and civilian/military background are in **Supplementary Tables 22-29**. A main effect of depression was significant in the hippocampal tail of civilians, and in the tail and CA1 of females (**Figure 4A, C**). We did not observe an effect of C-PTSD in civilians or females. The effects of depression or C-PTSD in males and military were not significant in any subregion. By contrast, joint models of C-PTSD and depression revealed significant interactions in the CA1 of military, and the CA1, subiculum, fimbria, and MOL of males (**Figure 4B, D**). Interactions were not significant in females or civilians, but subgroups were notably smaller than in males and military, which may explain these results.

**Figure 4A-D.**
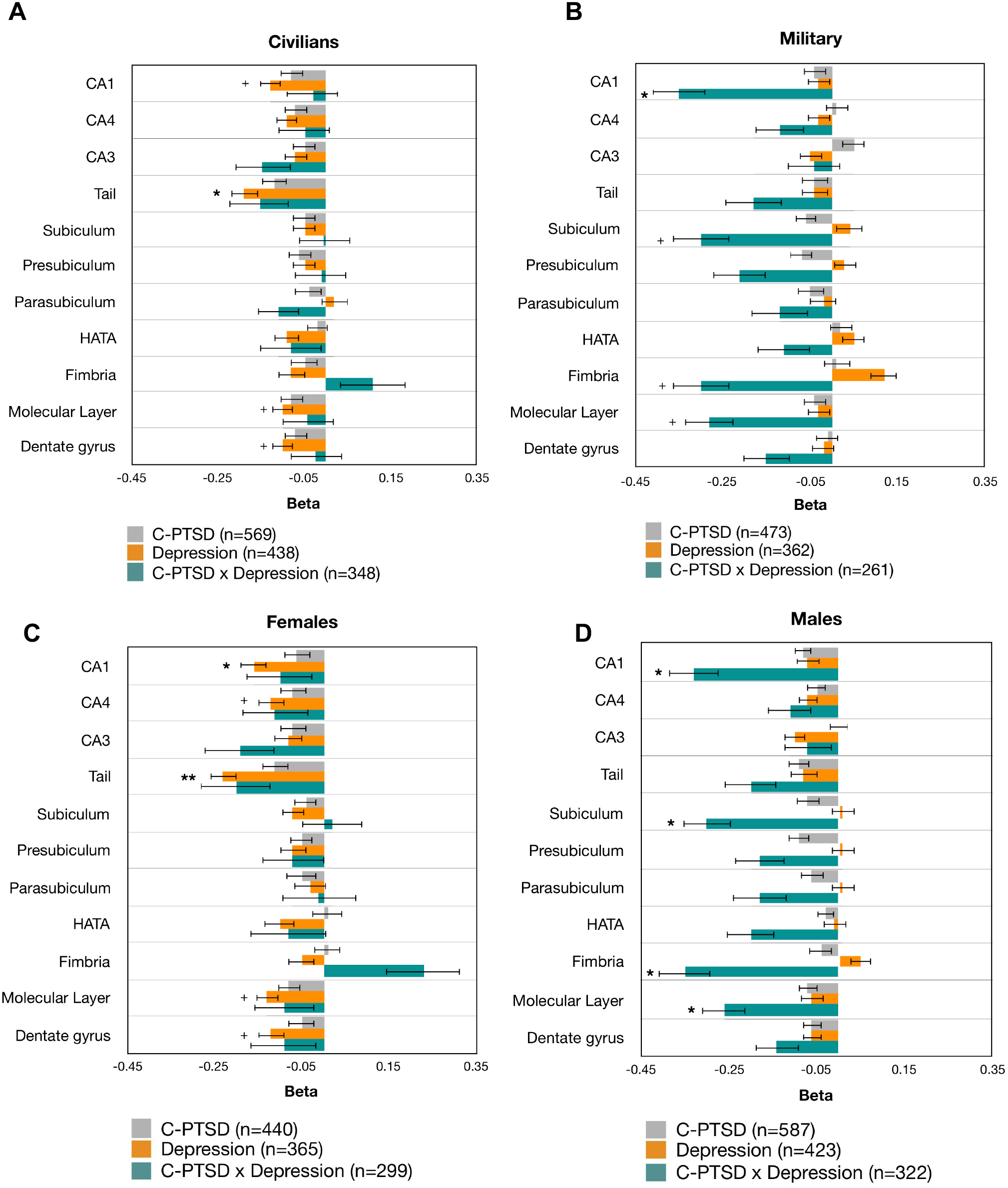
Effect sizes of C-PTSD, depression, and their interaction terms by sex and civilian/military background. Analyses of C-PTSD, depression, and C-PTSD × Depression interactions were computed in separate models. The beta weights of those terms were plotted on the same graph for comparison. Analyses of C-PTSD did not covary for depression, and analyses of depression did not covary for C-PTSD. Sex-related covariates (Sex, Age × Sex, and Age^2^ × Sex) were included in civilian/military analyses. Civilian/military background was included as a covariate in sex-specific analyses. Sample sizes for comparison groups are as follows (**A**) **civilians:** C-PTSD control, n=658; Depression control, n=732; *pairwise reference groups for interactions:* C-PTSD-only, n=203; Depression-only, n=48; control, n=573. (**B**) **military:** C-PTSD control, n=701; Depression control, n=724; *pairwise reference groups for interactions:* C-PTSD-only, n=178; Depression-only, n=90; control, n=523. (**C**) **females:** C-PTSD control, n=486; Depression control, n=513; *pairwise reference groups for interactions:* C-PTSD-only, n=126; Depression-only, n=41; control, n=415. (**D**) **males:** C-PTSD control, n=849; Depression control, n=916; *pairwise reference groups for interactions:* C-PTSD-only, n=258; Depression-only, n=97; control, n=705. Error bars reflect the standard error. Significance and trend effects were determined using FDR-adjusted p-values (q). + = 0.05<q<0.10, *= 0.01<q<0.05, ** = q<0.01, ***=q<0.001

Pairwise comparisons in males and military revealed stronger effects in males than military. Negative effects of depression and C-PTSD were significant in the CA1 of both clinical groups (**Figure 5B**), but the strongest group differences were observed in the fimbria (**Figure 5A-B**). Males with C-PTSD+Depression and C-PTSD-only had significantly smaller fimbriae than individuals with Depression-only (ß=−0.31, b=−0.25), but males with Depression-only had larger fimbriae than controls (ß=0.29). A positive effect of Depression-only also was observed in the subiculum (ß=0.23). Based on the pattern of results by sex and civilian/military background and their strong correlation (phi=0.631), effects found in civilians and military are likely attributable to sex. In males and military, C-PTSD*depression interactions were significant in all composite regions except the complete dentate (**Supplementary Table 30**). In military, the negative effect of depression was strongest in CA-only in the C-PTSD group. In males, the negative effect of C-PTSD was strongest in depressed individuals in composite regions that included the subiculum. A positive effect of Depression-only also was significant in the Sub-complex (**Figure 5C, D**).

**Figure 5A-D.**
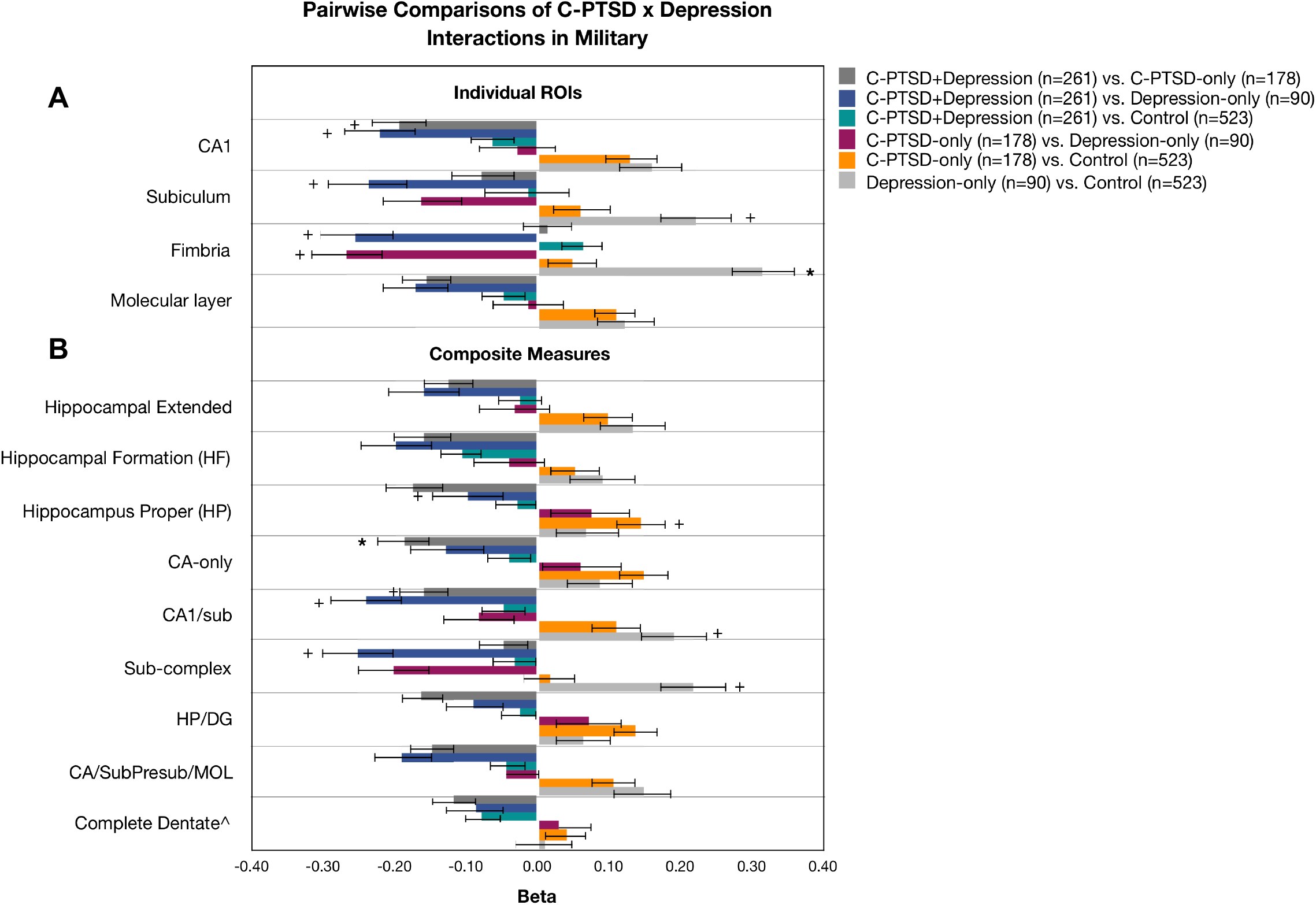
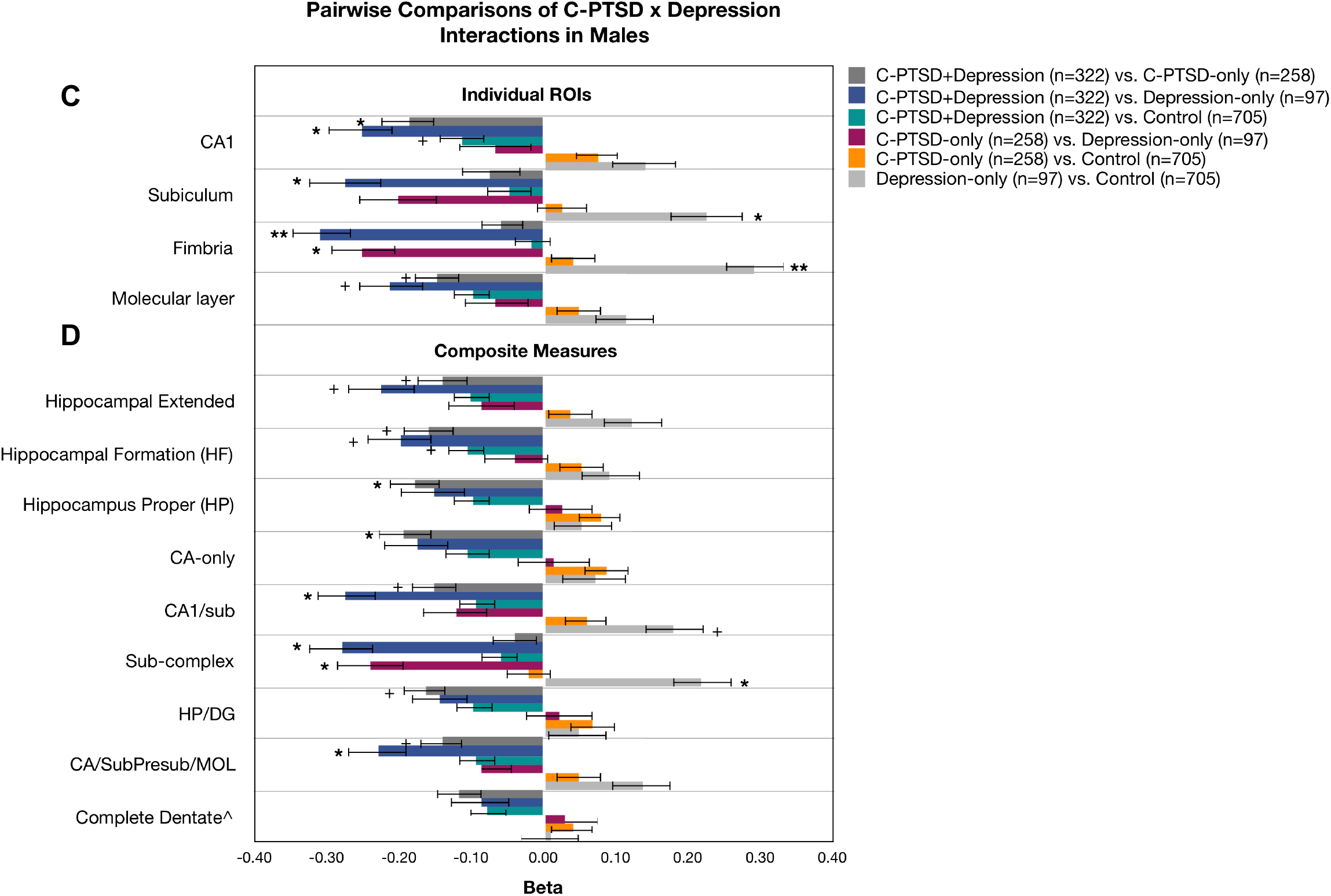
Effect sizes of pairwise comparisons of C-PTSD × Depression Interactions in military and males. Beta weights are depicted for pairwise comparisons of C-PTSD × Depression interaction terms for analyses of the individual atlas-defined subregions (**A**, **C**) and aggregate composite measures (**B**, **D**). Separate FDR corrections were applied to each set of analyses. Interactions were significant in the CA1, subiculum, and fimbria for analyses of males and military. The omnibus interaction in the molecular layer trended towards significance in the military group; interactions in this region were significant in males. ^ The complete dentate was the only composite measure that was not significant at the omnibus level for both analyses (p=0.211, military; p=0.202, males). The omnibus interaction in the HP/DG trended towards significance in the military (p=0.028, q=0.051); both composites are depicted on the graphs for visual comparison. Descriptions of composite measures are in **Supplementary Table 5**. Sub=subiculum, Presub= presubiculum, MOL=molecular layer, DG=dentate gyrus, CA= *cornu ammonis*. Error bars reflect the standard error. Significance and trend effects were determined using FDR-adjusted p-values (q). + = 0.05<q<0.10, *= 0.01<q<0.05, ** = q<0.01, ***=q<0.001.

Ethnicity-unadjusted analyses revealed significant negative main effects of C-PTSD and depression in the CA1, CA4, tail, subiculum, MOL, and DG, and the CA3 specifically in depression. We did not find significant interactions between C-PTSD and depression (**Table 3**).

**Table 3.** Ethnicity-unadjusted analyses

A. C-PTSD (n=1274) vs. no C-PTSD (n=1657)
|  | Beta | SE | DF | t | p | q |
| --- | --- | --- | --- | --- | --- | --- |
| CA1 | -0.08 | 0.03 | 2924.88 | -2.70 | 0.0069** | **0.031\*** |
| CA4 | -0.07 | 0.03 | 2924.85 | -2.33 | 0.020* | **0.044\*** |
| CA3 | -0.04 | 0.03 | 2923.31 | -1.21 | 0.228 | 0.279 |
| Tail | -0.09 | 0.03 | 2875.95 | -2.64 | 0.008** | **0.031\*** |
| Subiculum | -0.07 | 0.03 | 2927.35 | -2.25 | 0.0247* | **0.045\*** |
| Presubiculum | -0.06 | 0.03 | 2868.11 | -1.84 | 0.0655^ | 0.103 |
| Parasubiculum | -0.03 | 0.03 | 2845.31 | -0.89 | 0.372 | 0.409 |
| HATA | -0.04 | 0.03 | 2915.90 | -1.38 | 0.169 | 0.232 |
| Fimbria | -0.01 | 0.03 | 2926.66 | -0.43 | 0.664 | 0.664 |
| Molecular layer | -0.08 | 0.03 | 2925.82 | -2.85 | 0.004** | **0.031\*** |
| Dentate gyrus | -0.07 | 0.03 | 2925.31 | -2.34 | 0.019* | **0.044\*** |

B. Depression (n=967) vs. No Depression (n=1743)
|  | Beta | SE | DF | t | p | q |
| --- | --- | --- | --- | --- | --- | --- |
| CA1 | -0.12 | 0.03 | 2708.37 | -3.85 | 0.0001*** | **0.001\*\*** |
| CA4 | -0.10 | 0.03 | 2702.52 | -3.43 | 0.0006*** | **0.002\*\*** |
| CA3 | -0.10 | 0.03 | 2705.10 | -2.96 | 0.0031** | **0.006\*\*** |
| Tail | -0.13 | 0.04 | 2683.42 | -3.46 | 0.0005*** | **0.002\*\*** |
| Subiculum | -0.04 | 0.03 | 2709.93 | -1.13 | 0.257 | 0.351 |
| Presubiculum | -0.03 | 0.03 | 2696.82 | -1.07 | 0.287 | 0.351 |
| Parasubiculum | -0.02 | 0.04 | 2695.44 | -0.60 | 0.551 | 0.606 |
| HATA | -0.06 | 0.03 | 2706.43 | -1.74 | 0.081^ | 0.128 |
| Fimbria | -0.01 | 0.04 | 2708.51 | -0.36 | 0.717 | 0.717 |
| Molecular layer | -0.10 | 0.03 | 2706.28 | -3.44 | 0.0006*** | **0.002\*\*** |
| Dentate gyrus | -0.10 | 0.03 | 2703.24 | -3.37 | 0.0008*** | **0.002\*\*** |
| C. C-PTSD x Depression |  |  |  |  |  |  |
|  | Beta | SE | DF | t | p | q |
| CA1 | -0.15 | 0.08 | 2692.02 | -2.04 | 0.042* | 0.386 |
| CA4 | -0.06 | 0.07 | 2685.48 | -0.80 | 0.426 | 0.426 |
| CA3 | -0.08 | 0.08 | 2687.06 | -0.94 | 0.350 | 0.426 |
| Tail | -0.10 | 0.09 | 2708.78 | -1.16 | 0.246 | 0.386 |
| Subiculum | -0.10 | 0.08 | 2696.65 | -1.37 | 0.172 | 0.386 |
| Presubiculum | -0.07 | 0.08 | 2704.48 | -0.84 | 0.401 | 0.426 |
| Parasubiculum | -0.12 | 0.09 | 2700.13 | -1.40 | 0.163 | 0.386 |
| HATA | -0.11 | 0.08 | 2690.42 | -1.33 | 0.184 | 0.386 |
| Fimbria | -0.11 | 0.09 | 2696.68 | -1.21 | 0.225 | 0.386 |
| Molecular layer | -0.12 | 0.07 | 2689.21 | -1.60 | 0.110 | 0.386 |
| Dentate gyrus | -0.06 | 0.07 | 2685.72 | -0.88 | 0.376 | 0.426 |

Note. Q-values depict FDR-corrected significance. ^ = 0.05<q<0.10, \* = 0.01<q<0.05, \*\* = q<0.01, \*\*\*=q<0.001. Panel C subgroups: C-PTSD+Depression, n=761, C-PTSD-only, n=442, Depression-only, n=168, healthy control, n=1354

### Post hoc models excluding participants with a history of childhood trauma

We analyzed C-PTSD*Depression interactions in individuals without a history of childhood trauma, as this has shown to alter the directional effects of psychiatric disorders on hippocampal subfield volumes (49, 50). Analyses yielded nominal negative effects of Depression-only compared to controls, particularly in the subiculum (ß=−0.43, p=0.036, q=0.201) and CA1/sub (ß=−0.43, p=0.041, q=0.198). The large reduction in sample size (n=377) limited power to detect other significant outcomes (**Supplementary Table 31**).

## Discussion

Our results reveal distinct influences of C-PTSD and depression on hippocampal subfield volumes across 31 international cohorts. Consistent with previous studies, we observed a significant negative effect of depression on volumes of the CA1 and hippocampal tail. The effect of C-PTSD was not significant, but the combined effect of C-PTSD and depression predicted significantly smaller CA1 volumes compared to healthy controls or either condition alone. Demographic sub-group analyses revealed sex-specific effects in the tail and fimbria, such that smaller volumes in the tail were observed in females with depression, whereas smaller fimbriae were specifically observed in males and were more sensitive to C-PTSD. Finally, sensitivity analyses of composite regions confirmed the importance of the individual atlas-defined CA1 in relation to different clinical phenotypes. Collectively these results reveal nuanced signatures of depression and C-PTSD in hippocampal substructure of this multi-site sample of clinically diverse individuals.

Neuroimaging delineations of PTSD and depression in trauma-exposed samples are controversial. Various lines of evidence suggest that PTSD and depression embody a single construct of general traumatic stress, while others suggest that one disorder is a derivative of the other, and still others conclude these conditions are independent sequelae of trauma exposure (1, 28, 51). Hundreds of papers identify neurotransmitter dysfunction as an etiological mechanism of both conditions, but this literature is complicated by numerous methodological factors including the analysis of candidate biomarkers and neuroendocrine diversity in PTSD and depression subtypes (52, 53). The paragraphs below outline plausible mechanistic interpretations for our key findings and emphasize the need for large-scale longitudinal studies that use careful clinical phenotyping of PTSD and depression and sophisticated multiplex assays that map neuroendocrine function and clinical symptoms to their underlying neural architecture.

### Potential mechanisms underlying morphometry in the CA1

Our results extend prior work showing smaller volumes in the CA1 among individuals with C-PTSD and depression (36, 43, 44), and suggest that CA1 integrity is more vulnerable to the effects of depression than PTSD. CA1 volume differences may reflect stress-induced alterations in negative feedback projections from the hippocampus to the hypothalamus, which are principally supplied by the CA1 pyramidal cell (7, 54). Excitatory input to the CA1 occurs at different layers; distal apical dendrite tufts receive glutamatergic signals from the entorhinal cortex (ERC) through the perforant pathway (ERC layerIII→CA1), whereas dendrites proximal to the soma receive glutamatergic signals from the amygdala and Schaffer collateral (ERC layerII→DG→CA2/3→CA1) (54, 55). Neurotransmitter firing patterns are influenced by diverse populations of GABAergic interneurons that regulate CA1 output to other brain regions, including the hypothalamus, amygdala, and prefrontal cortex (PFC). The compartmentalized structure of the pyramidal cell allows GABAergic interneurons to modulate action potentials at different layers through a temporospatial conductance matrix (56, 57). This is believed to support novelty detection and threat recognition through the integration of previously encoded memories with ongoing sensory and emotional representations from the ERC and amygdala. Reciprocal connections between the CA1 and amygdala also facilitate the consolidation and expression of emotional memories, and are critical for positive adaptation to stress and fear conditioning (56, 57).

GABAergic interneuron activity in the CA1 changes non-uniformly under stress according to the affected interneuron type. In particular, somatostatin (SOM)-expressing interneurons limit excitation to CA1 dendrites and thus are critical for feed-forward inhibition (57) and contextual fear learning (58). CRH mediates SOM secretion and may perpetuate SOM suppression and dendritic atrophy under chronic stress (59). Low levels of SOM may suppress cross-talk between the CA1 and PFC, leading to impaired pattern separation and increased negative-emotion rumination. This idea is consistent with resting-state fMRI work showing suppressed synchronization between the CA1 and superior frontal gyrus following acute social stress (60), and additional work showing that repetitive and anticipatory ruminations moderate the relationship between PTSD and depression (61).

### Alignment to neuroendocrine studies of PTSD and depression

Neuroendocrine studies challenge the hypothesis that hippocampal reduction is a consequence of PTSD, and suggest that smaller hippocampal volume is a pre-existing vulnerability factor (28). The dexamethasone (DEX) suppression test is a common measure of HPA axis inhibition and GR responsivity in the pituitary (8, 28) that mimics the endogenous cortisol response to stress. Paradoxically, individuals with PTSD often show cortisol hyper-suppression in response to low-dose DEX compared to healthy controls (28, 62). Cortisol hyper-suppression is a putative marker of enhanced GR sensitivity that may precede PTSD development (13, 16). Accordingly, twin studies suggest that certain gene-environment interactions may predispose for smaller hippocampal volume and subsequent PTSD risk (63). This latter observation seems to generalize to cumulative stressful, yet non-traumatizing events that interact with genetic factors (64). Childhood trauma has been postulated to alter the developmental programming of the HPA axis, which may leave an “endocrine scar” of low cortisol that primes hippocampal atrophy in adulthood and corresponding risk for mental illness.

History of childhood trauma may explain the bias for larger volumes in several individual ROIs and composite subregions of C-PTSD-only and Depression-only groups compared to controls (**Figures 3, 5**). Larger hippocampal volumes were previously reported in adolescents with PTSD relative to trauma-exposed controls (65), and recent studies revealed subfield hypertrophy in clinical groups with a history of childhood trauma (49, 50). Specifically, Mikolas et al. (49) revealed larger volumes in the CA1, CA3, and MOL of MDD patients with a history of childhood trauma (vs. MDD without childhood trauma) and genetic risk for PTSD (e.g., T allele of *FKBP5*). Janiri et al. (50) revealed larger volumes in the CA1, subiculum, and presubiculum in adults with bipolar disorder (BD) and a history of childhood trauma relative to unexposed BD patients. Our sample was underpowered to test interactions between childhood trauma, C-PTSD, and depression, but a post hoc analysis in individuals without childhood trauma revealed trends for smaller volumes in the C-PTSD-only and Depression-only groups, suggesting a specific role of childhood trauma in the presentation of substructural hippocampal volumetry across psychiatric conditions.

In contrast to PTSD, classical neuroendocrine models of depression show higher levels of awakening cortisol and cortisol hypo-suppression in response to DEX compared to healthy controls (28). However, divergent cortisol patterns have been reported among individuals with melancholic versus atypical depression. While melancholic depression is associated with the classical hypercortisol depression phenotype, atypical depression is associated with suppressive cortisol changes compared to controls, similar to PTSD (66, 67). Interestingly, atypical depression is more common in individuals with a history of trauma exposure and comorbid PTSD (68, 69), and has greater familial specificity (70). Information on depression subtypes were not available in our study, but it is possible that the exacerbated effect of C-PTSD+Depression on lower CA1 volume marks a specific interaction with the atypical depression subtype.

### Localization of subfield results by participant demographics

Sex-specific analyses showed a distinct patterns of subfield volumetry between males and females in the tail and fimbria. Few neuroimaging studies have investigated sex differences in hippocampal subfield volumes in clinical populations, as sex is typically modeled as a nuisance covariate. However, recent work revealed the hippocampal tail as a sexually dimorphic structure (71), with smaller volumes in females that emerge during prepubertal life (72). Thus, smaller volumes in the tail of females may mark a sex-specific vulnerability to depression and explain the strong female-specific effect of depression in our study. Conversely, the fimbria was uniquely affected by C-PTSD+Depression in males. A history of childhood physical neglect has been associated with smaller fimbriae of adults with PTSD and social anxiety disorder, though this was not specific to males (40). Further, Teicher et al. (73) showed that early physical neglect in males is the most significant predictor of hippocampal volume in young adults. However, little is known about the role of the fimbria in clinical and non-clinical samples, as it was not included in earlier hippocampal segmentation protocols. Although it is visible on MRI at 1mm resolution, its image intensity shifts towards that of gray matter with advanced age and is susceptible to partial volume effects (46). However, the FS6 atlas incorporated subject-specific hyperparameters to correct partial voluming and would not explain sex differences in this region.

Finally, the significant main effects of C-PTSD and non-significant C-PTSD*Depression interactions in our ethnicity-unadjusted analyses emphasize the importance of including statistical adjustments for ethnicity in diverse samples. Genetic ancestry data were not available so we cannot infer whether our ethnicity covariates capture variance related to genetics or biopsychosocial characteristics, which should be examined in future work.

### Study limitations

Limitations include the absence of cross-site standardization of clinical raters, scanner operating system, and *a priori* harmonization of MRI parameters. However, FS segmentations of T1-weighted scans acquired from multiple scanners are largely immune to this heterogeneity (74). The uneven availability of covariates across cohorts also precluded analysis of potentially important contributors (e.g., PTSD/depression duration, trauma chronicity/cumulative trauma load, treatment); the absence of these variables limits interpretation. Finally, although we applied sample-wide exclusion criteria to limit heterogeneity attributable to psychiatric disorders beyond depression and AUD, we cannot account for the potential impact of undiagnosed or remitted psychiatric comorbidities that predated the onset of PTSD (75).

### Conclusions

This study is the most powerful investigation of hippocampal subfield volumes in PTSD and depression to date. Our results align with earlier work suggesting that comorbid PTSD and depression represents a unique biological phenotype, with dominant vulnerability of the hippocampal CA1 and sex effects in the tail and fimbria. Studies using data-driven methods are needed to parse the dimensionality of trauma-related symptoms in connection to biopsychosocial and neuroendocrine factors known to influence hippocampal architecture.

## Acknowledgements

The ENIGMA-PGC PTSD Working Group is supported in part by NIH grant R01 MH111671 (PIs: RAM, MWL, PMT, KJR). The ENIGMA Consortium is supported in part by NIH grant U54 EB020403 from the Big Data to Knowledge (BD2K) Program (PI: PMT). These PIs are also supported by grants R56AG058854, R01MH116147, R01MH111671, and P41 EB015922 (PMT); R01NS08688505, I01CX00178301A1, VISN6 MIRECC (RAM); R01MH10882604, I01BX00419201A1, I01BX00347703 (MWL); R01MH10659503, R21AA02745001A1, R01MH10866504, P50MH11587401A1, R01MH11729202, U01 MH11548403 (KJR). LES is supported in part by the NIMH L30MH114379. NJ is supported by R01MH111671, R01MH117601, R01AG059874, MJFF 14848. ELD was supported by K99NS096116. The Booster Cohort was supported by ZonMw, the Netherlands organization for Health Research and Development (40-00812-98-10041), and by a grant from the Academic Medical Center Research Council (110614) both awarded to MO. GLF is supported by DoD W81XWH-10-1-0925, Center for Brain and Behavior Research Pilot Grant, and South Dakota Governor’s Research Center Grant. Research reported in this publication was supported by the South African Medical Research Council for the “Shared Roots” Flagship Project, Grant No MRC-RFA-IFSP-01-2013/SHARED ROOTS through funding received from the South African National Treasury under its Economic Competitiveness and Support Package. Its contents are solely the responsibility of the authors and do not necessarily represent the official views of the South African Medical Research Council. SS is funded by the Department of Science and Technology and the National Research Foundation. LLvdH is supported by the South African Medical Research Council through its Division of Research Capacity Development under the SAMRC Clinician Researcher (MD, PHD) Scholarship Program funded by the South African National Treasury. The content hereof is the sole responsibility of the authors and do not necessarily represent the official views of the SAMRC or the funders. SJHVR is a NARSAD Young Investigator. BSJ is supported by NARSAD 27040. Part of this study was supported by the NSF Graduate Research Fellowship (DWG), UCI - LBVA Biumvirate Grant and R21MH097196 (TGMVE), the National Center for Research Resources and the National Center for Advancing Translational Sciences, National Institutes of Health, through Grant UL1 TR000153, European Research Council Starting Grant 677697 ("BUNGEE-TOOLS") (JEI), the Dana Foundation (JBN), CIHR & CIMVHR (RL), and the South African Medical Research Council (DJS). It was also supported by the following grant numbers: 01-CX000715 & I01-CX001542 (AS); CX001600 VA CDA (JB); F32MH109274 (LAML); K23MH090366-01 (SEB); K23MH101380 (NF); R01AG050595 (CEF, MJK, WDK), R01AG022381 (WDK); R01MH103291 (KAM); R21MH112956 (MLK); R01MH043454 and T32MH018931 (RJD); 1R21MH102634 (IL); R01MH105535 and 1R21MH102634 (IHR); VA RR&D 1K1RX002325 and 1K2RX002922 (SGD); VA RR&D 1IK2RX000709 (NDD); VA RR&D I01RX000622 and CDMRP W81XWH-08– 2–0038 (SRS); and DoD W81XWH08-2-0159 (MBS); DoD W81XWH-10-1-0925 (GF, RS, JS, VM); R01MH11036404, R01MH11168203S1, R01MH10012205, (TJ); R01MH105355 and R01DA035484 (YN, TDW, XZ); R01MH11700901A1 (JS); U01AA021681-08 (MD); and the VA National Center for PTSD (CLA). Additionally, SCM is supported in part by a BOF 2-4 year project (01J05415), and JKD is supported by the German Research Foundation (DFG) – Project 254170585, and EU Rosalind-Franklin Fellowship Program. TPG and UKH are supported by the Research Council of Norway (223273). CRKC is supported by NIA T32AG058507 and NIMH 5T32MH073526. MLK is additionally supported by the Anonymous Women’s Health Fund, Kasparian Fund, Trauma Scholars Fund, and Barlow Family Fund. TS is supported by the German Research Foundation (DFG; SFB/ TRR 58: C06, C07). RB is supported by NHMRC Program Grant No 1073041. RHP is supported by the NIH, grant numbers 1R01 MH114722, 1R01 MH113560, 1R01 MH113406. Earlier versions of this work were presented at the annual meetings for the Society of Biological Psychiatry (May 2017, San Diego CA, USA; May 2019, Chicago IL, USA), Organization for Human Brain Mapping (June 2017, Vancouver, Canada), International Neuropsychological Society (February 2018, Washington D.C., USA), and Society for Brain Mapping and Therapeutics (March 2019, Los Angeles CA, USA).

## Footnotes

1 The hippocampal extended measurement is equivalent to that of whole hippocampal volume. This summed measurement is different than the measure of whole hippocampal volume in Logue et al. (27), which used the autosegmentation ‘aseg’ output from FreeSurfer 5.3 to calculate whole hippocampal volume.

