## Supplementary Table 1 for "Hippocampal subfield volumes are uniquely affected in PTSD and depression: International analysis of 31 cohorts from the PGC-ENIGMA PTSD Working Group": PTSD_subfields_supplement82119.pdf

### SUPPLEMENTARY MATERIAL

#### METHODS

##### *Participants*

All participants had information about current PTSD status, and more than 90% of the total sample was assessed for depression. Subsets of individuals had detailed clinical and demographic information, allowing us to conduct subsample analyses based on demographics (sex, civilian/military status), PTSD status (remitted vs. current PTSD), and common concurrent risk factors (e.g., childhood trauma, mild TBI, AUD).

##### *Assessment and Harmonization of Clinical Covariates*

**PTSD.** Diagnoses were determined using one of seven instruments (**Supplementary Table 2**), primarily using *Diagnostic Statistical Manual-IV* criteria. Overall, 65% of participants were assessed using the Clinician Administered PTSD Scale (CAPS)(1,2). Current PTSD (C-PTSD) symptom severity indexed with the CAPS was available for 838 patients and 733 controls. Data on lifetime PTSD (L-PTSD) history were available for 70% of the total sample. Our main index of PTSD was C-PTSD, as current diagnoses were available in all cohorts.

**Depression**. Ratings of depressive symptoms were available for 91% of the total sample, using one of nine instruments. To facilitate harmonization and to maximize the sample size with depression data, we created a binary index of clinically relevant depression using published thresholds, which served as our primary outcome measure of depression (**Supplementary Table 3**). Approximately 62% of the C-PTSD group met criteria for clinically relevant depression symptoms using these thresholds, compared to 11% of C-PTSD controls. Approximately 64% of the C-PTSD group met criteria for clinically relevant depression symptoms using these thresholds, compared to 13% of C-PTSD controls. The Beck Depression Inventory-II (BDI)(3) was the most frequently used instrument for depression assessment, and total BDI scores were used as our proxy index of depression severity.

**Childhood trauma.** The Childhood Trauma Questionnaire (CTQ; Bernstein et al., 1993) was administered in 37% of participants to index the latent effects of childhood trauma in adults. The CTQ consists of 28 items that assess physical abuse and neglect, emotional abuse and neglect, and sexual abuse, yielding five subscale scores ranging from 5 (*no exposure*) to 25 (*severe exposure*). We used the following published cut points (Bernstein & Fink, 1998) to identify individuals with clinically relevant maltreatment exposure: physical abuse ≥8, physical neglect ≥8, emotional abuse ≥9, emotional neglect ≥10, and sexual abuse ≥6. Participants were then grouped into one of 3 categories: 1) no exposure (n=461), 2) exposure to one type of trauma (n=212), and 3) exposure to two or more trauma types (polytrauma; n=493). This harmonization strategy was selected based on evidence that polytraumatic stress during childhood is associated with worse health outcomes relative to a single type of exposure (4).

**Alcohol use.** Self-reported alcohol use was recorded for 46% of the total sample, with 16% meeting criteria for an alcohol use disorder (AUD). Alcohol use was defined dichotomously as normal or problematic (abuse and/or dependence) using published thresholds. We used binary outcome measures to be consistent with current nomenclature defined by the DSM-5.

**Mild traumatic brain injury (TBI).** Self-reported history of mild TBI (mTBI) was available for 33% of the total sample. Mild TBI was determined by at least one of the following criteria in accordance with severity guidelines outlined by the Veteran Affairs (VA)/Department of Defense (DoD). Mild TBI was recorded as a binary variable (mTBI, n=482; no mTBI, n=552).

**Psychotropic Medication.** Detailed information related to medication use was limited in the source samples for this mega-analysis, but we attempted to account for self-reported medication use in *post hoc* analyses when data were available (n=1,281). We divided participants into one of two groups (medicated, n=275; not medicated, n=1,006) based on self-reported use of any psychoactive medication (antidepressants, anxiolytics, analgesics, and stimulants).

***Neuroimaging***

Identifying the location of the fissure within the hippocampal mask from the general subcortical segmentation, served as an indicator of reliable co-registration of the subfield atlas to the T1-weighted image. As there was no significant difference between the left and right hippocampal effects of PTSD in our recent meta-analyses and other prior work (5)(6), we averaged subfield volumes from left and right hemispheres to decrease measurement noise and reduce the total number of statistical tests, yielding 11 subfield volumes.

Volumes were averaged across hemispheres yielding 11 regions of interest (ROIs). Structural neuroimaging data were obtained using 3T MRI for the majority of scans (>96%). Although two cohorts (n=129; 67 PTSD, 62 controls) acquired images on a 1.5T scanner, we included them in the study because recent work showed strong compatibility between 1.5T and 3T platforms in all hippocampal subregions defined by FreeSurfer (FS) v.6, except the fissure (7). One of these cohorts (n=45) was not included the majority of study analyses due to lack of ethnicity data. Imaging protocols for each site are provided in **Supplementary Table 4**. Sensitivity analyses were conducted in composite measures of subregions described by Roddy et al. (8). Descriptions of each composite measure are provided in **Supplementary Table 5**.

#### Statistical Approach

Descriptive analyses were computed for the whole sample and by cohort (**Table 1**, **Supplementary Tables 1-4**). Continuous variables were z-transformed (mean=0 and standard deviation set to 1 across all cohorts) to standardize the beta weights for a more direct interpretation of explained variance; continuous interaction terms were mean-centered prior to analysis. We used the lmerTest package in R (9) to compute random-effects regression models for primary and secondary analyses, and Satterthwaite approximations were used to compute significance of all fixed effects (10,11). To account for variability across cohorts, we included random effects for the study sample and location of MRI acquisition - scanner manufacturer was included as a nested variable within scanning site to account for variance that occurred when more than one scanner was used at the same site. Basic covariate selection was determined by comparing Akaike information criterion and log likelihood ratios across random effects models consisting of intracranial volume (ICV) and demographic factors only. We observed optimal fit when the following variables were modeled as fixed effects: age, age$^2$, sex, age*sex, age$^2$*sex, ethnicity, and ICV (**Supplementary Table 6**). Civilian/military background was strongly correlated with sex (phi= 0.631, p<0.001) and thus was not included as a covariate in our main models of PTSD and depression.

***PTSD**.* We first tested whether hippocampal subfields differed significantly between C-PTSD and all controls (trauma-exposed + trauma-unexposed), and then between trauma-exposed controls only. C-PTSD severity was analyzed in all participants with CAPS data (n=936), and then separately in the C-PTSD group (n=626). Individuals with remitted PTSD (R-PTSD, n=157) were excluded from these analyses. Separate analyses examined the main effects of trauma exposure in the control group, and the main effects of L-PTSD (n=1192). Participants were only included in the L-PTSD analysis if the parent study confirmed that controls had never received a PTSD diagnosis. We analyzed the effect of R-PTSD by conducting a separate three-level comparison of "PTSD status", which compared trauma-exposed controls, R-PTSD, and C-PTSD.

We tested the relative importance of PTSD on subfield volumes after separately covarying for childhood trauma, mTBI, AUD, and use of psychotropic medication. To determine whether individual subregions could explain a significant degree of variance in the effects of PTSD beyond what is explained by effects on the whole hippocampus, we re-ran our primary analyses covarying for whole hippocampal volume instead of ICV.

**Demographic Analyses.** We examined the effects of C-PTSD, depression, and C-PTSD*Depression separately in males and females and in military and civilian cohorts. We also compared hippocampal subregions between patients and controls, in both C-PTSD and depression, without adjusting for ethnicity to compare results to those in our ethnicity-adjusted models.

**Meta-Analysis.** To detect any bias due to the utilization of a mega-analysis of heterogeneous groups, we duplicated the analysis for C-PTSD main effects, depression main effects, and C-PTSD*depression interactions using a meta-analytic framework. However, this necessitated a reduction in sample sizes, as many cohorts consisted of all cases or controls and hence could not individually generate a stable effect size estimate. Our lower sample-size bound for inclusion in the main effects meta-analysis was a minimum of 5 each cases and controls, and our lower bound for inclusion in the C-PTSD*Depression interaction meta-analyses 5 subjects with each of the 4 C-PTSD by Depression combinations (C-PTSD+Depression, Depression-only, C-PTSD-only, healthy control). Linear regression models were fit for each cohort to estimate the effects effect of C-PTSD, depression, and C-PTSD*depression. Covariates included ICV, sex, age, age$^2$, sex*age, and sex*age$^2$ interaction terms. Ethnicity and site/scanner also were included as covariates if there were variation on these variables within a cohort. Cohen's D estimates were obtained from using a method described in Nakagawa and Cuthill (12), and incorporated a correction for small sample size (sometimes called Hedge's G) (13). We then combined the Cohen's D estimates across all cohorts using a random-effects meta-analysis calculated by the RMA function from the R metaphor package (14).

#### RESULTS

##### *Meta-analysis*

Results of the meta-analyses can be found in **Supplementary Tables 32-33**. Forest plots of the effects of depression in the CA1 and tail are depicted in **Supplementary Figures 3-6**. Similar to the mega-analyses, a negative effect of depression trended towards significance in the CA1 (ß=-0.12, p=0.013, q=0.03) and tail (ß =-0.12, p=0.015, q=0.083) but the effect of C-PTSD was not trending or significant. Interactions between C-PTSD and depression trended towards significance in the CA1 (ß =-0.15, p=0.007, q=0.077), which explained lower volumes in the combined C-PTSD+Depression group versus controls. These effects likely did not pass the significance thresholds due to the reduced sample size in the meta-analysis.

#### SUPPLEMENTARY MATERIAL

| **ST1. Selection criteria for each independent cohort** | | |
|---|---|---|
| Cohort | Inclusion criteria | Exclusion criteria |
| ADNI DoD | *PTSD*: Subjects must be Veterans of the Vietnam War, 50-90 years of age. Subjects who meet the SCID-I (for DSM-IV-TR) criteria for current/chronic PTSD (identified by records and verified by our telephone assessments). In addition to meeting DSM-IV-TR criteria for current/chronic PTSD, subjects must have a minimum current CAPS score of 50 as determined by telephone assessment. The PTSD symptoms contributing to the PTSD Diagnosis and Current CAPS score must be related to a Vietnam War related trauma. Must live within 150 miles of the closest ADNI clinic in subject's area. *Control:* Subjects must be Veterans of the Vietnam War, 50-90 years of age. Comparable in age, gender, and education with TBI and PTSD groups May be receiving VA disability payments for something other than TBI or PTSD – or no disability at all. Must live within 150 miles of the closest ADNI clinic in subject's area | *PTSD*: Mild Cognitive Impairment/Dementia Documented or self-report history of mild/moderate severe TBI Any history of head trauma associated with injury onset cognitive complaints, or Loss of consciousness for >5minutes. Control: MCI/Dementia Presence of PTSD by SCID-I for DSM-IV-TR criteria, or a CAPS score of >30 (Both current and/or a history of PTSD will be excluded). Documented or self-report history of mild/moderate severe TBI Any history of head trauma associated with injury onset cognitive complaints, or Loss of Consciousness for >5 minutes History of PTSD or current PTSD Exclusionary criteria applied to TBI/PTSD will be applied to controls. ALL: MCI/dementia History of psychosis or bipolar affective disorder; History of alcohol or substance abuse/dependence within the past 5 years (by DSM IV – TR criteria); MRI-related exclusions: aneurysm clips, metal implants that are determined to be unsafe for MRI; and/or claustrophobia; Contraindications for lumbar puncture, PET scan, or other procedures in this study; Any major medical condition must be stable for at least 4 months prior to enrollment. These include but are not limited to clinically significant hepatic, renal, pulmonary, metabolic or endocrine disease, cancer, HIV infection and AIDS, as well as cardiovascular disease. Seizure disorder or any systemic illness affecting brain function during the past 5 years will be exclusionary Clinical evidence of stroke. Have a history of relevant severe drug allergy or hypersensitivity. Subjects with current clinically significant unstable medical comorbidities, as indicated by history or physical exam, that pose a potential safety risk to the subject. |
| Booster (AMC) | All: 18-65 years of age, police officers, eligible for MRI. PTSD: current PTSD diagnosis, with CAPS ≥ 45. Controls: exposure to at least one traumatic event (according to DSM-IV A1 criterion), with CAPS < 15 | All: history of neurological disorders, any severe or chronic systemic disease or unstable medical condition (including endocrinological disorders), use of psychotropic medications. Females: pregnancy or breastfeeding. PTSD: current psychotic disorder, substance-related disorder, severe personality disorder, severe major depressive disorder (MDD) (i.e., involving high suicidal risk and/or psychotic symptoms) or current suicidal risk. Controls: any current Axis-1 disorder and lifetime history of PTSD or MDD |
| Cape Town | Between 18 and 65 years, speak English, Afrikaans or Xhosa | Mental retardation, critical medical condition, current psychotic episode/disorder, contraindications for MRI (e.g., metal objects in body, pace makers) |
| Columbia | <u>PTSD</u>: 1. Males or females between the ages of 18 and 60, 2. Current DSM-IV PTSD, 3. CAPS score equal or greater than 50, 4. Able | <u>PTSD</u>: 1. History of Axis I psychiatric diagnosis (other than as specified), e.g., psychotic disorder, bipolar disorder, tic disorder, or eating disorder. Note that comorbid current major depressive disorder will be allowed in up to one half of the PTSD group. This will enable inclusion of this common comorbidity. 2. Depression which is antecedent to PTSD; |

| | | |
|---|---|---|
| | to give consent, fluent in English or Spanish. Clinical interview<br><br><u>Trauma-exposed control</u>: 1. Males or females between the ages of 18 and 60, 2. Capacity to provide consent, 3. History of trauma exposure that fulfills DSM-IV-TR PTSD criteria A1, 4. Able to give consent, fluent in English or Spanish. | score of > 25 on the Hamilton Rating Scale for Depression (HAM-D-17-item); significant depression and /or depression related impairment that is judged to warrant pharmacotherapy or combined medication and psychotherapy. 3.Individuals at risk for suicide based on history and current mental state. 4. For PTSD, history of substance/alcohol dependence within the past six months, and abuse within past two months History, urine toxicology, 5. Patients who are receiving effective medication for their PTSD, and/or depression. Antipsychotic, antidepressant, or mood stabilizer medications in the last 4 weeks prior to the study (6 weeks for fluoxetine). Standing daily dosing of benzodiazepine class of medication in the 2 weeks prior to the study (as needed use of benzodiazepines is not an exclusion but must be clinically judged to tolerate no benzodiazepines for the 72-hour period before each of the fMRI days). Triptan anti-migraine medications. Other medications that may interfere with fear circuitry and fear memory such as blood-brain-barrier penetrating β-blockers. 7. Pregnancy or plans to become pregnant during the period of the study. 8.Paramagnetic metallic implants or devices contraindicating magnetic resonance imaging or any other non-removable paramagnetic metal in the body. 9. Formal CBT psychotherapy initiated within 3 months of beginning this study. 10. Medical illness that could interfere with assessment of diagnosis, treatment response or biological measures (SCR, fMRI), including organic brain impairment from stroke, CNS tumor, or demyelinating disease; and renal, thyroid, hematologic or hepatic impairment. 11. Current unstable or untreated medical illness, and resting SBP≥140 and DBP≥90 and HR<60 and HR>100. 12. Any condition that would exclude clinical MR exam (e.g. pacemaker, paramagnetic metallic prosthesis, surgical clips, shrapnel, necessity for constant medicinal patch, some tattoos). 13. Significant claustrophobia that would preclude ability to remain calm within the MRI scanner. 14. (Supplement PTSD Participants Only): In the treatment of PTSD: Prior intolerance of paroxetine; prior failed adequate trial of paroxetine; or prior failure of adequate trials of 3 different SSRIs. <u>Trauma-exposed control:</u> 1. Current symptomatic major Axis I psychiatric diagnosis, e.g., major depressive disorder, psychotic disorder, bipolar disorder, obsessive compulsive disorder (OCD), PTSD, panic disorder, agoraphobia, or eating disorder. (Specific Phobia will be permitted) 2. CAPS score greater than 20. 3. Depression score greater than 7 on the Hamilton Rating Scale for Depression (HAM-D-17-item). 4 Lifetime history of substance/alcohol dependence or abuse. 5. Pregnancy or plans to become pregnant during the period of the study. 6. Medical illness that could interfere with assessment of response or biological measures (SCR, fMRI), including organic brain impairment from stroke, CNS tumor, or demyelinating disease; and renal, thyroid, hematologic or hepatic impairment. 7.Paramagnetic metallic implants or devices contraindicating magnetic resonance imaging or any other non-removable paramagnetic metal in the body. 8. Medical illness that could interfere with assessment of diagnosis, treatment response or biological measures (SCR, fMRI), including organic brain impairment from stroke, CNS tumor, or demyelinating disease; and renal, thyroid, hematologic or hepatic impairment. 9. Current unstable or untreated medical illness, and resting SBP>140 and DBP>90 and HR<59 and HR>100. 10. Any condition that would exclude clinical MR exam (e.g. pacemaker, paramagnetic metallic prosthesis, surgical |

| | | |
|---|---|---|
| | | clips, shrapnel, necessity for constant medicinal patch, some tattoos). 11. Significant claustrophobia that would preclude ability to remain calm within the MRI scanner |
| Duke/Durham VA | 18-65, OEF/OIF veterans, fluent in English, free of implanted metal objects or metal shards in eyes, antidepressant, sleep, and anti-anxiety medication permitted | Axis I other than PTSD or MDD, current substance abuse or lifetime substance dependence (other than nicotine), high risk for suicide, claustrophobia, neurological disorders, learning disability or developmental delay, major medical conditions |
| Emory GTP | 18-65 years of age, endorsed at least 1 criterion A trauma, English-speaking | Current psychotic symptoms or bipolar disorder, current substance or alcohol dependence, history of head trauma, taking any psychoactive medication, current illegal drug use (verified with urine drug screen within 24 hours of scan) |
| Ghent | Childhood trauma group: experience(s) of physical, sexual, and/or emotional abuse occurring before 17 years of age as per SLESQ. No childhood trauma group: no experience of childhood trauma and no experience of abuse-related trauma (e.g. emotional abuse, physical/sexual assault, etc.) later in life. For both: MRI compatibility (i.e., no pregnancy or metal implants), fluency in Dutch, normal or corrected-to-normal vision, and being 18-60 years of age | Male sex, History of severe head trauma or severe neurological condition |
| Groningen (Charité Berlin) | 20-60 years, women with current PTSD diagnosis determined via the CAPS-4. civilian, no Trauma controls, no healthy controls, Clinical interviews conducted: CAPS, SCID-I, section for personality disorders of SCID-II, SCID-D, sufficient proficiency in German; MRI compatible | Contraindication for MRI, neurologic disorders, history of substance abuse or dependence within the last 6 months; history of head injury, cerebral incidental findings verified by a neuroradiologist after the MR scan; intake of benzodiazepines, tricyclic antidepressants, anticonvulsants; primary diagnosis of borderline personality disorder; current Axis I disorder other than PTSD, comorbid depressive disorder, anxiety disorders, eating disorders, and accentuated borderline personality disorder. |
| INTRUST | Study 1: Deployment during recent OEF/OIF/Operation New Dawn (OND) conflict and meeting DSM-IV diagnostic criteria for one or more anxiety or depressive disorders.<br>Study 2: Cognitive complaints and either PTSD and/or TBI<br>Study 3: Veteran or civilian outpatients with a primary diagnosis of PTSD and Clinician Administered PTSD Scale score greater than 49.<br>Study 4: Returning from OEF/OIF theatre for medical reasons, being a patient at WRAMC or | Study 1: Significant cognitive impairment, severe psychopathology (psychosis, bipolar disorder, imminent suicidality, untreated substance dependence), recent change in pharmacological intervention, concurrent psychotherapy for the target complaint, anticipated change in circumstances that would prevent study completion (e.g., deployment).<br>Study 2: Prior adverse reaction to or taking medications that would interfere with study medications, women who were or were planning to become pregnant or lactating, certain medical exclusions, inability to meet English language proficiency, or history of psychotic disorder, bipolar I, alcohol or stimulant use disorder, tic disorder, severe depression or acute suicidality.<br>Study 3: Unstable medical disease or alanine transferase/aspartate transferase levels significant above normal (or abnormal liver function tests), moderate or severe TBI, pregnant or nursing women, history of psychotic, bipolar, or neurodegenerative disease, |

| | | |
|---|---|---|
| | NNMC, diagnosed as TBI positive (with initial score of 13-15 on the Glasgow Coma Scale) or negative using DoD criteria, aged 18-40, and DEERS eligible.<br><br>Study 5: English language literacy to provide content, negative pregnancy test for women, Glasgow Coma Scale score of 15, extension of GCS with 7-point amnesia scale score of 6 or 7, and clinically judged to be at low risk for taking tramadol.<br><br>Study 6: Age 18-60, active duty or veteran recently screened positive for TBI, receiving care at one the designated study clinical sites, capable of giving informed consent, Defense Enrollment Eligibility Reporting System (DEERS) eligible (subset of participants), verified TBI positive (subset), for those with TBI, GCS score of 13-15. | history of substance use disorder, imminent suicidality, current use of psychotropic medications or recent use of investigational medications, current engagement in evidence-based treatment for PTSD, current litigation for traumatic event, unwillingness to abstain from grapefruit products during the trial, or lack of English proficiency.<br><br>Study 4: Non-medical military separation, inability to provide consent (e.g., due to English language proficiency level), history of penetrating head injury, significant neurological conditions, current treatment for illness that could impact brain functioning, history of major psychiatric conditions including drug or alcohol use disorders, IV medication use for pain.<br><br>Study 5: Pregnant or nursing, homeless, active suicidal or homicidal with plans/ intent, history of lifetime opioid dependence or abuse, psychosis, other substance abuse or dependence (except tobacco) in the past 60 days, anorexia nervosa, antisocial personality disorder, or other psychiatric conditions more clinically prominent than PTSD, serious or unstable illness or history of stroke, seizures, or brain tumor, use of non-study medications unless approved by PI (e.g., benzodiazepines), current psychotherapy, reaction to tramadol.<br><br>Study 6: Speech/language deficit of sufficient severity to preclude answering interview questions, unable or unwilling to provide informed consent and Health Insurance Portability and Accountability Act (HIPAA) authorization ("unable" includes cases in which the potential subject cannot read and understand English well enough to provide informed consent), second level in-depth TBI evaluation done prior to SAFE TBI interview (subset), penetrating head injury (subset), record of drug or alcohol abuse or dependence in the past six months as documented in medical chart (subset), Structured Clinical Interview for Diagnostic and Statistical Manual of Mental Disorders -IV (SCID) current or lifetime PTSD diagnosis related to life events that occurred prior to most recent deployment (subset), taking intravenous medications for pain; participation will be delayed until such medication has been discontinued (subset). |
| Mannheim | PTSD: Women aged 18-65 years, PTSD after childhood sexual or physical abuse before the age of 18 years , Sexual or physical assault must be the index trauma, At least 3 criteria of BPD (including criterion 6: affective instability; IPDE), Commitment and possibility to attend weekly therapy sessions for one year; no planned absence for more than 4 weeks in this period. TC: Women aged 18-65 years, Childhood sexual or physical abuse before the age of 18 years. HC: Women aged 18-65 years | PTSD: General exclusion criteria were traumatic brain injuries, current and lifetime schizophrenia or bipolar-I disorder, mental retardation, severe psychopathology or somatic illness that needs to be treated immediately in another setting (e.g., BMI<16), medical conditions making exposure-based treatment impossible, a suicide attempt within the last two months, and substance dependency with no abstinence within two months prior to the study. TC&HC: any current or previous mental disorder, any psychotherapeutic experience or any intake of psychotropic medication lifetime and at the moment. For the current fMRI study, further exclusion criteria were metal implants, pregnancy, left-handedness, and claustrophobia |
| McLean | History of childhood maltreatment; Legal and mental competency of the patient; Female; All ethnic backgrounds; Age between 18 and 60; | Male; Under 18 or over 60; Delirium secondary to medical illness; History of neurological conditions that may cause significant psychiatric symptomatology (e.g., dementia); Any contraindication to MR scans, including claustrophobia, pregnancy, metal implants, etc.; |

| | | |
|---|---|---|
| | Fluent English speakers; Normal or Corrected Vision | Current alcohol or substance use disorder (within the last month); A history of schizophrenia or other psychotic disorder; History of head injury or loss of consciousness for longer than 5 min (including concussion); Positive pregnancy test |
| Munster | All patients fulfilled the diagnostic criteria for PTSD as primary diagnosis according to the DSM-IV-TR, assessed by the German version of the Structured Clinical Interview for DSM-IV (SCID). Given the focus on interpersonal violence (IPV)-PTSD, the experience of a trauma related to IPV (e.g., rape, sexual or physical abuse) at least once was an inclusion criterion for the patient group. All participants had normal or corrected-to-normal vision and were right-handed as determined by the Edinburgh Handedness Inventory. | No control had lifetime PTSD. MRI contraindications. |
| South Dakota | Military cohort: OEF/OIF<br>SAP cohort: Participants were undergraduate students who were identified as an adult child of an alcoholic parent (ACoA), based on the Children of Alcoholics Screening Test (CAST). A score of 6 or above on the CAST indicated the participant was more than likely the child of an alcoholic/s and raised by this parent/s. | Participants were excluded for current or previous seizure history, contraindications to MRI, or if they exhibited possible psychotic or other psychological symptoms that would make inclusion in the study potentially hazardous to them. |
| Stellenbosch | Adult patients aged ≥18 years with a diagnosis of current PTSD. Able to read and understand the Informed Consent documents and be able to read and write in English or Afrikaans. | Any other major psychiatric disorder (e.g. severe mood and psychotic disorders) or any neurological disorder, and significant head injury, any alcohol or drug use disorder within the past 6 months. Positive pregnancy test. |
| UNSW | 18-60 years | Neurological disorders, under 18; over 60; traumatic brain injury; psychosis |
| U of Sydney | 18-65 years of age, endorsed at least 1 criterion A trauma, English-speaking | History of neurological illness (Huntington's, Parkinson's, dementia, MS, etc.). History of Seizure Disorders, unrelated to head injury. Current diagnosis of schizophrenia spectrum or other psychotic disorders (not related to PTSD). Current diagnosis of bipolar or related disorders (not related to PTSD). Current active homicidal and/or suicidal ideation with intent requiring crisis intervention. Cognitive disorder due to general medical condition other than TBI. Unstable psychological diagnosis that would interfere with accurate data collection, determined by consensus of at least two doctorate-level psychologists. Also, MRI contra-indications for MRI including metallic implants or foreign objects deemed unsafe by the MRI technician (such as but not limited to pace-maker, shrapnel, metallic screws). Surgery in the past 2 months except as approved by the MRI technician (such as but not limited to dental work, colonoscopy). Weight exceeding the capacity of the scanner table. |

| | | |
|---|---|---|
| U of Wisconsin Madison | Age range of 18-50, Capable of giving informed consent, Fluent in English, Exposure to one or more life-threatening war zone trauma events per the Combat Experiences Scale and documented by DD-214, Combat Action Ribbon (Marines), Combat Infantry Badge (Army), or other clear evidence of war zone trauma exposure in Iraq or Afghanistan since 2001, Pharmacological or psychotherapeutic treatment stable for at least 8 weeks prior to beginning of study, with no intent to begin a new course of treatment during the study period | Medical: Weight over 352 pounds (due to constraints of MRI scanner), Women of childbearing potential with positive pregnancy test, looking to conceive during the research timeline, or who are breastfeeding, Metallic implants such as prostheses or aneurysm clip, or electronic implants such as cardiac pacemakers, Neurological or serious medical condition that may contraindicate MRI or that may overlap with physiological substrates of psychiatric conditions, History of seizures or seizure disorder, Moderate or severe traumatic brain injury (over 20 minutes unconscious), Psychiatric/Behavioral: Current active substance dependence or dependence within 3 months (other than nicotine), Meets DSM-IV criteria for bipolar disorder, schizophrenia, schizoaffective disorder, psychotic disorder NOS, delirium, or any DSM-IV cognitive disorder. Substance dependence disorder within 3 months or any current substance dependence, Severe psychiatric instability or severe situational life crises, including evidence of being actively suicidal or homicidal, or any behavior that poses an immediate danger to patient or others. Participants with extensive experience in yoga or meditation, Medications/Therapies: Current use of benzodiazepines and beta-blockers |
| U of Michigan/ VA Ann Arbor | All: 18-55 years of age, combat veterans and civilians, eligible for MRI. PTSD: current PTSD diagnosis, with CAPS ≥ 50. Trauma controls: exposure to at least one traumatic event (according to DSM-IV A1 criterion), with CAPS < 15, Community Controls, CAPS < 15 | All: history of neurological disorders, any severe or chronic disorder; alcohol or drug abuse and/or dependence during course of the study. |
| UMSL | Women with a history of interpersonal violence; Ages 18-55; DSM-IV-TR dx of PTSD (CAPS >45); eligible for MRI. Trauma-exposed controls: women, 18-55, history of interpersonal violence, no lifetime PTSD dx | Male; current history of severe psychiatric illness, including schizophrenia, Bipolar I, drug or alcohol abuse within 3 months prior to screening, active suicidal or homicidal ideation; individuals who are unwilling to terminate other forms of psychotherapy; those having serious, unstable or terminal medical illness that would compromise study participation; mental retardation or another pervasive developmental disorder; a continuing intimate relationship with the perpetrator of the sexual assault; are currently being treated by a pharmacologic agent for the treatment of an Axis I disorder and are unwilling to terminate pharmacological treatment under their physician's care. |
| UMC Utrecht | All: 18-60 years of age, eligible for MRI. PTSD: current PTSD diagnosis, with CAPS ≥ 45, military deployment >4 months. Trauma controls: exposure to at least one traumatic event (according to DSM-IV A1 criterion), with CAPS < 15, no current psychiatric disorder, military deployment >4 months; healthy controls: no current psychiatric disorder according to DSM-IV. | All: history of neurological disorders, any severe or chronic disorder; alcohol or drug abuse and/or dependence during course of the study. |
| VA Minneapolis | veterans of Operation Enduring Freedom and/or Operation Iraqi Freedom, age 22-62, | Participants were excluded from the study if they met criteria for 1) a current substance-induced psychotic disorder or psychotic disorder due to a general medical condition (other |

| | | |
|---|---|---|
| | who had been exposed to combat during their deployment(s). | than TBI), 2) current DSM-IV substance abuse or dependence other than alcohol, caffeine, or nicotine, 3) a moderate or severe traumatic brain injury from either impact or blast, 4) a neurologic condition other than TBI, 5) a current unstable medical condition that would likely affect brain function (e.g., uncontrolled diabetes), or 6) significant imminent risk of suicidal or homicidal behavior. |
| VA West Haven | combat-exposed Veterans with PTSD and 21 age-matched male combat-exposed healthy controls (combat controls; CC). All participants had been deployed on one or more tours to Iraq and/or Afghanistan and reported exposure to combat-related experiences. All participants were 18 to 50 years of age. | Participants were excluded based on moderate and severe TBI, neurological disorder, and MRI contraindications. Participants with PTSD were also excluded on the basis of a diagnosis of current drug/alcohol abuse, recent change in antidepressant medications (stable dose for 4-weeks required). |
| Vietnam Era Twin Study of Aging (VETSA); UCSD | VET Registry members: both twin brothers served in the military between 1965 and 1975 (Vietnam Era). Only male-male twin pairs. At baseline (VETSA 1: 2002-2008): 51-59 years old when recruited, both twins agree to participate; VETSA 2: Follow-up of VETSA 1 (2009-2014; N=1016) or age-matched attrition replacement (N=189). Age 56-66. **MRI sample is a subset participant starting midway through VETSA 1. In addition to above criteria, had to pass standard MRI safety criteria (1.5 T VETSA 1; 3 T VETSA 2). Data provided has been from VETSA 2. | Inclusion: Vietnam Era veteran, age, participation of brother. VETSA was designed as a study of early risk and protective factors for cognitive and brain aging with a focus on genetic and environmental influences. In VETSA 2 the PCL was added to the protocol partly due to the increased interest/findings concerning PTSD/brain/cognition in aging veterans.<br><br>Standard medical, metal, psychological MRI contra-indications for MRI. |
| VUMC Amsterdam | 18-65 | Antisocial personality disorder, DID, recurrent psychoses, current drug abuse or dependence, medication other than stable SSRIs or infrequent benzodiazepine use |
| Western Ontario | Primary diagnosis PTSD | Psychotic disorder, bipolar disorder, traumatic brain injury, narcotic use, active substance use disorder within 3 months of study entry |
| Yale | Participants ranged in age from 21 to 60 and had been deployed on one or more combat tours. | Individuals were excluded from the study if they met any of the following criteria: a diagnosis of bipolar disorder or psychotic disorder, as assessed by the SCID-IV (First, Spitzer, Gibbon, & Williams, 2002); current benzodiazepine use; a history of ADHD, learning disorder, moderate or severe traumatic brain injury (TBI), brain tumor, epilepsy, or a neurological disorder; current inpatient status; or an MRI contraindication. |

**ST2. Descriptive characteristics by cohort**

| Cohort | Age M (SD) | M, F [a] n | White/ Cauc n | Black/ AA n | Multi-racial n | Other Race n | Mil, Civ [b] n | C-PTSD (y, n) | Depression (y, n) | Child Trauma (y, n) | Meds (y, n) | AUD (y, n) | mTBI (y, n) |
|---|---|---|---|---|---|---|---|---|---|---|---|---|---|
| ADNI DoD | 69.1 (4.7) | 102, 0 | 86 | 7 | 4 | 4 | 102, 0 | 50, 52^ | 4, 98 | 0 | -- | -- | 39, 43 |
| BOOSTER | 40.0 (9.9) | 40, 35 | 73 | 0 | 0 | 2 | 0, 75 | 38, 37 | 26, 47 | 10 | -- | 4, 71 | -- |
| Cape Town | 27.9 (6.0) | 0, 65 | 0 | 39 | 26 | 0 | 0, 65 | 6, 59^ | -- | 18, 44 | -- | 12, 48 | -- |
| Columbia | 35.9 (9.9) | 28, 56 | 24 | 24 | 0 | 35 | 0, 83 | 49, 35 | 33, 46 | -- | 0, 84 | 0, 84 | -- |
| Duke | 39.2 (9.9) | 196, 42 | 119 | 106 | 7 | 6 | 238, 0 | 77, 161^ | 73, 163 | 37, 49 | 50, 188 | 29, 209 | -- |
| Emory GTP | 39.6 (12.7) | 4, 142 | 0 | 142 | 1 | 2 | 0, 146 | 43, 103^ | 37, 109 | 68, 78 | 0, 146 | 12, 111 | -- |
| Ghent | 37.1 (12.1) | 0, 67 | -- | -- | -- | -- | 0, 67 | 8, 59 | 17, 49 | -- | 10, 57 | 1, 63 | 1, 66 |
| Groningen | 40.5 (9.7) | 0, 29 | 28 | 0 | 0 | 1 | 0, 29 | 29, 0 | 15, 12 | 0, 27 | -- | -- | -- |
| INTRUST | 35.5 (12.5) | 195, 141 | 254 | 48 | 0 | 17 | 102, 210 | 79, 257 | 66, 269 | 157, 174 | 29, 56 | 56, 277 | 169, 157 |
| Mannheim | 39.9 (12.0) | 0, 27 | 27 | 0 | 0 | 0 | 0, 27 | 27, 0 | 26, 0 | 0, 27 | 15, 12 | -- | -- |
| McLean | 40.0 (12.5) | 0, 55 | 46 | 2 | 4 | 3 | 0, 55 | 40, 15 | 39, 16 | 11, 41 | 36, 16 | 9, 43 | -- |
| Munster | 27.1 (7.1) | 5, 41 | 46 | 0 | 0 | 0 | 0, 46 | 21, 25 | 14, 32 | -- | 5, 41 | -- | -- |
| South Dakota 1 | 31.6 (6.1) | 85, 9 | 86 | 4 | 0 | 4 | 94,0 | 58, 26 | 27, 66 | -- | 9, 83 | 45, 45 | -- |
| South Dakota 2 | 21.2 (2.2) | 13, 15 | 26 | 0 | 2 | 0 | 0, 28 | 20, 8 | 10, 18 | -- | 3, 25 | 14, 14 | -- |
| Stellenbosch | 41.4 (13.0) | 72, 188 | 0 | 0 | 260 | 0 | 0, 260 | 121, 139 | 80, 180 | 113, 147 | -- | -- | -- |
| UCI | 33.5 (7.2) | 28, 0 | 18 | 2 | 0 | 8 | 28, 0 | 13, 15 | 11, 1 | -- | 6, 7 | -- | -- |
| U. Chicago | 31.7 (8.2) | 44, 0 | 41 | 2 | 0 | 1 | 44, 0 | 24, 20 | 18, 6 | -- | 24, 0 | -- | -- |
| U. Michigan | 31.1 (11.0) | 57, 11 | 36 | 4 | 0 | 4 | 30, 22 | 26, 42 | 20, 45 | -- | 2, 42 | -- | -- |
| UMSL | 32.4 (9.7) | 0, 84 | 50 | 25 | 4 | 3 | 0, 84 | 65, 19 | 52, 25 | -- | -- | -- | -- |
| UNSW | 40.2 (12.6) | 101, 65 | 165 | 0 | 0 | 0 | 0, 166 | 82, 84 | 38, 116 | -- | -- | 37, 123 | -- |
| U. Sydney | 35.9 (8.7) | 32, 30 | -- | -- | -- | -- | 0, 62 | 27, 35^ | 6, 0 | -- | -- | -- | -- |
| U. Wisconsin | 30.4 (6.2) | 54, 0 | 51 | 1 | 0 | 1 | 38, 16 | 16, 38 | 39, 15 | -- | 15, 0 | 2, 51 | -- |
| UMC Utrecht | 36.1 (9.8) | 105, 0 | -- | -- | -- | -- | 105, 0 | 54, 51 | 30, 75 | -- | -- | 0, 95 | 10, 81 |
| VA Ann Arbor | 30.7 (7.7) | 61, 0 | 52 | 4 | 2 | 3 | 41, 20 | 41, 20 | 38, 29 | -- | 28, 33 | -- | -- |
| VA Durham | 43.9 (11.9) | 19, 10 | 18 | 11 | 0 | 0 | 29, 0 | 5, 24^ | 7, 22 | -- | -- | 2, 27 | 9, 12 |
| VA Minneapolis | 32.7 (7.9) | 226, 11 | 196 | 0 | 6 | 13 | 237, 0 | 94, 143^ | 78, 100 | -- | -- | -- | 183, 54 |
| VA West Haven | 34.9 (9.6) | 60, 7 | -- | -- | -- | -- | 67, 0 | 36, 31 | 38, 29 | -- | -- | -- | -- |
| VETSA | 61.8 (2.6) | 232, 0 | 206 | 17 | 5 | 4 | 232, 0 | 34, 198 | 52, 165 | -- | 37, 174 | 10, 28 | 71, 139 |

| | | | | | | | | | | | | | |
|---|---|---|---|---|---|---|---|---|---|---|---|---|---|
| VUMC Amsterdam | 36.1 (10.7) | 0, 45 | -- | -- | -- | -- | 0, 45 | 13, 32 | 10, 34 | -- | 6, 39 | -- | -- |
| Western Ontario | 35.9 (12.5) | 33, 78 | -- | -- | -- | -- | 0, 111 | 66, 45 | 49, 54 | 32, 74 | -- | -- | -- |
| Yale | 29.9 (7.8) | 58, 11 | 37 | 2 | 0 | 8 | 70, 0 | 22, 48 | 19, 51 | 25, 44 | -- | -- | -- |
| **TOTAL** | **38.9 (13.9)** | **1850, 1264** | **1685 (65.7%)** | **440 (17.2%)** | **321 (12.5%)** | **119 (4.6%)** | **1457, 1618** | **1284, 1831** | **981, 1854** | **705, 461** | **275, 1006** | **233, 1289** | **482, 552** |

*Note*. Numbers differ from Table 1 as these descriptives reflect the whole sample before removing participants without ethnicity data and those with remitted PTSD. Ethnicity percentages reflect the proportion of each reported race group to the total sample. ^ Remitted PTSD cases constituted the following proportions of controls from the respective cohorts: ADNI DoD, n=8; Cape Town, n=37; Duke, n=36; Durham VA, n= 3; Emory, n=30; Minneapolis VA, n=51; and Sydney, n=4. These participants were only analyzed in models of lifetime and remitted PTSD. For frequencies of childhood trauma, we combined number of types into a single group of "exposed". UCI= University of California Irvine, UMSL= University of Missouri, St. Louis, UNSW, University of New South Wales.

**ST3. Assessments and harmonization for cohorts included in secondary analyses of clinical variables**

| Cohort | PTSD Diagnostic Tool | Depression Tool | Depression Cut Score | CTQ data (y/n) | Alcohol Tool | AUD Cut Score |
|---|---|---|---|---|---|---|
| ADNI DoD | CAPS-4 | GDS | 10 | N | -- | -- |
| AMC Amsterdam | CAPS-4 | HADS-D | 8 | N | AUDIT | 8 |
| Cape Town | MINI | -- | -- | Y | ASSIST | 11 |
| Columbia | CAPS-4 | HAM-D | 10 | N | -- | -- |
| Duke | CAPS-4/CAPS-5 | BDI | 17 | Y | AUDIT | 8 |
| Emory GTP | PSS | BDI | 17 | Y | AUDIT | 8 |
| Groningen | CAPS-4 | BDI | 17 | Y | -- | -- |
| INTRUST | CAPS-4/ MINI / PCL-M/ SCID | PHQ-9 | 10 | Y | AUDIT | 8 |
| Mannheim | SCID | BDI | 17 | Y | -- | -- |
| McLean | CAPS-5 | BDI | 17 | Y | AUDIT | 8 |
| Munster | SCID | BDI | 17 | N | -- | -- |
| South Dakota (STUDY 1, STUDY 2) | PCL-M / PCL-C | CESD, BDI | 16, 17 | N | AUDIT | 8 |
| Stellenbosch | CAPS-5/ MINI | HAM-D | 10 | Y | -- | -- |
| U. California Irvine | CAPS-4 | BDI | 17 | N | -- | -- |
| U. Illinois Chicago | CAPS-4 | BDI | 17 | N | -- | -- |
| U. Michigan | CAPS-4 | BDI/ DASS 21 | 17/ 10 | N | -- | -- |
| U. Missouri St. Louis | CAPS-4 | BDI | 17 | N | -- | -- |
| U. New South Wales | CAPS-4 | HAM-D | 10 | N | MINI | |
| U. Wisconsin | CAPS-4 | BDI | 17 | N | AUDIT | 8 |
| VA Ann Arbor | CAPS-4 | DASS 21 | 10 | N | -- | -- |
| VA Durham | CAPS-4 | BDI | 17 | N | AUDIT | 8 |
| VA Minneapolis (MPLS) | CAPS-4 | SCID | 3* | N | -- | -- |
| VETSA | PCL-C | CESD | 16 | N | AUDIT | 8 |
| Yale | CAPS-4 | BDI | 17 | Y | -- | -- |

*Note.* The table only depicts cohorts with information about race or ethnicity, as these were the cohorts used in analyses of clinical covariates. * For MPLS, depression scores were derived from the sum of the first two criteria on the SCID-IV depression module (low mood % anhedonia; range 2-6). References for each scale are denoted for each of the following measures: PTSD diagnostic tools: CAPS-4 (Clinician Administered PTSD Scale for DSM IV (1)), MINI (Mini-International Neuropsychiatric Interview; (18), CAPS-5 (CAPS for DSM-V; (2)), PSS (Posttraumatic Stress Scale; (20)), PCL-M (PTSD Checklist- Military; (21)), SCID (Structured Clinical Interview for DSM IV; (22)), PCL-C (PTSD Checklist- Civilian; (23)). Depression tools: GDS (Geriatric Depression Scale; (24)). HADS-D (Hospital Anxiety and Depression Scale-Depression; (25)), HAM-D (Hamilton Depression Scale; (26)), BDI (Beck Depression Inventory-II; (3)), PHQ-9 (Patient Health Questionnaire-9; (27)), CESD (Center for Epidemiologic Studies Depression Scale; (28)), DASS 21 (Depression, Anxiety, Stress Scale-21 item short form; (29)). Alcohol tool: AUDIT (Alcohol Use Disorders Identification Test; (30)), ASSIST (Alcohol, Smoking and Substance Involvement Screening Test; (31)).

**ST4. Imaging Acquisition Parameters by Cohort**

| Cohort | Tesla | Scanner | Model | channels in head coil | acquisition sequence | voxel size (mm) | FOV (mm) | acquisition orientation | TR (ms) | TE (ms) | flip angle | slice thickness |
|---|---|---|---|---|---|---|---|---|---|---|---|---|
| ADNI DoD | 3T | GE | Signa HDxt; Discovery MR750; Discovery MR750w | 8;24; 50 | SPGR; FSPGR | 1x1x1.2 | 256x256 | Sagittal | 6.98; 7.34; 7.65 | 2.8; 3.0; 3.1 | 11 | 1.2 |
| BOOSTER | 3T | Philips | Achieva | 32 | FAST MPRage sequence | 1x1x1 | 240x188 | Axial | 8200 | 3.8 | 8 | 1 |
| Cape Town | 3T | Siemen s | Magnetom Allegra | 4 | MPRAGE | 1x1x1.5 | 256x256 | Sagittal | 2000 | 1.5;3.2; 4.8;6.6 | 20 | 1.5 |
| Columbia | 1.5T | GE | 1.5T Excite 3 HD | 8 | MPRAGE | 1x1x1 | 256x256 | Coronal oblique | 7.25 | 3 | 7 | 1 |
| Duke | 3T | GE | SIgna EXCITE | 8 | FSPGR BRAVO | 1x1x1 | 240x240 | Axial | 8148; 7840; 8160 | 3.2; 2.9; 3.2 | 12 | 1 |
| | 3T | GE | MR750 | 8 | FSPGR BRAVO | 0.9375x 0.9375x 1 | 256x256 | Axial | 8160 | 3.2 | 12 | 1 |
| | 3T | GE | 4T LX Nvi | 8 | Spin-echo co-planar | 1x1x1.9 | 240x240 | Axial | 12000 | 5.4 | 20 | 2 |
| Emory GTP | 3T | Siemen s | TimTrio | 12 | MPRAGE | 1x1x1 | 224x256 | Sagittal | 2600 | 3.0 | 8 | 1 |
| Ghent | 3T | Siemen s | TimTrio | 32 | MPRAGE | 1x1x1 | 256x256 | Transversa l | 2250 | 4.2 | 9 | 1 |
| Groningen | 3T | Siemen s | TimTrio | 12 | MPRAGE | 1x1x1 | 256x256 | Sagittal | 1900 | 2.5 | 9 | 1 |
| INTRUST | 3T | GE | Multiple | 8 | SPGR-BRAVO | 1x1x1 | 256x256 | Sagittal | 9150 | 3.7 | 10 | 1 |
| | 3T | Siemen s | Multiple | 12 | MPRAGE | 1x1x1 | 256x256 | Sagittal | 2530 | 3.3 | 7 | 1 |
| | 3T | Philips | Multiple | 8 | TFESENS E | 1x1x1 | 256x256 | Sagittal | 7600 | 3.5 | 7 | 1 |
| Mannheim | 3T | Siemen s | Trio | 32 | SPGR | 1x1x1 | 192x192 | Axial | 2000 | 3.0 | 80 | 3 |

| | | | | | | | | | | | | |
|---|---|---|---|---|---|---|---|---|---|---|---|---|
| McLean | 3T | Siemens | TimTrio | 12 | MEMPRAGE | 1x1x1 | 256x256 | Sagittal | 2530 | 1.6; 3.5; 5.4; 7.2 | 7 | 1 |
| Munster | 3T | Siemens | Magnetom Prisma | HE1-4 | MPRAGE | 1x1x1 | 256x256 | Sagittal | 2130 | 2.3 | 8 | 1 |
| South Dakota | 3T | Siemens | Skyra | 20 | MPRAGE | 1x1x1 | 240x240x180; 256x256x256 | Sagittal, interleaved | 1900 | 2.1 | 9 | .9 |
| Stellenbosch | 3T | Siemens | Allegra | 4 | MPRAGE | 1x1x1 | 256x256 | Sagittal | 2530 | 1.5; 3.2; 4.9; 6.6 | 7 | 1 |
| | 3T | Siemens | Skyra | 32 | MPRAGE | 1x1x1 | 280x280 | Sagittal | 2530 | 1.63; 3.47; 5.31; 7.15 | 7 | 1 |
| UCI | 3T | Philips | Achieva | 8 with SENSE | MPRAGE (T1TFE) | 1x1x1.2 | 240x256x204 | Sagittal | 6788 | 3.2 | 9 | 1.2 |
| U Michigan | 3T | Philips | Achieva 3.0T X-series | 8 with SENSE | SPGR | 1x1x1mm | 256x256 | Sagittal | 9800 | 4.6 | 8 | 1 |
| UMSL | 3T | Siemens | TimTrio | 12 | MPRAGE | 1x1x1 | 256 x 176 | Sagittal | 2400 | 3.1 | 8 | 1 |
| UNSW | 3T | GE | Signa HDx | 8 | 3D SPGR | 1x1x1mm | 256x256 | Sagittal | 8300 | 3.2 | 11 | 1 |
| U Sydney | 3T | GE | MR750 | 8 | 3D | 0.9x.0.9x0.9 | 256x256 | Sagittal | 7256 | 2.8 | 12 | 0.9 |
| U Wisconsin | 3T | GE | X750 Discovery | 8 with ASSET, acceleration factor of 2 | BRAVO | 1x1x1 | 256x256 | Axial | 8160 | 3.2 | 12 | 1 |
| UMC Utrecht | 3T | Philips | Achieva | 8 with SENSE | 3D FAST Echo Field sequence | 0.8x0.7480.748 | 240x240 | Sagittal | 10000 | 3.8 | 8 | 0.748 |
| VA Ann Arbor | 3T | Philips | Achieva X | 8 | 3D TFE | 1x1x1 | 256x256 | Axial | 9800 | 4.6 | 8 | 1 |
| VA Durham | 3T | Philips | Ingenia | 8 | 3D TFE SENSE | .9375x.9375x1 | 240x240 | Axial | 8148 | 3.7 | 8 | 1 |
| VA Minneapolis | 3T | Siemens | TimTrio | 12 | MPRAGE | 1x1x1 | 256x256 | Coronal | 2530 | 3.7 | 7 | 1 |

| | | | | | | | | | | | | |
|---|---|---|---|---|---|---|---|---|---|---|---|---|
| VA West Haven | 3T | Siemens | TimTrio | 12 | MPRAGE | 1x1x1 | 256x256 | Sagittal | 2500 | 2.8 | 7 | 1 |
| VETSA | 3T | Siemens | TimTrio | 32 | MPRAGE | 1x1x1.2 | 256x256 | Sagittal | 2170 | 4.3 | 7 | 1.2 |
| | | GE | Discovery 750 | 8 phased array | 3D FSPGR | 1x1x1.2 | 240x240 | Sagittal | 8084 | 3.16 | 8 | 1.2 |
| VUMC Amsterdam | 1.5T | Siemens | Sonata MR System | 8 | MPRAGE | 1x1x1.5 | 256x256 | Coronal | 2700 | 4.0 | 8 | 1.5 |
| Western Ontario | 3T | Siemens | Magnetom Allegra | 32 phased array | MPRAGE | 1x1x1 | 256x240 | Sagittal | 2300 | 2.9 | 9 | 1 |
| Yale | 3T | Siemens | TimTrio | 32 | MPRAGE | 1x1x1 | 256x256 | Sagittal | 2530 | 2.7 | 7 | 1 |

**ST5. Summary of composite measures derived from the FreeSurfer 6.0 hippocampal subfields extraction protocol**

| Composite label | Subfields added | Descriptions |
|---|---|---|
| CA1/sub* | CA1 + subiculum | Sum of regions that form the CA1-subiculum pathway, the major projection for neurotransmitter output to the entorhinal cortex. The boundary between the CA1 and subiculum is the most inconsistent across imaging protocols (Yushkevich et al. 2015) |
| Complete dentate | CA4 + granule cell layers of the dentate gyrus (DG) | The CA4 is a deep polymorphic hilus of the DG, thus the combination of the CA4 and granule cell layers of the DG comprise the 'Complete dentate'. |
| Hippocampus Proper (HP) | CA1 + CA2/3 + CA4 | CA regions comprise the classical anatomic divisions of the hippocampus. |
| CA-only | CA1 + CA2/3 | Axons from CA3 pyramidal cells form the Schaffer collaterals, which project to the CA1 to facilitate the integration of emotion and memory. |
| HP/DG | CA1 + CA2/3 + CA4 + DG | CA regions collectively with the DG form the major targets of the trisynaptic loop – the major hippocampal relay circuit for synaptic transmission. |
| Sub-complex* | subiculum + presubiculum + parasubiculum | Architectonically distinct subfields that are frequently combined in hippocampal subfield protocols. |
| CA/SubPresub/MOL* | CA1 + CA2/3 + subiculum + presubiculum + molecular layer | Exploratory composite measure encompassing adjacent subregions with synergistic functions. |
| Hippocampal Formation (HF) | CA1 + CA2/3 + CA4 + DG + subiculum + tail | Sum of all subregions that contain components of the HP, DG, subiculum, and tail. Collectively transmit signals through the performant path of the hippocampus. |
| Hippocampal extended | CA1 + CA2/3 + CA4 + DG + subiculum + presubiculum + parasubiculum + molecular layer + tail + HATA + fimbria | Sum of all subfields minus the fissure, which was used for QC purposes in this study. |

*Note.* Composite measures and descriptions are largely based on measures used in Roddy et al. (8). Descriptions can be referenced to Iglesias et al. (15), Yuchkevich et al. (16), Zeineh et al. (17), and Roddy et al. (8). * Denotes composite measures that were not examined in the Roddy et al. paper. CA= *cornu ammonis*, HATA= hippocampal-amygdala-transition-area

**ST6. Multivariate effect of demographic factors on hippocampal subfields**

| | CA1 | | | | CA4 | | | | CA3 | | | |
|---|---|---|---|---|---|---|---|---|---|---|---|---|
| | | df | F | p | | df | F | p | | df | F | p |
| Ethnicity | 3 | 2141.20 | 13.46 | 1.05E-08*** | 3 | 2393.88 | 1.64 | 0.177 | 3 | 2335.08 | 0.28 | 0.842 |
| ICV | 1 | 2552.95 | 699.10 | 2.3E-136*** | 1 | 2543.79 | 758.69 | 2.2E-146*** | 1 | 2548.09 | 455.89 | 3.4E-93*** |
| Age | 1 | 1774.88 | 5.16 | 0.023* | 1 | 2234.52 | 10.21 | 0.001** | 1 | 2316.81 | 6.65 | 0.010* |
| Sex | 1 | 2030.84 | 1.18 | 0.278 | 1 | 2332.63 | 0.05 | 0.829 | 1 | 2306.19 | 0.27 | 0.603 |
| $Age^2$ | 1 | 1455.55 | 8.37 | 0.004** | 1 | 2031.95 | 13.34 | 0.0003*** | 1 | 2142.80 | 7.31 | 0.007** |
| Age*Sex | 1 | 1966.34 | 2.96 | 0.085^ | 1 | 2324.39 | 0.73 | 0.392 | 1 | 2383.11 | 1.37 | 0.242 |
| $Age^2$*Sex | 1 | 1634.96 | 4.04 | 0.045* | 1 | 2139.22 | 1.24 | 0.265 | 1 | 2227.05 | 1.55 | 0.214 |
| | Tail | | | | Subiculum | | | | Presubiculum | | | |
| | | df | F | p | | df | F | p | | df | F | p |
| Ethnicity | 3 | 1174.07 | 2.32 | 0.073^ | 3 | 2030.38 | 5.79 | 0.001** | 3 | 1395.09 | 16.60 | 1.3E-10*** |
| ICV | 1 | 2539.41 | 290.61 | 8.8E-62*** | 1 | 2554.65 | 681.65 | 2.1E-133*** | 1 | 2515.14 | 608.98 | 1.4E-120*** |
| Age | 1 | 1575.86 | 14.45 | 0.0001*** | 1 | 2451.64 | 10.81 | 0.001** | 1 | 2232.90 | 8.17 | 0.004** |
| Sex | 1 | 1309.08 | 0.13 | 0.718 | 1 | 2188.61 | 0.27 | 0.605 | 1 | 1614.09 | 1.51 | 0.220 |
| $Age^2$ | 1 | 1235.90 | 15.77 | 7.6E-05*** | 1 | 2321.67 | 10.51 | 0.001** | 1 | 2013.77 | 12.85 | 0.0003*** |
| Age*Sex | 1 | 1794.96 | 3.71 | 0.054^ | 1 | 2438.70 | 1.16 | 0.282 | 1 | 2321.83 | 1.51 | 0.220 |
| $Age^2$*Sex | 1 | 1418.00 | 4.87 | 0.027* | 1 | 2309.20 | 2.30 | 0.130 | 1 | 2112.65 | 2.24 | 0.135 |
| | Parasubiculum | | | | HATA | | | | Fimbria | | | |
| | | df | F | p | | df | F | p | | DF | F | p |
| Ethnicity | 3 | 1558.81 | 9.35 | 3.98E-06*** | 3 | 2276.36 | 5.19 | 0.001** | 3 | 1803.07 | 7.03 | 0.0001*** |
| ICV | 1 | 2547.52 | 348.32 | 5.8E-73*** | 1 | 2549.89 | 374.00 | 7.3E-78*** | 1 | 2554.93 | 229.04 | 1.3E-49*** |
| Age | 1 | 1961.64 | 0.12 | 0.732 | 1 | 2287.22 | 10.93 | 0.001** | 1 | 530.73 | 1.34 | 0.248 |
| Sex | 1 | 1649.62 | 1.12 | 0.290 | 1 | 2240.92 | 0.41 | 0.524 | 1 | 1379.16 | 0.32 | 0.569 |
| $Age^2$ | 1 | 1634.25 | 0.53 | 0.466 | 1 | 2090.58 | 15.24 | 9.8E-05*** | 1 | 375.73 | 5.22 | 0.023* |
| Age*Sex | 1 | 2087.41 | 0.10 | 0.747 | 1 | 2360.58 | 3.42 | 0.064^ | 1 | 847.97 | 2.26 | 0.133 |
| $Age^2$*Sex | 1 | 1767.57 | 0.01 | 0.909 | 1 | 2182.40 | 4.97 | 0.026* | 1 | 555.89 | 1.67 | 0.196 |
| | Molecular Layer | | | | Dentate Gyrus | | | | | | | |
| | | df | F | p | | df | F | p | | | | |
| Ethnicity | 3 | 2247.47 | 7.63 | 4.5E-05*** | 3 | 2385.44 | 2.72 | 0.0432* | | | | |
| ICV | 1 | 2549.31 | 839.26 | 9.2E-160*** | 1 | 2543.65 | 786.11 | 5.8E-151*** | | | | |
| Age | 1 | 1755.65 | 13.43 | 0.0002*** | 1 | 2157.30 | 12.07 | 0.0005*** | | | | |
| Sex | 1 | 2080.34 | 0.01 | 0.932 | 1 | 2306.15 | 0.00 | 0.969 | | | | |
| $Age^2$ | 1 | 1432.73 | 19.06 | 1.4E-05*** | 1 | 1927.55 | 16.06 | 6.4E-05*** | | | | |
| Age*Sex | 1 | 1946.84 | 2.82 | 0.093^ | 1 | 2266.52 | 1.06 | 0.302 | | | | |
| $Age^2$*Sex | 1 | 1611.55 | 4.22 | 0.040* | 1 | 2052.28 | 1.82 | 0.178 | | | | |

*Note.* p-values were not corrected for FDR since they were used to determine optimal model fit. Results are depicted in all participants (prior to removing individuals with remitted PTSD). ^ = p<0.10, *** = p <0.001, **=p<0.01, * = p<0.05

**ST7. Main Effects of Trauma Exposure and Current PTSD**

| | Beta | SE | df | t.value | p.value | FDR-q |
|---|---|---|---|---|---|---|
| **A. Current PTSD (n=1042) vs. All Controls (n=1359)** | | | | | | |
| CA1 | -0.07 | 0.03 | 2397.22 | -2.25 | 0.024* | 0.088^ |
| CA4 | -0.05 | 0.03 | 2398.57 | -1.71 | 0.087^ | 0.137 |
| CA3 | -0.02 | 0.03 | 2397.83 | -0.48 | 0.631 | 0.650 |
| Tail | -0.09 | 0.04 | 2339.11 | -2.39 | 0.017* | 0.088^ |
| Subiculum | -0.06 | 0.03 | 2394.39 | -1.79 | 0.073^ | 0.134 |
| Presubiculum | -0.07 | 0.03 | 2331.71 | -2.11 | 0.035* | 0.096^ |
| Parasubiculum | -0.04 | 0.04 | 2311.00 | -1.17 | 0.241 | 0.331 |
| HATA | -0.02 | 0.04 | 2389.82 | -0.54 | 0.588 | 0.650 |
| Fimbria | -0.02 | 0.04 | 2384.49 | -0.45 | 0.650 | 0.650 |
| Molecular layer | -0.08 | 0.03 | 2398.86 | -2.41 | 0.016* | 0.088^ |
| Dentate gyrus | -0.06 | 0.03 | 2398.89 | -1.79 | 0.073^ | 0.134 |
| **B. Current PTSD (n=1042) vs. Trauma-Exposed Controls (n=1094)** | | | | | | |
| | Beta | SE | df | t.value | p.value | FDR-q |
| CA1 | -0.06 | 0.03 | 2126.34 | -1.76 | 0.078^ | 0.250 |
| CA4 | -0.03 | 0.03 | 2136.42 | -0.92 | 0.356 | 0.490 |
| CA3 | 0.002 | 0.04 | 2133.85 | 0.04 | 0.966 | 0.966 |
| Tail | -0.07 | 0.04 | 1975.52 | -1.86 | 0.063^ | 0.250 |
| Subiculum | -0.06 | 0.04 | 2100.80 | -1.57 | 0.116 | 0.255 |
| Presubiculum | -0.06 | 0.04 | 2069.08 | -1.69 | 0.091^ | 0.250 |
| Parasubiculum | -0.05 | 0.04 | 2076.76 | -1.15 | 0.250 | 0.458 |
| HATA | -0.01 | 0.04 | 2131.57 | -0.19 | 0.852 | 0.937 |
| Fimbria | -0.01 | 0.04 | 2109.24 | -0.35 | 0.727 | 0.889 |
| Molecular layer | -0.06 | 0.03 | 2133.12 | -1.78 | 0.075^ | 0.250 |
| Dentate gyrus | -0.03 | 0.03 | 2136.34 | -1.03 | 0.302 | 0.475 |

*Note*. Q-values depict FDR-corrected significance, values in bold ink passed the FDR threshold. ^ = 0.05<q<0.10, * = 0.01<q<0.05, ** = 0.001<q<0.01, ***=q<0.001.

**ST8. Effect of C-PTSD severity indexed by total CAPS scores (current)**

| | beta | SE | DF | t | p | q |
|---|---|---|---|---|---|---|
| **A. In the C-PTSD group** | | | | | | |
| CA1 | -0.02 | 0.03 | 617.96 | -0.50 | 0.619 | 0.973 |
| CA4 | 0.003 | 0.03 | 617.28 | 0.12 | 0.907 | 0.975 |
| CA3 | -0.001 | 0.03 | 623.74 | -0.03 | 0.975 | 0.975 |
| Tail | -0.06 | 0.04 | 563.09 | -1.53 | 0.125 | 0.501 |
| Subiculum | -0.03 | 0.03 | 612.50 | -1.02 | 0.309 | 0.680 |
| Presubiculum | -0.05 | 0.03 | 598.69 | -1.49 | 0.137 | 0.501 |
| Parasubiculum | -0.05 | 0.04 | 616.66 | -1.34 | 0.182 | 0.501 |
| HATA | -0.06 | 0.03 | 616.31 | -1.75 | 0.081^ | 0.501 |
| Fimbria | -0.01 | 0.04 | 622.96 | -0.31 | 0.758 | 0.975 |
| Molecular layer | -0.02 | 0.03 | 618.41 | -0.79 | 0.429 | 0.787 |
| Dentate gyrus | -0.003 | 0.03 | 623.78 | -0.10 | 0.918 | 0.975 |
| **B. In the whole sample** | | | | | | |
| | beta | SE | DF | t | p | q |
| CA1 | -0.03 | 0.02 | 1076.66 | -1.46 | 0.145 | 0.451 |
| CA4 | -0.02 | 0.02 | 1073.78 | -0.81 | 0.416 | 0.524 |
| CA3 | -0.02 | 0.03 | 1073.53 | -0.71 | 0.476 | 0.524 |
| Tail | -0.06 | 0.03 | 1075.07 | -2.28 | 0.023* | 0.253 |
| Subiculum | -0.03 | 0.02 | 1077.49 | -1.39 | 0.164 | 0.451 |
| Presubiculum | -0.02 | 0.02 | 1077.92 | -0.91 | 0.365 | 0.524 |
| Parasubiculum | -0.02 | 0.03 | 1077.50 | -0.73 | 0.465 | 0.524 |
| HATA | -0.03 | 0.03 | 1067.18 | -1.19 | 0.234 | 0.515 |
| Fimbria | 0.01 | 0.03 | 1067.76 | 0.28 | 0.780 | 0.780 |
| Molecular layer | -0.04 | 0.02 | 1075.54 | -1.64 | 0.101 | 0.451 |
| Dentate gyrus | -0.02 | 0.02 | 1074.86 | -0.96 | 0.335 | 0.524 |

*Note.* Model coefficients for C-PTSD and CAPS severity scores on subfield volumes, after adjusting for basic covariates. Q-values depict FDR-corrected significance, values in bold ink passed the FDR threshold. ^ = 0.05<q<0.10, * = 0.01<q<0.05, ** = 0.001<q<0.01, ***=q<0.001.

**ST9. Analyses of Trauma Exposure and Lifetime PTSD**

**A. Trauma Exposed (n=1094) vs. Trauma Unexposed (n=256)**

| | Beta | SE | df | t | p | q |
|---|---|---|---|---|---|---|
| CA1 | 0.05 | 0.05 | 1400.71 | 0.98 | 0.326 | 0.717 |
| CA4 | 0.03 | 0.05 | 1419.89 | 0.54 | 0.590 | 0.810 |
| CA3 | 0.01 | 0.05 | 1414.59 | 0.19 | 0.852 | 0.852 |
| Tail | 0.06 | 0.05 | 1197.52 | 1.05 | 0.292 | 0.717 |
| Subiculum | 0.07 | 0.05 | 1375.58 | 1.36 | 0.174 | 0.717 |
| Presubiculum | 0.04 | 0.05 | 1332.84 | 0.74 | 0.462 | 0.810 |
| Parasubiculum | 0.09 | 0.05 | 1372.81 | 1.67 | 0.094^ | 0.717 |
| HATA | 0.01 | 0.05 | 1421.41 | 0.27 | 0.791 | 0.852 |
| Fimbria | -0.02 | 0.06 | 1404.16 | -0.44 | 0.663 | 0.810 |
| Molecular layer | 0.05 | 0.05 | 1411.04 | 1.13 | 0.259 | 0.717 |
| Dentate gyrus | 0.02 | 0.05 | 1420.37 | 0.50 | 0.616 | 0.810 |

**B. Lifetime PTSD (n=1192) vs. Controls (n=614)**

| | Beta | SE | df | t | p | q |
|---|---|---|---|---|---|---|
| CA1 | -0.09 | 0.04 | 1800.94 | -2.32 | 0.020* | 0.073^ |
| CA4 | -0.08 | 0.04 | 1801.13 | -2.01 | 0.045* | 0.103 |
| CA3 | -0.06 | 0.04 | 1802.70 | -1.33 | 0.183 | 0.224 |
| Tail | -0.08 | 0.05 | 1665.80 | -1.80 | 0.072^ | 0.113 |
| Subiculum | -0.10 | 0.04 | 1793.17 | -2.47 | 0.013* | 0.072^ |
| Presubiculum | -0.07 | 0.04 | 1754.28 | -1.66 | 0.098^ | 0.135 |
| Parasubiculum | -0.08 | 0.04 | 1779.91 | -1.85 | 0.064^ | 0.113 |
| HATA | -0.04 | 0.04 | 1804.03 | -1.01 | 0.310 | 0.341 |
| Fimbria | -0.01 | 0.05 | 1796.18 | -0.12 | 0.907 | 0.907 |
| Molecular layer | -0.10 | 0.04 | 1801.54 | -2.63 | 0.009** | 0.072^ |
| Dentate gyrus | -0.08 | 0.04 | 1800.28 | -1.99 | 0.047* | 0.103 |

*Note*. Significance and trend effects were determined using FDR-adjusted p-values (q). ^ = 0.05<q<0.10, * = 0.01<q<0.05, ** = 0.001<q<0.01, ***=q<0.001.

| ST10. Omnibus effect of PTSD status (control vs. remitted vs. current PTSD) | | | | | | |
|---|---|---|---|---|---|---|
| | term | DF | | F | p | q |
| CA1 | PTSD status | 2 | 1615.54 | 2.30 | 0.101 | 0.370 |
| CA4 | PTSD status | 2 | 1640.95 | 1.74 | 0.177 | 0.394 |
| CA3 | PTSD status | 2 | 1632.67 | 0.76 | 0.469 | 0.538 |
| Tail | PTSD status | 2 | 1338.20 | 1.43 | 0.240 | 0.394 |
| Subiculum | PTSD status | 2 | 1596.56 | 5.00 | 0.007** | 0.077^ |
| Presubiculum | PTSD status | 2 | 1543.20 | 0.72 | 0.489 | 0.538 |
| Parasubiculum | PTSD status | 2 | 1576.65 | 1.39 | 0.251 | 0.394 |
| HATA | PTSD status | 2 | 1641.55 | 0.99 | 0.373 | 0.513 |
| Fimbria | PTSD status | 2 | 1605.32 | 0.60 | 0.547 | 0.547 |
| Molecular layer | PTSD status | 2 | 1625.34 | 2.91 | 0.055^ | 0.303 |
| Dentate gyrus | PTSD status | 2 | 1641.27 | 1.66 | 0.190 | 0.394 |

*Note*. PTSD status represents a 3-level grouping variable that compared trauma exposed controls, to remitted PTSD and current PTSD. Significance and trend effects were determined using FDR-adjusted p-values (q). ^ = 0.05<q<0.10, * = 0.01<q<0.05, ** = 0.001<q<0.01, ***=q<0.001.

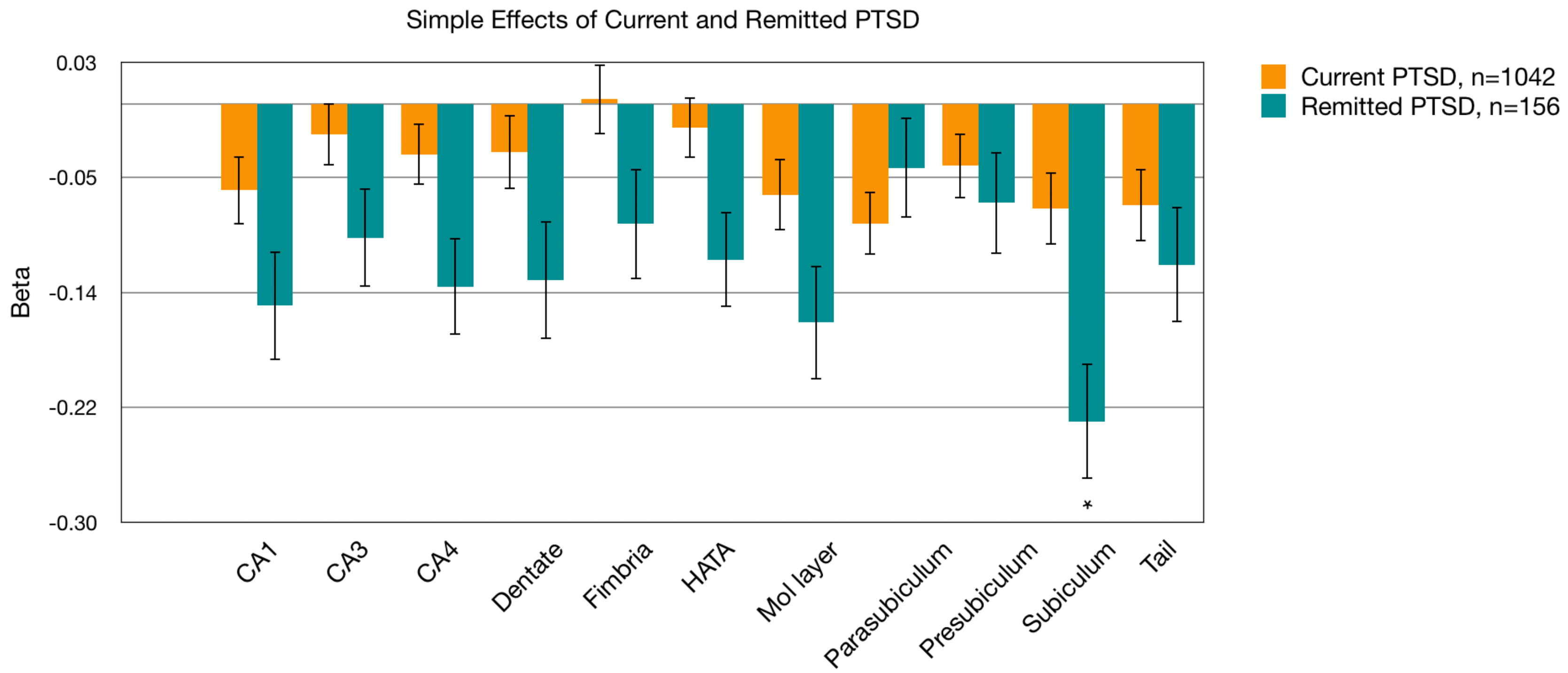


**SF1.** Trauma exposed controls (n=462) served as the reference group and were not output by the model. Group N's were smaller in this analysis based on variable knowledge of lifetime PTSD within the trauma-exposed control group. P values for simple effects are adjusted for multiple comparisons across the lifetime and current diagnostic categories (22 comparisons). ^ = 0.05<q<0.10, * = 0.01<q<0.05, ** = 0.001<q<0.01, ***=q<0.001.

**ST11. Main effects of C-PTSD, controlling for whole hippocampal volume**

| | beta | SE | DF | t | p | q |
|---|---|---|---|---|---|---|
| CA1 | -0.002 | 0.01 | 2318.36 | -0.12 | 0.905 | 0.905 |
| CA4 | 0.01 | 0.02 | 2375.88 | 0.82 | 0.414 | 0.905 |
| CA3 | 0.04 | 0.03 | 2338.61 | 1.52 | 0.129 | 0.905 |
| Tail | -0.03 | 0.03 | 2354.82 | -1.05 | 0.295 | 0.905 |
| Subiculum | 0.01 | 0.02 | 2262.08 | 0.40 | 0.690 | 0.905 |
| Presubiculum | -0.02 | 0.02 | 2358.25 | -0.66 | 0.506 | 0.905 |
| Parasubiculum | -0.004 | 0.04 | 2397.87 | -0.12 | 0.901 | 0.905 |
| HATA | 0.03 | 0.03 | 2376.28 | 1.21 | 0.225 | 0.905 |
| Fimbria | 0.01 | 0.04 | 2381.38 | 0.38 | 0.704 | 0.905 |
| Molecular layer | -0.002 | 0.01 | 2371.17 | -0.24 | 0.808 | 0.905 |
| Dentate gyrus | 0.01 | 0.01 | 2352.74 | 0.84 | 0.403 | 0.905 |

*Note.* Q values depict FDR-adjusted p-values. ICV was removed as a covariate prior to this analysis. Significance and trend effects were determined using FDR-adjusted p-values (q). ^ = 0.05<q<0.10, * = 0.01<q<0.05, ** = q<0.01, ***=q<0.001.

**ST12. Omnibus effects of C-PTSD and medication use after adjusting for covariates**

| | term | Beta | SE | df | t | p | q |
|---|---|---|---|---|---|---|---|
| CA1 | C-PTSD | -0.06 | 0.05 | 1162.99 | -1.19 | 0.233 | 0.393 |
| | PsychotropicMeds | -0.20 | 0.06 | 1136.11 | -3.15 | 0.002** | **0.015*** |
| CA3 | C-PTSD | -0.05 | 0.06 | 1162.85 | -0.95 | 0.343 | 0.503 |
| | PsychotropicMeds | -0.13 | 0.07 | 1153.35 | -1.83 | 0.068^ | 0.150 |
| CA4 | C-PTSD | -0.06 | 0.05 | 1162.51 | -1.15 | 0.249 | 0.393 |
| | PsychotropicMeds | -0.19 | 0.06 | 1158.16 | -2.95 | 0.003** | **0.017*** |
| Dentate gyrus | C-PTSD | -0.06 | 0.05 | 1162.62 | -1.15 | 0.250 | 0.393 |
| | PsychotropicMeds | -0.19 | 0.06 | 1157.77 | -2.91 | 0.004** | **0.018*** |
| Fimbria | C-PTSD | -0.03 | 0.06 | 1150.75 | -0.56 | 0.578 | 0.795 |
| | PsychotropicMeds | 0.03 | 0.07 | 1056.85 | 0.40 | 0.692 | 0.801 |
| HATA | C-PTSD | 0.02 | 0.06 | 1162.58 | 0.44 | 0.659 | 0.801 |
| | PsychotropicMeds | -0.17 | 0.07 | 1149.65 | -2.40 | 0.017* | **0.042*** |
| Molecular layer | C-PTSD | -0.06 | 0.05 | 1163.71 | -1.15 | 0.250 | 0.393 |
| | PsychotropicMeds | -0.22 | 0.06 | 1148.80 | -3.49 | 0.00051*** | **0.006**** |
| Parasubiculum | C-PTSD | -0.01 | 0.06 | 1160.91 | -0.10 | 0.920 | 0.920 |
| | PsychotropicMeds | -0.18 | 0.07 | 1034.11 | -2.43 | 0.015* | **0.041*** |
| Presubiculum | C-PTSD | -0.01 | 0.05 | 1115.35 | -0.20 | 0.841 | 0.911 |
| | PsychotropicMeds | -0.18 | 0.06 | 869.49 | -2.83 | 0.005** | **0.018*** |
| Subiculum | C-PTSD | -0.02 | 0.05 | 1156.82 | -0.49 | 0.621 | 0.801 |
| | PsychotropicMeds | -0.16 | 0.06 | 1090.22 | -2.57 | 0.010* | **0.031*** |
| Tail | C-PTSD | -0.01 | 0.06 | 1120.90 | -0.16 | 0.870 | 0.911 |
| | PsychotropicMeds | -0.31 | 0.07 | 901.28 | -4.30 | 0.00002*** | **0.0004***** |

*Note.* Model coefficients and beta weights for current PTSD when adjusting for self-reported psychotropic medication use. Coefficients and effect size are adjusted for basic covariates. Q-values depict FDR-corrected significance, values in bold ink passed the FDR threshold. ^ = 0.05<q<0.10, * = 0.01<q<0.05, ** = 0.001<q<0.01, ***=q<0.001.

**ST13, SF2 A-B.** Model coefficients and beta weights for childhood trauma (A), and the covaried effect of childhood and C-PTSD (B). Q-values depict FDR-corrected significance, values in bold ink passed the FDR threshold. ^ = 0.05<q<0.10, * = 0.01<q<0.05, ** = 0.001<q<0.01, ***=q<0.001.

| **ST13. A.** Omnibus effects of child trauma after adjusting for basic covariates | | | | | |
|---|---|---|---|---|---|
| Term | DF | F | p | q | ROI |
| Child trauma | 2 | 882.0 | 1.76 | 0.172 | 0.250 | CA1 |
| Child trauma | 2 | 848.8 | 2.52 | 0.081^ | 0.250 | CA4 |
| Child trauma | 2 | 922.8 | 0.55 | 0.575 | 0.575 | CA3 |
| Child trauma | 2 | 874.7 | 3.14 | 0.044* | 0.250 | Tail |
| Child trauma | 2 | 897.9 | 2.31 | 0.100 | 0.250 | Subiculum |
| Child trauma | 2 | 844.6 | 1.71 | 0.182 | 0.250 | Presubiculum |
| Child trauma | 2 | 915.3 | 2.03 | 0.132 | 0.250 | Parasubiculum |
| Child trauma | 2 | 766.4 | 0.80 | 0.448 | 0.548 | HATA |
| Child trauma | 2 | 917.8 | 0.59 | 0.553 | 0.575 | Fimbria |
| Child trauma | 2 | 850.1 | 2.03 | 0.131 | 0.250 | Molecular layer |
| Child trauma | 2 | 878.8 | 1.83 | 0.160 | 0.250 | Dentate gyrus |
| **B.** Covaried effects of PTSD and child trauma after adjusting for basic covariates | | | | | | |
| C-PTSD | 1 | 704.6 | 0.50 | 0.481 | 0.927 | CA1 |
| C-PTSD | 1 | 722.5 | 0.05 | 0.817 | 0.976 | CA4 |
| C-PTSD | 1 | 917.8 | 0.13 | 0.722 | 0.976 | CA3 |
| C-PTSD | 1 | 709.2 | 0.01 | 0.908 | 0.976 | Tail |
| C-PTSD | 1 | 784.1 | 0.02 | 0.893 | 0.976 | Subiculum |
| C-PTSD | 1 | 812.7 | 0.29 | 0.590 | 0.927 | Presubiculum |
| C-PTSD | 1 | 838.4 | 0.00 | 0.967 | 0.976 | Parasubiculum |
| C-PTSD | 1 | 520.0 | 0.08 | 0.771 | 0.976 | HATA |
| C-PTSD | 1 | 866.6 | 0.00 | 0.976 | 0.976 | Fimbria |
| C-PTSD | 1 | 640.7 | 0.53 | 0.468 | 0.927 | Molecular layer |
| C-PTSD | 1 | 787.3 | 0.05 | 0.831 | 0.976 | Dentate gyrus |
| Child trauma | 2 | 916.6 | 1.35 | 0.259 | 0.400 | CA1 |
| Child trauma | 2 | 895.9 | 2.49 | 0.083^ | 0.354 | CA4 |
| Child trauma | 2 | 918.5 | 0.58 | 0.561 | 0.561 | CA3 |
| Child trauma | 2 | 914.2 | 2.86 | 0.058^ | 0.354 | Tail |
| Child trauma | 2 | 916.3 | 2.08 | 0.125 | 0.708 | Subiculum |
| Child trauma | 2 | 898.5 | 1.24 | 0.291 | 0.800 | Presubiculum |
| Child trauma | 2 | 922.2 | 1.99 | 0.137 | 0.708 | Parasubiculum |
| Child trauma | 2 | 881.0 | 0.84 | 0.431 | 0.927 | HATA |
| Child trauma | 2 | 922.7 | 0.59 | 0.554 | 0.927 | Fimbria |
| Child trauma | 2 | 887.9 | 1.62 | 0.198 | 0.726 | Molecular layer |
| Child trauma | 2 | 904.6 | 1.83 | 0.161 | 0.708 | Dentate gyrus |

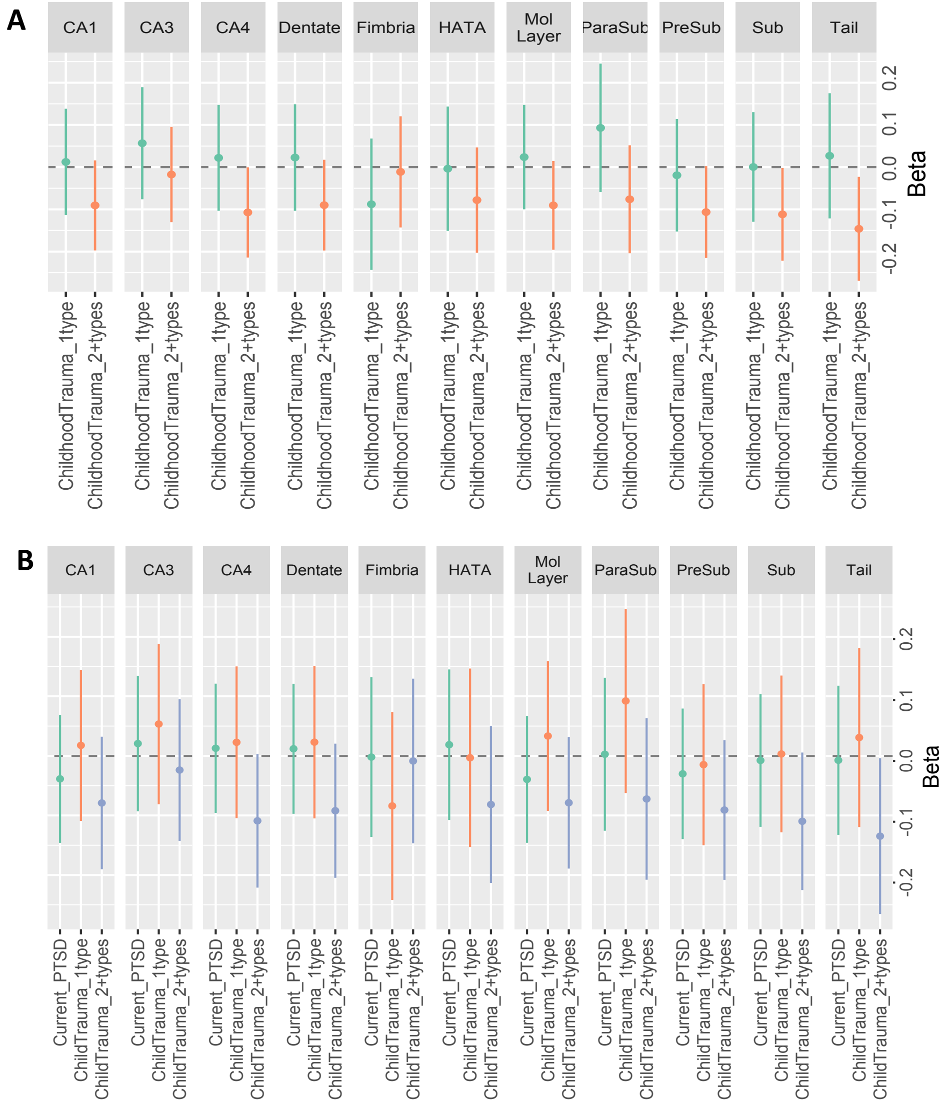

**ST14. Main Effects of C-PTSD (n=694) and Alcohol Use Disorder (AUD, n=209), after adjusting for basic covariates**

| | term | beta | SE | DF | t | p | q |
|---|---|---|---|---|---|---|---|
| CA1 | C-PTSD | -0.04 | 0.04 | 1605.1 | -0.90 | 0.369 | 0.792 |
| | AUD | 0.02 | 0.06 | 1602.4 | 0.38 | 0.707 | 0.951 |
| CA3 | C-PTSD | 0.00 | 0.04 | 1606.3 | 0.02 | 0.986 | 0.991 |
| | AUD | -0.08 | 0.06 | 1601.8 | -1.31 | 0.190 | 0.792 |
| CA4 | C-PTSD | -0.03 | 0.04 | 1608.9 | -0.69 | 0.493 | 0.792 |
| | AUD | 0.00 | 0.06 | 1597.2 | -0.04 | 0.967 | 0.991 |
| Dentate | C-PTSD | -0.03 | 0.04 | 1608.9 | -0.69 | 0.493 | 0.792 |
| | AUD | 0.00 | 0.06 | 1597.2 | 0.01 | 0.991 | 0.991 |
| Fimbria | C-PTSD | -0.04 | 0.05 | 1555.9 | -0.83 | 0.407 | 0.792 |
| | AUD | -0.02 | 0.07 | 1607.7 | -0.28 | 0.776 | 0.951 |
| HATA | C-PTSD | 0.03 | 0.04 | 1593.2 | 0.67 | 0.504 | 0.792 |
| | AUD | 0.01 | 0.07 | 1607.3 | 0.08 | 0.938 | 0.991 |
| Molecular layer | C-PTSD | -0.04 | 0.04 | 1607.8 | -1.10 | 0.271 | 0.792 |
| | AUD | 0.02 | 0.06 | 1599.6 | 0.28 | 0.778 | 0.951 |
| Parasubiculum | C-PTSD | -0.04 | 0.05 | 1518.4 | -0.81 | 0.420 | 0.792 |
| | AUD | -0.13 | 0.07 | 1607.9 | -1.91 | 0.056^ | 0.792 |
| Presubiculum | C-PTSD | -0.04 | 0.04 | 1515.9 | -1.08 | 0.280 | 0.792 |
| | AUD | 0.06 | 0.06 | 1595.3 | 0.90 | 0.366 | 0.792 |
| Subiculum | C-PTSD | -0.04 | 0.04 | 1604.1 | -0.86 | 0.392 | 0.792 |
| | AUD | 0.03 | 0.06 | 1602.2 | 0.43 | 0.665 | 0.951 |
| Tail | C-PTSD | -0.04 | 0.05 | 1555.3 | -0.78 | 0.435 | 0.792 |
| | AUD | 0.06 | 0.07 | 1608.6 | 0.80 | 0.425 | 0.792 |

*Note.* C-PTSD and AUD were corrected separately for FDR. Q-values depict FDR-corrected significance, values in bold ink passed the FDR threshold. C-PTSD control, n=915; AUD control, n=1400. ^ = 0.05<q<0.10, * = 0.01<q<0.05, ** = 0.001<q<0.01, ***=q<0.001.

**ST15. Main Effects of C-PTSD covaried for mild traumatic brain injury (mTBI) after adjusting for basic covariates**

| | term | beta | SE | DF | t | p | q |
|---|---|---|---|---|---|---|---|
| CA1 | C-PTSD | 0.01 | 0.07 | 725.43 | 0.20 | 0.845 | 0.887 |
| | mTBI | -0.03 | 0.06 | 781.10 | -0.45 | 0.655 | 0.947 |
| CA3 | C-PTSD | 0.13 | 0.07 | 629.94 | 1.86 | 0.063^ | 0.576 |
| | mTBI | 0.01 | 0.06 | 764.85 | 0.21 | 0.837 | 0.947 |
| CA4 | C-PTSD | 0.10 | 0.06 | 699.29 | 1.62 | 0.105 | 0.576 |
| | mTBI | 0.00 | 0.05 | 765.48 | 0.06 | 0.949 | 0.947 |
| DG | C-PTSD | 0.07 | 0.06 | 685.61 | 1.10 | 0.273 | 0.676 |
| | mTBI | 0.00 | 0.06 | 765.52 | 0.05 | 0.960 | 0.947 |
| Fimbria | C-PTSD | 0.10 | 0.08 | 712.62 | 1.35 | 0.178 | 0.676 |
| | mTBI | 0.03 | 0.07 | 771.25 | 0.40 | 0.689 | 0.947 |
| HATA | C-PTSD | 0.03 | 0.07 | 706.71 | 0.36 | 0.722 | 0.887 |
| | mTBI | -0.10 | 0.06 | 771.02 | -1.56 | 0.120 | 0.947 |
| Molecular layer | C-PTSD | 0.04 | 0.06 | 746.46 | 0.63 | 0.531 | 0.676 |
| | mTBI | -0.02 | 0.06 | 770.76 | -0.41 | 0.681 | 0.947 |
| Parasubiculum | C-PTSD | 0.07 | 0.08 | 689.88 | 0.80 | 0.426 | 0.882 |
| | mTBI | 0.09 | 0.07 | 760.70 | 1.23 | 0.219 | 0.947 |
| Presubiculum | C-PTSD | 0.01 | 0.07 | 708.90 | 0.14 | 0.887 | 0.653 |
| | mTBI | -0.04 | 0.06 | 757.27 | -0.68 | 0.498 | 0.947 |
| Subiculum | C-PTSD | 0.06 | 0.07 | 717.71 | 0.89 | 0.373 | 0.730 |
| | mTBI | -0.03 | 0.06 | 771.37 | -0.59 | 0.552 | 0.947 |
| Tail | C-PTSD | -0.06 | 0.08 | 708.09 | -0.79 | 0.430 | 0.676 |
| | mTBI | -0.06 | 0.07 | 769.14 | -0.95 | 0.340 | 0.947 |

*Note.* Q-values depict FDR-corrected significance, values in bold ink passed the FDR threshold. ^ = 0.05<q<0.10, * = 0.01<q<0.05, ** = 0.001<q<0.01, ***=q<0.001.

**ST16. Main effects of Depression**

| | **A. Depression (n=800) vs. Control (n=1456) – no C-PTSD Covariate** | | | | | | **B. Depression -Covaried by C-PTSD** | | | | | |
|---|---|---|---|---|---|---|---|---|---|---|---|---|
| | beta | SE | df | t | p | Q | beta | SE | df | t | p | Q |
| CA1 | -0.09 | 0.03 | 2255.90 | -2.73 | 0.006** | **0.033*** | -0.09 | 0.04 | 2247.8 | -2.12 | 0.034* | 0.125 |
| CA4 | -0.08 | 0.03 | 2250.34 | -2.29 | 0.022* | 0.053^ | -0.07 | 0.04 | 2242.2 | -1.90 | 0.057^ | 0.157 |
| CA3 | -0.08 | 0.04 | 2252.08 | -2.08 | 0.037* | 0.068^ | -0.10 | 0.04 | 2243.8 | -2.34 | 0.019* | 0.116 |
| Tail | -0.13 | 0.04 | 2233.62 | -3.22 | 0.001** | **0.011*** | -0.11 | 0.05 | 2243.4 | -2.31 | 0.021* | 0.116 |
| Sub | -0.02 | 0.04 | 2254.86 | -0.52 | 0.601 | 0.741 | 0.01 | 0.04 | 2255.2 | 0.18 | 0.858 | 0.858 |
| Presubiculum | -0.02 | 0.04 | 2238.12 | -0.52 | 0.606 | 0.741 | 0.02 | 0.04 | 2253.3 | 0.57 | 0.571 | 0.636 |
| Parasubiculum | -0.01 | 0.04 | 2243.46 | -0.14 | 0.886 | 0.886 | 0.03 | 0.05 | 2254.6 | 0.56 | 0.578 | 0.636 |
| HATA | -0.04 | 0.04 | 2253.68 | -0.94 | 0.347 | 0.545 | -0.05 | 0.04 | 2246.6 | -1.08 | 0.280 | 0.440 |
| Fimbria | 0.01 | 0.04 | 2252.60 | 0.33 | 0.745 | 0.820 | 0.03 | 0.05 | 2243.1 | 0.57 | 0.571 | 0.636 |
| MOL | -0.08 | 0.03 | 2254.45 | -2.39 | 0.017* | 0.053^ | -0.06 | 0.04 | 2243.7 | -1.60 | 0.110 | 0.202 |
| DG | -0.07 | 0.03 | 2250.45 | -2.25 | 0.024* | 0.053^ | -0.07 | 0.04 | 2241.6 | -1.80 | 0.072^ | 0.158 |

| | **C. C-PTSD –Covaried by depression** | | | | | |
|---|---|---|---|---|---|---|
| | beta | SE | df | t | p | Q |
| CA1 | -0.02 | 0.04 | 2245.43 | -0.43 | 0.668 | 0.816 |
| CA4 | -0.004 | 0.04 | 2254.71 | -0.11 | 0.910 | 0.910 |
| CA3 | 0.05 | 0.04 | 2255.80 | 1.09 | 0.277 | 0.724 |
| Tail | -0.04 | 0.05 | 2172.41 | -0.85 | 0.395 | 0.724 |
| Sub | -0.05 | 0.04 | 2241.79 | -1.21 | 0.225 | 0.724 |
| Presubiculum | -0.08 | 0.04 | 2214.28 | -1.92 | 0.055^ | 0.605 |
| Parasubiculum | -0.06 | 0.05 | 2235.82 | -1.30 | 0.193 | 0.724 |
| HATA | 0.02 | 0.04 | 2255.90 | 0.53 | 0.595 | 0.816 |
| Fimbria | -0.03 | 0.05 | 2210.34 | -0.55 | 0.581 | 0.816 |
| MOL | -0.03 | 0.04 | 2247.97 | -0.86 | 0.389 | 0.724 |
| DG | -0.01 | 0.04 | 2254.34 | -0.24 | 0.808 | 0.889 |

*Note.* Predictor coefficients for depression (panel A) are not adjusted for PTSD. Main effects of PTSD (panel B) were not significant or trending towards significance. Q-values depict FDR-corrected significance, values in bold ink passed the FDR threshold. ^ = 0.05<q<0.10, * = 0.01<q<0.05, ** = 0.001<q<0.01, ***=q<0.001.

| ST17. Effect of depression severity, indexed with the BDI | | | | | | |
|---|---|---|---|---|---|---|
| | | | **A. Individuals with depression** | | | |
| | beta | DF | SE | t | p | q |
| CA1 | -0.04 | 387.94 | 0.04 | -1.12 | 0.262 | 0.702 |
| CA4 | -0.02 | 372.39 | 0.04 | -0.43 | 0.671 | 0.787 |
| CA3 | 0.02 | 376.25 | 0.04 | 0.36 | 0.723 | 0.787 |
| tail | -0.05 | 365.46 | 0.04 | -1.10 | 0.272 | 0.702 |
| Sub | -0.02 | 363.86 | 0.04 | -0.53 | 0.597 | 0.787 |
| PreSub | -0.04 | 366.50 | 0.04 | -1.0 | 0.319 | 0.702 |
| ParaSub | -0.10 | 356.29 | 0.04 | -2.15 | 0.033* | 0.363 |
| HATA | 0.01 | 391.90 | 0.04 | 0.28 | 0.784 | 0.787 |
| Fimbria | 0.02 | 385.59 | 0.04 | 0.48 | 0.631 | 0.787 |
| MOL | -0.04 | 377.36 | 0.04 | -1.08 | 0.280 | 0.702 |
| DG | -0.01 | 377.49 | 0.04 | -0.27 | 0.787 | 0.787 |
| | | | **B. Whole sample with BDI data** | | | |
| | beta | DF | SE | t | p | q |
| CA1 | -0.07 | 680.50 | 0.03 | -2.28 | 0.023* | 0.077^ |
| CA4 | -0.06 | 674.35 | 0.03 | -2.12 | 0.035* | 0.077^ |
| CA3 | -0.05 | 692.93 | 0.03 | -1.41 | 0.159 | 0.219 |
| tail | -0.10 | 623.93 | 0.03 | -3.02 | 0.003** | **0.033*** |
| Sub | -0.03 | 558.06 | 0.03 | -1.04 | 0.300 | 0.367 |
| PreSub | -0.05 | 553.05 | 0.03 | -1.86 | 0.063^ | 0.099^ |
| ParaSub | -0.06 | 441.20 | 0.03 | -1.94 | 0.053^ | 0.097^ |
| HATA | 0.00 | 707.77 | 0.03 | -0.08 | 0.935 | 0.935 |
| Fimbria | 0.03 | 679.35 | 0.03 | 0.92 | 0.357 | 0.393 |
| MOL | -0.07 | 651.51 | 0.03 | -2.51 | 0.012* | 0.066^ |
| DG | -0.06 | 678.19 | 0.03 | -2.11 | 0.035* | 0.077^ |

*Note.* Model coefficients for BDI scores on subfield volumes in the PTSD group (**A**), and in the whole sample (**B**), after adjusting for basic covariates. Q-values depict FDR-corrected significance, values in bold ink passed the FDR threshold. ^ = 0.05<q<0.10, * = 0.01<q<0.05, ** = 0.001<q<0.01, ***=q<0.001.

| ST18. Interaction analyses between C-PTSD and depression | | | | | | | | | | |
|---|---|---|---|---|---|---|---|---|---|---|
| | Beta | SE | DF | t | p | q | C-PTSD-only | Depression-only | C-PTSD +depression | Control |
| CA1 | -0.24 | 0.08 | 2242.95 | -2.92 | 0.004** | **0.044*** | 384 | 138 | 621 | 1120 |
| CA4 | -0.11 | 0.08 | 2234.61 | -1.45 | 0.148 | 0.181 | 384 | 138 | 621 | 1120 |
| CA3 | -0.10 | 0.09 | 2235.93 | -1.20 | 0.232 | 0.232 | 384 | 138 | 621 | 1120 |
| Tail | -0.19 | 0.10 | 2254.18 | -2.01 | 0.045* | 0.124 | 384 | 138 | 621 | 1120 |
| Subiculum | -0.17 | 0.08 | 2247.78 | -2.10 | 0.036* | 0.124 | 384 | 138 | 621 | 1120 |
| Presubiculum | -0.14 | 0.08 | 2253.52 | -1.61 | 0.108 | 0.160 | 384 | 138 | 621 | 1120 |
| Parasubiculum | -0.13 | 0.10 | 2249.24 | -1.35 | 0.176 | 0.194 | 384 | 138 | 621 | 1120 |
| HATA | -0.15 | 0.09 | 2239.51 | -1.64 | 0.102 | 0.160 | 384 | 138 | 621 | 1120 |
| Fimbria | -0.15 | 0.10 | 2245.12 | -1.57 | 0.116 | 0.160 | 384 | 138 | 621 | 1120 |
| Molecular layer | -0.19 | 0.08 | 2239.06 | -2.48 | 0.013* | 0.072^ | 384 | 138 | 621 | 1120 |
| Dentate gyrus | -0.12 | 0.08 | 2234.06 | -1.58 | 0.114 | 0.160 | 384 | 138 | 621 | 1120 |

Note. Q-values depict FDR-corrected significance, values in bold ink passed the FDR threshold. ^ = 0.05<q<0.10, * = 0.01<q<0.05, ** = 0.001<q<0.01, ***=q<0.001.

**ST19. Pairwise comparisons of C-PTSD*Depression interactions**

| ROI | Pairwise Groups | beta | SE | df | t | lowerCI | upperCI | p | Pairwise-q |
|---|---|---|---|---|---|---|---|---|---|
| CA1 | **C-PTSD+Depression - C-PTSD-only** | -0.17 | 0.05 | 2248.39 | -3.46 | -0.27 | -0.08 | 0.001** | **0.006**** |
| | **C-PTSD+Depression – Depression-only** | -0.18 | 0.07 | 2248.52 | -2.63 | -0.32 | -0.05 | 0.009** | **0.018*** |
| | **C-PTSD+Depression - control** | -0.12 | 0.04 | 2249.58 | -2.99 | -0.20 | -0.04 | 0.003** | **0.009**** |
| | C-PTSD-only - Depression-only | -0.01 | 0.07 | 2244.80 | -0.08 | -0.15 | 0.14 | 0.938 | 0.938 |
| | C-PTSD-only - control | 0.06 | 0.05 | 2247.45 | 1.22 | -0.03 | 0.15 | 0.223 | 0.335 |
| | Depression-only - control | 0.06 | 0.06 | 2236.28 | 0.97 | -0.06 | 0.19 | 0.331 | 0.397 |
| CA3 | C-PTSD+Depression - C-PTSD-only | -0.14 | 0.05 | 2249.41 | -2.59 | -0.24 | -0.03 | 0.010* | 0.060^ |
| | C-PTSD+Depression – Depression-only | -0.03 | 0.07 | 2236.89 | -0.35 | -0.17 | 0.12 | 0.725 | 0.725 |
| | C-PTSD+Depression - control | -0.06 | 0.04 | 2254.98 | -1.46 | -0.14 | 0.02 | 0.145 | 0.224 |
| | C-PTSD-only - Depression-only | 0.11 | 0.08 | 2239.98 | 1.44 | -0.04 | 0.27 | 0.149 | 0.224 |
| | C-PTSD-only - control | 0.08 | 0.05 | 2255.51 | 1.56 | -0.02 | 0.18 | 0.119 | 0.224 |
| | Depression-only - control | -0.04 | 0.07 | 2226.63 | -0.52 | -0.17 | 0.10 | 0.605 | 0.725 |
| CA4 | C-PTSD+Depression - C-PTSD-only | -0.12 | 0.05 | 2247.68 | -2.39 | -0.21 | -0.02 | 0.017* | 0.075^ |
| | C-PTSD+Depression – Depression-only | -0.08 | 0.07 | 2236.12 | -1.25 | -0.21 | 0.05 | 0.212 | 0.424 |
| | C-PTSD+Depression - control | -0.09 | 0.04 | 2255.36 | -2.25 | -0.16 | -0.01 | 0.025* | 0.075^ |
| | C-PTSD-only - Depression-only | 0.03 | 0.07 | 2238.89 | 0.48 | -0.11 | 0.18 | 0.632 | 0.758 |
| | C-PTSD-only - control | 0.03 | 0.05 | 2255.10 | 0.68 | -0.06 | 0.12 | 0.493 | 0.740 |
| | Depression-only - control | 0.00 | 0.06 | 2225.52 | -0.05 | -0.13 | 0.12 | 0.961 | 0.961 |
| Dentate | C-PTSD+Depression - C-PTSD-only | -0.12 | 0.05 | 2247.05 | -2.39 | -0.21 | -0.02 | 0.017* | 0.066^ |
| | C-PTSD+Depression – Depression-only | -0.09 | 0.07 | 2235.76 | -1.43 | -0.22 | 0.04 | 0.152 | 0.304 |
| | C-PTSD+Depression - control | -0.09 | 0.04 | 2255.48 | -2.29 | -0.16 | -0.01 | 0.022* | 0.066^ |
| | C-PTSD-only - Depression-only | 0.02 | 0.07 | 2238.06 | 0.31 | -0.12 | 0.16 | 0.758 | 0.905 |
| | C-PTSD-only - control | 0.03 | 0.05 | 2254.87 | 0.65 | -0.06 | 0.12 | 0.517 | 0.776 |
| | Depression-only - control | 0.01 | 0.06 | 2224.76 | 0.12 | -0.11 | 0.13 | 0.905 | 0.905 |
| Fimbria | C-PTSD+Depression - C-PTSD-only | -0.03 | 0.06 | 2235.87 | -0.51 | -0.15 | 0.09 | 0.607 | 0.832 |
| | C-PTSD+Depression – Depression-only | -0.13 | 0.08 | 2252.25 | -1.60 | -0.29 | 0.03 | 0.110 | 0.342 |
| | C-PTSD+Depression - control | -0.01 | 0.05 | 2229.80 | -0.19 | -0.10 | 0.08 | 0.848 | 0.848 |
| | C-PTSD-only - Depression-only | -0.10 | 0.09 | 2236.86 | -1.14 | -0.27 | 0.07 | 0.256 | 0.512 |
| | C-PTSD-only - control | 0.02 | 0.06 | 2210.42 | 0.39 | -0.09 | 0.13 | 0.693 | 0.832 |
| | Depression-only - control | 0.12 | 0.08 | 2240.64 | 1.58 | -0.03 | 0.27 | 0.114 | 0.342 |
| HATA | C-PTSD+Depression - C-PTSD-only | -0.10 | 0.06 | 2251.39 | -1.85 | -0.21 | 0.01 | 0.064^ | 0.384 |
| | C-PTSD+Depression – Depression-only | -0.08 | 0.08 | 2239.41 | -1.03 | -0.23 | 0.07 | 0.302 | 0.604 |
| | C-PTSD+Depression - control | -0.03 | 0.04 | 2252.54 | -0.79 | -0.12 | 0.05 | 0.429 | 0.644 |
| | C-PTSD-only - Depression-only | 0.03 | 0.08 | 2241.86 | 0.31 | -0.13 | 0.19 | 0.757 | 0.757 |
| | C-PTSD-only - control | 0.07 | 0.05 | 2254.47 | 1.33 | -0.03 | 0.17 | 0.183 | 0.549 |
| | Depression-only - control | 0.04 | 0.07 | 2231.10 | 0.61 | -0.10 | 0.18 | 0.539 | 0.647 |

| | | | | | | | | | |
|---|---|---|---|---|---|---|---|---|---|
| Molecular layer | C-PTSD+Depression - C-PTSD-only | -0.13 | 0.05 | 2245.35 | -2.78 | -0.23 | -0.04 | 0.005** | **0.015*** |
| | C-PTSD+Depression – Depression-only | -0.17 | 0.07 | 2243.52 | -2.53 | -0.30 | -0.04 | 0.012* | **0.024*** |
| | C-PTSD+Depression - control | -0.11 | 0.04 | 2254.76 | -2.83 | -0.18 | -0.03 | 0.005** | **0.015*** |
| | C-PTSD-only - Depression-only | -0.03 | 0.07 | 2240.13 | -0.45 | -0.17 | 0.11 | 0.656 | 0.656 |
| | C-PTSD-only - control | 0.03 | 0.05 | 2248.28 | 0.61 | -0.06 | 0.12 | 0.540 | 0.648 |
| | Depression-only - control | 0.06 | 0.06 | 2231.65 | 0.95 | -0.06 | 0.18 | 0.340 | 0.510 |
| Parasubiculum | C-PTSD+Depression - C-PTSD-only | -0.02 | 0.06 | 2255.55 | -0.38 | -0.14 | 0.09 | 0.703 | 0.713 |
| | C-PTSD+Depression – Depression-only | -0.15 | 0.08 | 2246.02 | -1.86 | -0.31 | 0.01 | 0.064^ | 0.320 |
| | C-PTSD+Depression - control | -0.04 | 0.05 | 2198.99 | -0.93 | -0.13 | 0.05 | 0.353 | 0.530 |
| | C-PTSD-only - Depression-only | -0.13 | 0.09 | 2248.42 | -1.47 | -0.30 | 0.04 | 0.143 | 0.320 |
| | C-PTSD-only - control | -0.02 | 0.05 | 2235.85 | -0.37 | -0.13 | 0.09 | 0.713 | 0.713 |
| | Depression-only - control | 0.11 | 0.08 | 2244.85 | 1.41 | -0.04 | 0.26 | 0.160 | 0.320 |
| Presubiculum | C-PTSD+Depression - C-PTSD-only | -0.03 | 0.05 | 2252.21 | -0.53 | -0.13 | 0.07 | 0.598 | 0.598 |
| | C-PTSD+Depression – Depression-only | -0.17 | 0.07 | 2246.11 | -2.42 | -0.31 | -0.03 | 0.016* | 0.096^ |
| | C-PTSD+Depression - control | -0.06 | 0.04 | 2177.56 | -1.58 | -0.14 | 0.02 | 0.115 | 0.173 |
| | C-PTSD-only - Depression-only | -0.14 | 0.08 | 2247.62 | -1.89 | -0.29 | 0.01 | 0.059^ | 0.173 |
| | C-PTSD-only - control | -0.04 | 0.05 | 2239.30 | -0.75 | -0.13 | 0.06 | 0.453 | 0.544 |
| | Depression-only - control | 0.11 | 0.07 | 2252.51 | 1.62 | -0.02 | 0.24 | 0.106 | 0.173 |
| Subiculum | C-PTSD+Depression - C-PTSD-only | -0.06 | 0.05 | 2255.96 | -1.13 | -0.16 | 0.04 | 0.257 | 0.308 |
| | C-PTSD+Depression – Depression-only | -0.17 | 0.07 | 2254.16 | -2.42 | -0.31 | -0.03 | 0.016* | 0.096^ |
| | C-PTSD+Depression - control | -0.05 | 0.04 | 2237.50 | -1.32 | -0.13 | 0.03 | 0.187 | 0.281 |
| | C-PTSD-only - Depression-only | -0.11 | 0.08 | 2255.53 | -1.48 | -0.26 | 0.04 | 0.140 | 0.280 |
| | C-PTSD-only - control | 0.01 | 0.05 | 2249.63 | 0.11 | -0.09 | 0.10 | 0.913 | 0.913 |
| | Depression-only - control | 0.12 | 0.07 | 2243.74 | 1.76 | -0.01 | 0.25 | 0.078^ | 0.234 |
| Tail | C-PTSD+Depression - C-PTSD-only | -0.18 | 0.06 | 2240.84 | -3.05 | -0.30 | -0.06 | 0.002** | **0.006**** |
| | C-PTSD+Depression – Depression-only | -0.17 | 0.08 | 2245.08 | -2.13 | -0.33 | -0.01 | 0.033* | **0.066*** |
| | C-PTSD+Depression - control | -0.16 | 0.05 | 2170.54 | -3.49 | -0.25 | -0.07 | 0.001** | **0.006**** |
| | C-PTSD-only - Depression-only | 0.01 | 0.09 | 2234.42 | 0.11 | -0.16 | 0.18 | 0.915 | 0.915 |
| | C-PTSD-only - control | 0.02 | 0.05 | 2216.32 | 0.37 | -0.09 | 0.13 | 0.709 | 0.915 |
| | Depression-only - control | 0.01 | 0.08 | 2253.54 | 0.15 | -0.14 | 0.16 | 0.883 | 0.915 |

*Note.* Q-values depict FDR-corrected significance, values in bold ink passed the FDR threshold. ^ = 0.05<q<0.10, * = 0.01<q<0.05, ** = 0.001<q<0.01, ***=q<0.001. Text in gray ink denotes pairwise comparisons in regions that did not survive FDR at the omnibus level. C-PTSD+Depression, n=621, C-PTSD-only, n=384, Depression-only, n=138, healthy control, n=1120

**ST20. Moderation of symptom severity by depression and PTSD in all participants**

| A. PTSD symptom severity by depression | | | | | | | | |
|---|---|---|---|---|---|---|---|---|
| | beta | SE | df | t | p | Q | N, Depression | N, No Depression |
| CA1 | -0.15 | 0.06 | 1026.05 | -2.56 | 0.011* | 0.121 | 462 | 565 |
| CA4 | -0.08 | 0.06 | 1024.87 | -1.45 | 0.148 | 0.233 | 462 | 565 |
| CA3 | -0.06 | 0.06 | 1022.32 | -0.99 | 0.321 | 0.392 | 462 | 565 |
| Tail | -0.13 | 0.07 | 1023.48 | -1.95 | 0.051^ | 0.132 | 462 | 565 |
| Subiculum | -0.11 | 0.06 | 1026.91 | -1.88 | 0.060^ | 0.132 | 462 | 565 |
| Presubiculum | -0.08 | 0.06 | 1026.51 | -1.37 | 0.172 | 0.237 | 462 | 565 |
| Parasubiculum | 0.02 | 0.07 | 1026.65 | 0.28 | 0.776 | 0.776 | 462 | 565 |
| HATA | -0.13 | 0.06 | 1016.96 | -2.11 | 0.035* | 0.128 | 462 | 565 |
| Fimbria | -0.03 | 0.07 | 1017.42 | -0.45 | 0.653 | 0.718 | 462 | 565 |
| Molecular Layer | -0.13 | 0.06 | 1025.59 | -2.30 | 0.022* | 0.121 | 462 | 565 |
| Dentate | -0.08 | 0.06 | 1024.66 | -1.49 | 0.136 | 0.233 | 462 | 565 |
| B. Depression symptom severity by C-PTSD | | | | | | | | |
| | beta | SE | df | t | p | Q | N, PTSD | N, No PTSD |
| CA1 | -0.04 | 0.08 | 725.00 | -0.56 | 0.576 | 0.953 | 462 | 565 |
| CA4 | 0.02 | 0.08 | 720.45 | 0.24 | 0.810 | 0.953 | 462 | 565 |
| CA3 | 0.02 | 0.09 | 725.99 | 0.18 | 0.856 | 0.953 | 462 | 565 |
| Tail | 0.02 | 0.09 | 699.52 | 0.21 | 0.834 | 0.953 | 462 | 565 |
| Subiculum | -0.01 | 0.08 | 654.95 | -0.06 | 0.953 | 0.953 | 462 | 565 |
| Presubiculum | -0.01 | 0.08 | 642.26 | -0.18 | 0.858 | 0.953 | 462 | 565 |
| Parasubiculum | -0.01 | 0.09 | 734.74 | -0.15 | 0.883 | 0.953 | 462 | 565 |
| HATA | -0.03 | 0.09 | 732.19 | -0.29 | 0.771 | 0.953 | 462 | 565 |
| Fimbria | 0.03 | 0.09 | 720.35 | 0.32 | 0.749 | 0.953 | 462 | 565 |
| Molecular Layer | -0.01 | 0.07 | 713.07 | -0.14 | 0.891 | 0.953 | 462 | 565 |
| Dentate | 0.01 | 0.08 | 722.34 | 0.17 | 0.866 | 0.953 | 462 | 565 |

*Note.* Q-values depict FDR-corrected significance, values in bold ink passed the FDR threshold. ^ = p<0.10, *** = p <0.001, **=p<0.01, * = p<0.05

**ST21. Pairwise comparisons of PTSD*Depression interactions for subfield composites**

| ROI | Pairwise Groups | beta | SE | df | t | lowerCI | upperCI | p | Pairwise-q |
|---|---|---|---|---|---|---|---|---|---|
| CA1/Sub | **C-PTSD+Depression - C-PTSD-only** | -0.13 | 0.05 | 2241.16 | -2.72 | -0.23 | -0.04 | 0.007** | **0.021*** |
| | **C-PTSD+Depression - Depression-only** | -0.18 | 0.07 | 2249.50 | -2.72 | -0.31 | -0.05 | 0.007** | **0.021*** |
| | **C-PTSD+Depression - control** | -0.10 | 0.04 | 2249.59 | -2.48 | -0.17 | -0.02 | 0.013* | **0.026*** |
| | C-PTSD-only - Depression-only | -0.05 | 0.07 | 2240.88 | -0.67 | -0.19 | 0.09 | 0.505 | 0.505 |
| | C-PTSD-only - control | 0.04 | 0.05 | 2230.01 | 0.84 | -0.05 | 0.13 | 0.398 | 0.478 |
| | Depression-only - control | 0.09 | 0.06 | 2237.31 | 1.37 | -0.04 | 0.21 | 0.170 | 0.255 |
| Complete Dentate | C-PTSD+Depression - C-PTSD-only | -0.12 | 0.05 | 2247.33 | -2.40 | -0.21 | -0.02 | 0.016* | 0.069^ |
| | C-PTSD+Depression - Depression-only | -0.09 | 0.07 | 2235.80 | -1.35 | -0.22 | 0.04 | 0.177 | **0.354** |
| | C-PTSD+Depression - control | -0.09 | 0.04 | 2255.40 | -2.28 | -0.16 | -0.01 | 0.023* | 0.069^ |
| | C-PTSD-only - Depression-only | 0.03 | 0.07 | 2238.39 | 0.39 | -0.11 | 0.17 | 0.698 | 0.838 |
| | C-PTSD-only - control | 0.03 | 0.05 | 2255.00 | 0.67 | -0.06 | 0.12 | 0.505 | 0.758 |
| | Depression-only - control | 0.001 | 0.06 | 2225.02 | 0.04 | -0.12 | 0.13 | 0.967 | 0.967 |
| HP | **C-PTSD+Depression - C-PTSD-only** | -0.16 | 0.05 | 2247.47 | -3.28 | -0.26 | -0.07 | 0.001** | **0.006**** |
| | C-PTSD+Depression - Depression-only | -0.13 | 0.07 | 2239.99 | -1.95 | -0.26 | 0.00 | 0.051^ | 0.102 |
| | **C-PTSD+Depression - control** | -0.10 | 0.04 | 2255.75 | -2.69 | -0.18 | -0.03 | 0.007** | **0.021*** |
| | C-PTSD-only - Depression-only | 0.03 | 0.07 | 2240.41 | 0.43 | -0.11 | 0.17 | 0.667 | 0.667 |
| | C-PTSD-only - control | 0.06 | 0.05 | 2253.37 | 1.27 | -0.03 | 0.15 | 0.204 | 0.306 |
| | Depression-only - control | 0.03 | 0.06 | 2227.92 | 0.43 | -0.10 | 0.15 | 0.666 | 0.667 |
| CA-only | **C-PTSD+Depression - C-PTSD-only** | -0.17 | 0.05 | 2247.97 | -3.46 | -0.27 | -0.07 | 0.001** | **0.006**** |
| | C-PTSD+Depression – Depression-only | -0.14 | 0.07 | 2242.54 | -2.11 | -0.28 | -0.01 | 0.035* | 0.070^ |
| | **C-PTSD+Depression - control** | -0.11 | 0.04 | 2255.02 | -2.75 | -0.18 | -0.03 | 0.006** | **0.018*** |
| | C-PTSD-only - Depression-only | 0.03 | 0.07 | 2241.85 | 0.41 | -0.11 | 0.17 | 0.684 | 0.684 |
| | C-PTSD-only - control | 0.07 | 0.05 | 2252.41 | 1.42 | -0.03 | 0.16 | 0.157 | 0.236 |
| | Depression-only - control | 0.04 | 0.06 | 2230.14 | 0.56 | -0.09 | 0.16 | 0.573 | 0.684 |
| HP/DG | **C-PTSD+Depression - C-PTSD-only** | -0.15 | 0.05 | 2247.07 | -3.13 | -0.25 | -0.06 | 0.002** | **0.012*** |
| | C-PTSD+Depression - Depression-only | -0.12 | 0.07 | 2238.52 | -1.86 | -0.25 | 0.01 | 0.062* | 0.124 |
| | **C-PTSD+Depression - control** | -0.10 | 0.04 | 2255.83 | -2.64 | -0.18 | -0.03 | 0.008** | **0.024*** |
| | C-PTSD-only - Depression-only | 0.03 | 0.07 | 2239.47 | 0.41 | -0.11 | 0.17 | 0.684 | 0.712 |
| | C-PTSD-only - control | 0.05 | 0.05 | 2253.72 | 1.15 | -0.04 | 0.14 | 0.251 | 0.377 |
| | Depression-only - control | 0.02 | 0.06 | 2226.67 | 0.37 | -0.10 | 0.15 | 0.712 | 0.712 |
| Sub-complex | C-PTSD+Depression - C-PTSD-only | -0.05 | 0.05 | 2234.21 | -0.90 | -0.14 | 0.05 | 0.370 | 0.444 |
| | **C-PTSD+Depression - Depression-only** | -0.18 | 0.07 | 2244.39 | -2.63 | -0.32 | -0.05 | 0.008** | **0.048*** |
| | C-PTSD+Depression - control | -0.06 | 0.04 | 2207.96 | -1.53 | -0.14 | 0.02 | 0.126 | 0.189 |
| | C-PTSD-only - Depression-only | -0.14 | 0.07 | 2222.01 | -1.84 | -0.28 | 0.01 | 0.066^ | 0.132 |
| | C-PTSD-only - control | -0.01 | 0.05 | 2225.86 | -0.32 | -0.11 | 0.08 | 0.751 | 0.751 |
| | Depression-only - control | 0.12 | 0.07 | 2247.00 | 1.87 | -0.01 | 0.25 | 0.062^ | 0.062^ |

| | | | | | | | | | |
|---|---|---|---|---|---|---|---|---|---|
| CA/SubPresub/MOL | **C-PTSD+Depression - C-PTSD-only** | -0.13 | 0.05 | 2238.77 | -2.66 | -0.22 | -0.03 | 0.008** | **0.022*** |
| | **C-PTSD+Depression - Depression-only** | -0.17 | 0.07 | 2247.29 | -2.58 | -0.30 | -0.04 | 0.010* | **0.022*** |
| | **C-PTSD+Depression - control** | -0.10 | 0.04 | 2252.44 | -2.56 | -0.17 | -0.02 | 0.011* | **0.022*** |
| | C-PTSD-only - Depression-only | -0.04 | 0.07 | 2237.90 | -0.58 | -0.18 | 0.10 | 0.565 | 0.565 |
| | C-PTSD-only - control | 0.03 | 0.04 | 2229.99 | 0.72 | -0.06 | 0.12 | 0.474 | 0.565 |
| | Depression-only - control | 0.07 | 0.06 | 2235.12 | 1.18 | -0.05 | 0.19 | 0.240 | 0.360 |
| HF | **C-PTSD+Depression - C-PTSD-only** | -0.16 | 0.05 | 2244.65 | -3.27 | -0.26 | -0.06 | 0.001** | **0.006**** |
| | **C-PTSD+Depression - Depression-only** | -0.16 | 0.07 | 2245.93 | -2.35 | -0.29 | -0.03 | 0.019* | **0.038*** |
| | **C-PTSD+Depression - control** | -0.12 | 0.04 | 2253.70 | -3.06 | -0.19 | -0.04 | 0.002** | **0.006**** |
| | C-PTSD-only - Depression-only | 0.004 | 0.07 | 2241.22 | 0.05 | -0.14 | 0.14 | 0.959 | 0.959 |
| | C-PTSD-only - control | 0.04 | 0.05 | 2243.83 | 0.94 | -0.05 | 0.13 | 0.347 | 0.521 |
| | Depression-only - control | 0.04 | 0.06 | 2233.92 | 0.62 | -0.08 | 0.16 | 0.533 | 0.640 |
| Hippocampal Extended | **C-PTSD+Depression - C-PTSD-only** | -0.14 | 0.05 | 2243.00 | -2.86 | -0.23 | -0.04 | 0.004** | **0.015*** |
| | **C-PTSD+Depression - Depression-only** | -0.17 | 0.07 | 2251.14 | -2.55 | -0.29 | -0.04 | 0.011* | **0.022*** |
| | **C-PTSD+Depression - control** | -0.11 | 0.04 | 2255.80 | -2.84 | -0.18 | -0.03 | 0.005** | **0.015*** |
| | C-PTSD-only - Depression-only | -0.03 | 0.07 | 2241.38 | -0.42 | -0.17 | 0.11 | 0.673 | 0.673 |
| | C-PTSD-only - control | 0.03 | 0.04 | 2238.17 | 0.68 | -0.06 | 0.12 | 0.496 | 0.595 |
| | Depression-only - control | 0.06 | 0.06 | 2239.37 | 0.98 | -0.06 | 0.18 | 0.329 | 0.494 |

*Note*. Q-values depict FDR-corrected significance, values in bold ink passed the FDR threshold. ^ = 0.05<q<0.10, * = 0.01<q<0.05, ** = 0.001<q<0.01, ***=q<0.001. Text in gray ink denotes pairwise comparisons in composite regions that did not survive FDR at the omnibus level. Descriptions: CA1/sub=CA1 + subiculum, Complete dentate=CA4+DG, HP (hippocampus proper)=CA1+CA2/3+CA4, CA-only= CA1+CA2/3, HP/DG= CA1+CA2/3+DG, Sub-complex=subiculum + presubiculum + parasubiculum, CA/SubPresub/MOL= CA1 + CA2/3 + subiculum + presubiculum + MOL, HF (hippocampal formation) = CA1 + CA2/3 + CA4 + DG + subiculum + tail, hippo extended= CA1 + CA2/3 + CA4 + DG + subiculum + presubiculum + parasubiculum + tail + MOL + HATA + fimbria. C-PTSD+Depression, n=621, C-PTSD-only, n=384, Depression-only, n=138, healthy control, n=1120

**ST22. Main effects of C-PTSD and Depression by Civilian/Military Status**

| | A. Effect of C-PTSD in Civilians (PTSD= 569, Control=658) | | | | | | C. Effect of Depression in Civilians (Depression=438, control=732) | | | | | |
|---|---|---|---|---|---|---|---|---|---|---|---|---|
| | beta | SE | DF | t | p | q | beta | SE | DF | t | p | q |
| CA1 | -0.08 | 0.05 | 1225.1 | -1.58 | 0.115 | 0.317 | -0.13 | 0.05 | 1164.18 | -2.58 | 0.010* | 0.055^ |
| CA4 | -0.07 | 0.05 | 1223.1 | -1.53 | 0.126 | 0.317 | -0.09 | 0.05 | 1161.92 | -1.96 | 0.050^ | 0.110 |
| CA3 | -0.05 | 0.05 | 1223.7 | -0.98 | 0.326 | 0.448 | -0.07 | 0.05 | 1162.23 | -1.39 | 0.165 | 0.227 |
| Tail | -0.12 | 0.06 | 1213.2 | -2.12 | 0.034* | 0.317 | -0.19 | 0.06 | 1160.42 | -3.28 | 0.001** | **0.011*** |
| Subiculum | -0.05 | 0.05 | 1196.6 | -1.01 | 0.311 | 0.448 | -0.05 | 0.05 | 1160.76 | -1.03 | 0.305 | 0.358 |
| Presubiculum | -0.06 | 0.05 | 1197.8 | -1.32 | 0.188 | 0.345 | -0.05 | 0.05 | 1150.27 | -0.99 | 0.325 | 0.358 |
| Parasubiculum | -0.04 | 0.06 | 1224.7 | -0.73 | 0.463 | 0.509 | 0.02 | 0.06 | 1169.99 | 0.42 | 0.677 | 0.677 |
| HATA | -0.02 | 0.05 | 1224.5 | -0.32 | 0.748 | 0.748 | -0.09 | 0.06 | 1162.83 | -1.54 | 0.123 | 0.222 |
| Fimbria | -0.05 | 0.06 | 1138.4 | -0.90 | 0.369 | 0.451 | -0.08 | 0.06 | 1126.47 | -1.47 | 0.141 | 0.222 |
| Molecular layer | -0.08 | 0.05 | 1225.5 | -1.81 | 0.071^ | 0.317 | -0.10 | 0.05 | 1164.17 | -2.15 | 0.032* | 0.088^ |
| Dentate | -0.07 | 0.05 | 1223.5 | -1.46 | 0.144 | 0.317 | -0.10 | 0.05 | 1162.30 | -2.16 | 0.031* | 0.088^ |
| | **B. Effect of C-PTSD in Military (PTSD=473, control=701)** | | | | | | **D. Effect of Depression in Military (Depression=362, control=724)** | | | | | |
| | beta | SE | DF | t | p | q | beta | SE | DF | t | p | q |
| CA1 | -0.04 | 0.05 | 1172.7 | -0.85 | 0.396 | 0.746 | -0.03 | 0.05 | 1085.09 | -0.65 | 0.516 | 0.747 |
| CA4 | 0.01 | 0.05 | 1173.1 | 0.13 | 0.898 | 0.905 | -0.03 | 0.05 | 1080.13 | -0.54 | 0.592 | 0.747 |
| CA3 | 0.05 | 0.05 | 1173.0 | 1.06 | 0.291 | 0.746 | -0.05 | 0.05 | 1082.69 | -0.89 | 0.374 | 0.747 |
| Tail | -0.04 | 0.06 | 1103.8 | -0.71 | 0.475 | 0.746 | -0.04 | 0.06 | 1071.08 | -0.72 | 0.472 | 0.747 |
| Subiculum | -0.06 | 0.05 | 1158.6 | -1.11 | 0.269 | 0.746 | 0.04 | 0.06 | 1084.53 | 0.63 | 0.527 | 0.747 |
| Presubiculum | -0.07 | 0.05 | 1103.4 | -1.48 | 0.138 | 0.746 | 0.03 | 0.05 | 1063.60 | 0.47 | 0.637 | 0.747 |
| Parasubiculum | -0.05 | 0.06 | 1100.4 | -0.91 | 0.361 | 0.746 | -0.02 | 0.06 | 1076.54 | -0.41 | 0.682 | 0.747 |
| HATA | 0.02 | 0.05 | 1171.1 | 0.40 | 0.686 | 0.905 | 0.05 | 0.05 | 1083.52 | 0.90 | 0.367 | 0.747 |
| Fimbria | 0.01 | 0.06 | 1163.9 | 0.17 | 0.864 | 0.905 | 0.12 | 0.06 | 1085.22 | 2.00 | 0.046* | 0.506 |
| Molecular layer | -0.04 | 0.05 | 1173.4 | -0.79 | 0.430 | 0.746 | -0.03 | 0.05 | 1083.97 | -0.55 | 0.583 | 0.747 |
| Dentate | -0.01 | 0.05 | 1173.1 | -0.12 | 0.905 | 0.905 | -0.02 | 0.05 | 1080.11 | -0.32 | 0.747 | 0.747 |

*Note.* Analyses of C-PTSD did not covary for depression, and analyses of depression did not covary for C-PTSD. Both sets of analyses covaried for sex. Q-values depict FDR-corrected significance, values in bold ink passed the FDR threshold. ^ = p<0.10, *** = p <0.001, **=p<0.01, * = p<0.05

**ST23. Main effects of C-PTSD and Depression by Sex**

| | A. C-PTSD in Females (PTSD=440, control=486) | | | | | | C. Depression in Females (Depression=365, control=513) | | | | | |
|---|---|---|---|---|---|---|---|---|---|---|---|---|
| | beta | SE | DF | t | p | q | beta | SE | DF | t | p | q |
| CA1 | -0.06 | 0.06 | 862.60 | -1.05 | 0.296 | 0.582 | -0.16 | 0.06 | 852.42 | -2.74 | 0.006** | **0.033*** |
| CA4 | -0.07 | 0.06 | 925.58 | -1.15 | 0.249 | 0.582 | -0.12 | 0.06 | 876.18 | -2.01 | 0.045* | 0.099^ |
| CA3 | -0.07 | 0.06 | 925.72 | -1.10 | 0.271 | 0.582 | -0.08 | 0.06 | 875.32 | -1.29 | 0.198 | 0.272 |
| Tail | -0.11 | 0.06 | 895.68 | -1.71 | 0.088^ | 0.582 | -0.23 | 0.06 | 859.05 | -3.65 | 0.0003*** | **0.003**** |
| Subiculum | -0.04 | 0.05 | 885.39 | -0.71 | 0.476 | 0.582 | -0.07 | 0.05 | 864.97 | -1.34 | 0.180 | 0.272 |
| Presubiculum | -0.05 | 0.05 | 907.66 | -0.83 | 0.406 | 0.582 | -0.07 | 0.06 | 868.96 | -1.20 | 0.229 | 0.280 |
| Parasubiculum | -0.05 | 0.07 | 922.82 | -0.79 | 0.428 | 0.582 | -0.03 | 0.07 | 876.56 | -0.51 | 0.607 | 0.607 |
| HATA | 0.01 | 0.07 | 910.37 | 0.17 | 0.864 | 0.582 | -0.10 | 0.07 | 866.82 | -1.60 | 0.110 | 0.202 |
| Fimbria | 0.01 | 0.06 | 860.56 | 0.23 | 0.818 | 0.582 | -0.05 | 0.06 | 772.75 | -0.75 | 0.453 | 0.498 |
| Molecular layer | -0.08 | 0.05 | 911.24 | -1.41 | 0.160 | 0.582 | -0.13 | 0.05 | 867.03 | -2.31 | 0.021* | 0.077^ |
| Dentate | -0.05 | 0.06 | 925.11 | -0.91 | 0.363 | 0.582 | -0.12 | 0.06 | 876.21 | -2.06 | 0.040* | 0.099^ |
| | B. C-PTSD in Males (PTSD=587, control=849) | | | | | | D. Depression in Males (Depression=423, control=916) | | | | | |
| | beta | SE | DF | t | p | q | beta | SE | DF | t | p | q |
| CA1 | -0.08 | 0.04 | 1435.11 | -1.77 | 0.076^ | 0.217 | -0.07 | 0.05 | 1333.70 | -1.47 | 0.142 | 0.363 |
| CA4 | -0.05 | 0.04 | 1428.88 | -1.18 | 0.240 | 0.330 | -0.07 | 0.04 | 1325.39 | -1.54 | 0.123 | 0.363 |
| CA3 | 0.00 | 0.04 | 1431.19 | 0.00 | 0.998 | 0.998 | -0.10 | 0.05 | 1329.10 | -2.04 | 0.042* | 0.363 |
| Tail | -0.09 | 0.05 | 1418.14 | -1.78 | 0.075^ | 0.217 | -0.08 | 0.06 | 1326.84 | -1.39 | 0.165 | 0.363 |
| Subiculum | -0.07 | 0.05 | 1373.18 | -1.62 | 0.106 | 0.233 | 0.01 | 0.05 | 1338.57 | 0.26 | 0.797 | 0.919 |
| Presubiculum | -0.09 | 0.05 | 1428.05 | -1.99 | 0.047* | 0.217 | 0.01 | 0.05 | 1333.09 | 0.27 | 0.786 | 0.919 |
| Parasubiculum | -0.06 | 0.05 | 1418.87 | -1.18 | 0.238 | 0.330 | 0.01 | 0.05 | 1330.36 | 0.10 | 0.919 | 0.919 |
| HATA | -0.03 | 0.04 | 1429.42 | -0.76 | 0.448 | 0.493 | -0.01 | 0.05 | 1333.99 | -0.11 | 0.910 | 0.919 |
| Fimbria | -0.04 | 0.05 | 1429.74 | -0.79 | 0.432 | 0.493 | 0.05 | 0.05 | 1331.49 | 0.90 | 0.368 | 0.578 |
| Molecular layer | -0.07 | 0.04 | 1432.74 | -1.76 | 0.079^ | 0.217 | -0.06 | 0.05 | 1329.92 | -1.29 | 0.198 | 0.363 |
| Dentate | -0.06 | 0.04 | 1429.11 | -1.44 | 0.149 | 0.273 | -0.06 | 0.04 | 1325.13 | -1.38 | 0.168 | 0.363 |

*Note.* Analyses of C-PTSD did not covary for depression, and analyses of depression did not covary for C-PTSD. Both sets of analyses covaried for civilian/military status. Q-values depict FDR-corrected significance, values in bold ink passed the FDR threshold. ^ = 0.05<q<0.10, * = 0.01<q<0.05, ** = 0.001<q<0.01, ***=q<0.001.

**ST24. Main effects of C-PTSD and Depression by Sex and Military Status**

| | A. C-PTSD in Civilian Females (PTSD=440, control=485) | | | | | | C. Depression in Civilian Females (Depression=338, control=461) | | | | | |
|---|---|---|---|---|---|---|---|---|---|---|---|---|
| | beta | SE | DF | t | p | q | beta | SE | DF | t | p | q |
| CA1 | -0.04 | 0.06 | 831.00 | -0.70 | 0.486 | 0.856 | -0.14 | 0.06 | 750.17 | -2.40 | 0.017* | 0.092^ |
| CA4 | -0.04 | 0.06 | 829.78 | -0.62 | 0.536 | 0.856 | -0.10 | 0.06 | 792.34 | -1.65 | 0.099^ | 0.204 |
| CA3 | -0.03 | 0.06 | 840.13 | -0.48 | 0.634 | 0.856 | -0.06 | 0.06 | 793.89 | -0.88 | 0.377 | 0.492 |
| Tail | -0.11 | 0.07 | 827.13 | -1.60 | 0.109 | 0.856 | -0.24 | 0.07 | 774.11 | -3.55 | 0.0004*** | **0.004**** |
| Subiculum | -0.02 | 0.06 | 807.09 | -0.37 | 0.714 | 0.856 | -0.05 | 0.06 | 782.23 | -0.97 | 0.332 | 0.492 |
| Presubiculum | -0.03 | 0.06 | 836.21 | -0.46 | 0.649 | 0.856 | -0.05 | 0.06 | 792.89 | -0.84 | 0.403 | 0.492 |
| Parasubiculum | -0.02 | 0.07 | 844.73 | -0.24 | 0.813 | 0.856 | 0.00 | 0.07 | 796.24 | 0.04 | 0.966 | 0.966 |
| HATA | 0.03 | 0.07 | 828.05 | 0.38 | 0.706 | 0.856 | -0.11 | 0.07 | 770.19 | -1.60 | 0.111 | 0.204 |
| Fimbria | 0.01 | 0.07 | 783.08 | 0.18 | 0.856 | 0.856 | -0.04 | 0.07 | 361.94 | -0.65 | 0.513 | 0.565 |
| Molecular layer | -0.05 | 0.06 | 834.74 | -0.93 | 0.355 | 0.856 | -0.11 | 0.06 | 738.42 | -1.86 | 0.063^ | 0.204 |
| Dentate | -0.02 | 0.06 | 823.05 | -0.38 | 0.706 | 0.856 | -0.10 | 0.06 | 790.93 | -1.68 | 0.094^ | 0.204 |
| | **B. C-PTSD in Military Males (PTSD=450, control=646)** | | | | | | **D. Depression in Military Males (Depression=334, control=672)** | | | | | |
| | beta | SE | DF | t | p | q | beta | SE | DF | t | p | q |
| CA1 | -0.01 | 0.05 | 1088.20 | -0.20 | 0.838 | 0.917 | 0.0004 | 0.06 | 1005.59 | 0.01 | 0.994 | 0.994 |
| CA4 | 0.04 | 0.05 | 1092.09 | 0.76 | 0.445 | 0.839 | 0.0005 | 0.05 | 1002.14 | 0.01 | 0.993 | 0.994 |
| CA3 | 0.09 | 0.05 | 1092.42 | 1.71 | 0.087^ | 0.839 | -0.02 | 0.06 | 1004.04 | -0.41 | 0.685 | 0.994 |
| Tail | -0.04 | 0.06 | 1034.58 | -0.69 | 0.493 | 0.839 | -0.03 | 0.06 | 989.57 | -0.47 | 0.639 | 0.994 |
| Subiculum | -0.03 | 0.05 | 1067.87 | -0.63 | 0.529 | 0.839 | 0.08 | 0.06 | 1004.29 | 1.34 | 0.181 | 0.840 |
| Presubiculum | -0.06 | 0.05 | 1014.92 | -1.20 | 0.232 | 0.839 | 0.05 | 0.06 | 977.86 | 0.95 | 0.343 | 0.943 |
| Parasubiculum | -0.03 | 0.06 | 1047.57 | -0.51 | 0.610 | 0.839 | -0.004 | 0.06 | 995.33 | -0.07 | 0.945 | 0.994 |
| HATA | 0.04 | 0.05 | 1088.98 | 0.82 | 0.411 | 0.839 | 0.07 | 0.06 | 1003.90 | 1.20 | 0.229 | 0.839 |
| Fimbria | 0.01 | 0.06 | 1082.52 | 0.19 | 0.850 | 0.917 | 0.14 | 0.06 | 1005.39 | 2.18 | 0.030* | 0.330 |
| Molecular layer | -0.01 | 0.05 | 1092.83 | -0.10 | 0.917 | 0.917 | 0.01 | 0.05 | 1004.95 | 0.18 | 0.859 | 0.994 |
| Dentate | 0.02 | 0.05 | 1092.16 | 0.51 | 0.609 | 0.839 | 0.01 | 0.05 | 1002.34 | 0.25 | 0.804 | 0.994 |

*Note.* Analyses of C-PTSD did not covary for depression, and analyses of depression did not covary for C-PTSD. Q-values depict FDR-corrected significance, values in bold ink passed the FDR threshold. ^ = 0.05<q<0.10, * = 0.01<q<0.05, ** = 0.001<q<0.01, ***=q<0.001.

**ST25. Interactions between C-PTSD and depression by civilian and military background**

| | | | | | | | C-PTSD - only | Depression-only | C-PTSD +depression | Control |
|---|---|---|---|---|---|---|---|---|---|---|
| **A. Civilians** | beta | SE | DF | t | p | q | N | N | N | N |
| CA1 | -0.030 | 0.12 | 1159.26 | -0.24 | 0.809 | 0.974 | 203 | 48 | 348 | 573 |
| CA4 | -0.049 | 0.12 | 1155.26 | -0.40 | 0.688 | 0.974 | 203 | 48 | 348 | 573 |
| CA3 | -0.147 | 0.13 | 1155.79 | -1.13 | 0.259 | 0.974 | 203 | 48 | 348 | 573 |
| Tail | -0.154 | 0.14 | 1165.40 | -1.07 | 0.284 | 0.974 | 203 | 48 | 348 | 573 |
| Subiculum | -0.004 | 0.12 | 1162.69 | -0.03 | 0.974 | 0.974 | 203 | 48 | 348 | 573 |
| Presubiculum | -0.011 | 0.12 | 1168.30 | -0.09 | 0.928 | 0.974 | 203 | 48 | 348 | 573 |
| Parasubiculum | -0.110 | 0.15 | 1164.13 | -0.75 | 0.451 | 0.974 | 203 | 48 | 348 | 573 |
| HATA | -0.082 | 0.14 | 1157.92 | -0.58 | 0.559 | 0.974 | 203 | 48 | 348 | 573 |
| Fimbria | 0.110 | 0.15 | 1169.00 | 0.76 | 0.450 | 0.974 | 203 | 48 | 348 | 573 |
| Molecular layer | -0.041 | 0.12 | 1157.51 | -0.35 | 0.728 | 0.974 | 203 | 48 | 348 | 573 |
| Dentate gyrus | -0.023 | 0.12 | 1155.53 | -0.20 | 0.845 | 0.974 | 203 | 48 | 348 | 573 |
| **B. Military** | | | | | | | C-PTSD - only | Depression-only | C-PTSD +depression | Control |
| | beta | SE | DF | t | p | q | N | N | N | N |
| CA1 | -0.35 | 0.12 | 1080.64 | -3.01 | 0.003** | **0.033*** | 178 | 90 | 261 | 523 |
| CA4 | -0.12 | 0.11 | 1075.55 | -1.09 | 0.275 | 0.378 | 178 | 90 | 261 | 523 |
| CA3 | -0.04 | 0.12 | 1077.85 | -0.31 | 0.760 | 0.760 | 178 | 90 | 261 | 523 |
| Tail | -0.18 | 0.13 | 1068.37 | -1.32 | 0.188 | 0.295 | 178 | 90 | 261 | 523 |
| Subiculum | -0.30 | 0.13 | 1077.41 | -2.38 | 0.017* | 0.062^ | 178 | 90 | 261 | 523 |
| Presubiculum | -0.21 | 0.12 | 1050.70 | -1.74 | 0.083^ | 0.183 | 178 | 90 | 261 | 523 |
| Parasubiculum | -0.12 | 0.13 | 1078.52 | -0.93 | 0.352 | 0.387 | 178 | 90 | 261 | 523 |
| HATA | -0.11 | 0.12 | 1078.57 | -0.93 | 0.352 | 0.387 | 178 | 90 | 261 | 523 |
| Fimbria | -0.30 | 0.13 | 1081.36 | -2.25 | 0.025* | 0.069^ | 178 | 90 | 261 | 523 |
| Molecular layer | -0.28 | 0.11 | 1078.76 | -2.49 | 0.013* | 0.062^ | 178 | 90 | 261 | 523 |
| Dentate gyrus | -0.15 | 0.11 | 1074.99 | -1.38 | 0.168 | 0.295 | 178 | 90 | 261 | 523 |

Note. Analysis models in civilians failed to converge for the dentate gyrus. Q-values depict FDR-corrected significance, values in bold ink passed the FDR threshold.
^ = 0.05<q<0.10, * = 0.01<q<0.05, ** = 0.001<q<0.01, ***=q<0.001.

**ST26. Interactions between C-PTSD and depression by sex**

**A. Males**

| | beta | SE | DF | t | p | q | C-PTSD -only N | Depression-only N | C-PTSD +depression N | Control N |
|---|---|---|---|---|---|---|---|---|---|---|
| CA1 | -0.33 | 0.11 | 1325.26 | -3.09 | 0.0020** | **0.022*** | 258 | 97 | 322 | 705 |
| CA4 | -0.11 | 0.10 | 1317.60 | -1.15 | 0.2504 | 0.275 | 258 | 97 | 322 | 705 |
| CA3 | -0.07 | 0.11 | 1320.46 | -0.61 | 0.5450 | 0.545 | 258 | 97 | 322 | 705 |
| Tail | -0.20 | 0.12 | 1337.76 | -1.58 | 0.1147 | 0.180 | 258 | 97 | 322 | 705 |
| Subiculum | -0.30 | 0.11 | 1330.25 | -2.67 | 0.0076** | **0.027*** | 258 | 97 | 322 | 705 |
| Presubiculum | -0.18 | 0.11 | 1336.65 | -1.58 | 0.1137 | 0.180 | 258 | 97 | 322 | 705 |
| Parasubiculum | -0.18 | 0.12 | 1335.86 | -1.47 | 0.1412 | 0.194 | 258 | 97 | 322 | 705 |
| HATA | -0.20 | 0.11 | 1325.42 | -1.84 | 0.0662^ | 0.146 | 258 | 97 | 322 | 705 |
| Fimbria | -0.35 | 0.12 | 1327.83 | -2.89 | 0.0040** | **0.022*** | 258 | 97 | 322 | 705 |
| Molecular layer | -0.26 | 0.10 | 1321.33 | -2.59 | 0.0098** | **0.027*** | 258 | 97 | 322 | 705 |
| Dentate gyrus | -0.14 | 0.10 | 1317.07 | -1.38 | 0.1687 | 0.206 | 258 | 97 | 322 | 705 |

**B. Females**

| | beta | SE | DF | t | p | q | C-PTSD -only N | Depression-only N | C-PTSD +depression N | Control N |
|---|---|---|---|---|---|---|---|---|---|---|
| CA1 | -0.10 | 0.15 | 873.80 | -0.67 | 0.502 | 0.760 | 126 | 41 | 299 | 415 |
| CA4 | -0.11 | 0.15 | 867.21 | -0.71 | 0.480 | 0.760 | 126 | 41 | 299 | 415 |
| CA3 | -0.19 | 0.16 | 866.60 | -1.19 | 0.233 | 0.760 | 126 | 41 | 299 | 415 |
| Tail | -0.20 | 0.16 | 877.40 | -1.24 | 0.216 | 0.760 | 126 | 41 | 299 | 415 |
| Subiculum | 0.02 | 0.14 | 873.20 | 0.11 | 0.910 | 0.760 | 126 | 41 | 299 | 415 |
| Presubiculum | -0.07 | 0.14 | 873.73 | -0.52 | 0.600 | 0.760 | 126 | 41 | 299 | 415 |
| Parasubiculum | -0.01 | 0.17 | 868.11 | -0.04 | 0.969 | 0.760 | 126 | 41 | 299 | 415 |
| HATA | -0.08 | 0.17 | 874.51 | -0.49 | 0.622 | 0.760 | 126 | 41 | 299 | 415 |
| Fimbria | 0.23 | 0.17 | 876.80 | 1.38 | 0.167 | 0.760 | 126 | 41 | 299 | 415 |
| Molecular layer | -0.09 | 0.14 | 871.40 | -0.66 | 0.510 | 0.760 | 126 | 41 | 299 | 415 |
| Dentate gyrus | -0.09 | 0.15 | 867.66 | -0.64 | 0.525 | 0.760 | 126 | 41 | 299 | 415 |

Note. Civilian/military status was used as a covariate. Analysis models in females failed to converge in the presubiculum. Q-values depict FDR-corrected significance, values in bold ink passed the FDR threshold. ^ = 0.05<q<0.10, * = 0.01<q<0.05, ** = 0.001<q<0.01, ***=q<0.001.

**ST27. Interactions between C-PTSD and depression in military and male subgroups**

| | | | | | | | C-PTSD - only | Depression-only | C-PTSD +depression | Control |
|---|---|---|---|---|---|---|---|---|---|---|
| **A. Military males** | | | | | | | | | | |
| | beta | SE | DF | t | p | q | N | N | N | N |
| CA1 | -0.32 | 0.12 | 1001.41 | -2.53 | 0.011* | 0.121 | 175 | 84 | 239 | 474 |
| CA4 | -0.06 | 0.12 | 997.55 | -0.52 | 0.601 | 0.661 | 175 | 84 | 239 | 474 |
| CA3 | 0.01 | 0.13 | 999.50 | 0.11 | 0.916 | 0.916 | 175 | 84 | 239 | 474 |
| Tail | -0.17 | 0.14 | 984.15 | -1.20 | 0.231 | 0.427 | 175 | 84 | 239 | 474 |
| Subiculum | -0.24 | 0.13 | 996.59 | -1.84 | 0.067^ | 0.184 | 175 | 84 | 239 | 474 |
| Presubiculum | -0.15 | 0.13 | 959.87 | -1.19 | 0.233 | 0.427 | 175 | 84 | 239 | 474 |
| Parasubiculum | -0.11 | 0.14 | 995.81 | -0.81 | 0.420 | 0.578 | 175 | 84 | 239 | 474 |
| HATA | -0.08 | 0.13 | 998.75 | -0.60 | 0.549 | 0.661 | 175 | 84 | 239 | 474 |
| Fimbria | -0.32 | 0.14 | 1001.77 | -2.29 | 0.022* | 0.121 | 175 | 84 | 239 | 474 |
| Molecular layer | -0.22 | 0.12 | 1000.25 | -1.90 | 0.058^ | 0.184 | 175 | 84 | 239 | 474 |
| Dentate gyrus | -0.10 | 0.12 | 997.19 | -0.84 | 0.404 | 0.578 | 175 | 84 | 239 | 474 |
| **B. Males, covarying the effects of mTBI** | | | | | | | | | | |
| | | | | | | | C-PTSD - only | Depression-only | C-PTSD +depression | Control |
| | beta | SE | DF | t | p | q | N | N | N | N |
| CA1 | -0.22 | 0.16 | 584.02 | -1.33 | 0.183 | 0.403 | 100 | 52 | 106 | 351 |
| CA4 | 0.03 | 0.14 | 580.01 | 0.20 | 0.842 | 0.842 | 100 | 52 | 106 | 351 |
| CA3 | 0.16 | 0.16 | 580.37 | 1.01 | 0.315 | 0.454 | 100 | 52 | 106 | 351 |
| Tail | -0.31 | 0.19 | 582.44 | -1.64 | 0.102 | 0.327 | 100 | 52 | 106 | 351 |
| Subiculum | -0.26 | 0.17 | 583.68 | -1.56 | 0.119 | 0.327 | 100 | 52 | 106 | 351 |
| Presubiculum | -0.17 | 0.17 | 584.51 | -0.97 | 0.330 | 0.454 | 100 | 52 | 106 | 351 |
| Parasubiculum | -0.20 | 0.19 | 584.33 | -1.04 | 0.297 | 0.454 | 100 | 52 | 106 | 351 |
| HATA | -0.28 | 0.17 | 584.10 | -1.70 | 0.089^ | 0.327 | 100 | 52 | 106 | 351 |
| Fimbria | -0.51 | 0.18 | 583.26 | -2.78 | 0.006** | 0.066^ | 100 | 52 | 106 | 351 |
| Molecular layer | -0.13 | 0.15 | 581.53 | -0.88 | 0.379 | 0.463 | 100 | 52 | 106 | 351 |
| Dentate gyrus | -0.04 | 0.15 | 580.83 | -0.25 | 0.802 | 0.842 | 100 | 52 | 106 | 351 |

Note. In addition to covarying for TBI, civilian/military status was also used as a covariate. 53% of this subsample indicated a history of a mild TBI. Q-values depict FDR-corrected significance, values in bold ink passed the FDR threshold. ^ = 0.05<q<0.10, * = 0.01<q<0.05, ** = 0.001<q<0.01, ***=q<0.001.

| ST28. Moderation of PTSD symptom severity by depression | | | | | | | | |
|---|---|---|---|---|---|---|---|---|
| **A. Military** | | | | | | | | |
| | beta | SE | df | t | p | Q | N, Depression | N, No Depression |
| CA1 | -0.22 | 0.08 | 543.07 | -2.77 | 0.006** | 0.056^ | 233 | 312 |
| CA4 | -0.09 | 0.08 | 544.33 | -1.17 | 0.243 | 0.414 | 233 | 312 |
| CA3 | -0.07 | 0.09 | 534.39 | -0.81 | 0.419 | 0.461 | 233 | 312 |
| Tail | -0.13 | 0.09 | 514.35 | -1.54 | 0.124 | 0.274 | 233 | 312 |
| Subiculum | -0.20 | 0.09 | 544.45 | -2.29 | 0.022* | 0.082^ | 233 | 312 |
| Presubiculum | -0.15 | 0.08 | 527.23 | -1.94 | 0.053^ | 0.147 | 233 | 312 |
| Parasubiculum | -0.08 | 0.08 | 541.35 | -0.96 | 0.339 | 0.414 | 233 | 312 |
| HATA | -0.08 | 0.08 | 539.81 | -0.96 | 0.337 | 0.414 | 233 | 312 |
| Fimbria | -0.02 | 0.10 | 540.34 | -0.21 | 0.836 | 0.836 | 233 | 312 |
| Molecular Layer | -0.19 | 0.07 | 544.32 | -2.58 | 0.010* | 0.056^ | 233 | 312 |
| Dentate | -0.08 | 0.07 | 543.89 | -1.07 | 0.287 | 0.414 | 233 | 312 |
| **B. Civilians** | | | | | | | | |
| | beta | SE | df | t | p | Q | N, Depression | N, No Depression |
| CA1 | -0.02 | 0.09 | 477.88 | -0.17 | 0.864 | 0.995 | 229 | 253 |
| CA4 | -0.05 | 0.09 | 475.84 | -0.58 | 0.561 | 0.995 | 229 | 253 |
| CA3 | -0.04 | 0.10 | 475.63 | -0.43 | 0.665 | 0.995 | 229 | 253 |
| Tail | -0.08 | 0.11 | 478.15 | -0.71 | 0.480 | 0.995 | 229 | 253 |
| Subiculum | 0.00 | 0.09 | 481.63 | -0.01 | 0.995 | 0.995 | 229 | 253 |
| Presubiculum | 0.01 | 0.09 | 480.84 | 0.12 | 0.908 | 0.995 | 229 | 253 |
| Parasubiculum | 0.18 | 0.11 | 481.26 | 1.61 | 0.108 | 0.704 | 229 | 253 |
| HATA | -0.15 | 0.10 | 475.43 | -1.52 | 0.128 | 0.704 | 229 | 253 |
| Fimbria | -0.03 | 0.11 | 480.53 | -0.32 | 0.751 | 0.995 | 229 | 253 |
| Molecular Layer | -0.02 | 0.09 | 477.24 | -0.21 | 0.834 | 0.995 | 229 | 253 |
| Dentate | -0.06 | 0.09 | 475.90 | -0.73 | 0.466 | 0.995 | 229 | 253 |
| **C. Females** | | | | | | | | |
| | beta | SE | df | t | p | Q | N, Depression | N, No Depression |
| CA1 | -0.10 | 0.12 | 332.95 | -0.88 | 0.377 | 0.596 | 181 | 152 |
| CA4 | -0.18 | 0.11 | 333.00 | -1.61 | 0.108 | 0.396 | 181 | 152 |
| CA3 | -0.13 | 0.13 | 325.74 | -1.00 | 0.318 | 0.596 | 181 | 152 |

| | | | | | | | | |
|---|---|---|---|---|---|---|---|---|
| Tail | -0.01 | 0.13 | 331.75 | -0.11 | 0.914 | 0.914 | 181 | 152 |
| Subiculum | -0.08 | 0.12 | 331.76 | -0.66 | 0.509 | 0.700 | 181 | 152 |
| Presubiculum | -0.06 | 0.12 | 330.56 | -0.55 | 0.584 | 0.714 | 181 | 152 |
| Parasubiculum | 0.25 | 0.14 | 332.92 | 1.78 | 0.077^ | 0.396 | 181 | 152 |
| HATA | -0.12 | 0.13 | 332.08 | -0.88 | 0.379 | 0.596 | 181 | 152 |
| Fimbria | -0.05 | 0.14 | 331.76 | -0.37 | 0.712 | 0.783 | 181 | 152 |
| Molecular Layer | -0.13 | 0.11 | 332.53 | -1.22 | 0.225 | 0.596 | 181 | 152 |
| Dentate | -0.19 | 0.11 | 333.00 | -1.75 | 0.082 | 0.396 | 181 | 152 |
| **D. Males** | | | | | | | | |
| | beta | SE | df | t | p | Q | N, Depression | N, No Depression |
| CA1 | -0.16 | 0.07 | 693.89 | -2.25 | 0.025* | 0.165 | 281 | 413 |
| CA4 | -0.05 | 0.07 | 691.66 | -0.78 | 0.436 | 0.513 | 281 | 413 |
| CA3 | -0.05 | 0.08 | 691.00 | -0.65 | 0.513 | 0.513 | 281 | 413 |
| Tail | -0.15 | 0.08 | 678.31 | -1.88 | 0.061^ | 0.165 | 281 | 413 |
| Subiculum | -0.14 | 0.08 | 692.74 | -1.79 | 0.075^ | 0.165 | 281 | 413 |
| Presubiculum | -0.10 | 0.07 | 686.84 | -1.33 | 0.185 | 0.339 | 281 | 413 |
| Parasubiculum | -0.08 | 0.08 | 692.77 | -0.99 | 0.324 | 0.509 | 281 | 413 |
| HATA | -0.14 | 0.07 | 687.25 | -1.92 | 0.055^ | 0.165 | 281 | 413 |
| Fimbria | -0.06 | 0.08 | 687.64 | -0.73 | 0.467 | 0.513 | 281 | 413 |
| Molecular Layer | -0.13 | 0.07 | 692.87 | -1.91 | 0.056^ | 0.165 | 281 | 413 |
| Dentate | -0.06 | 0.07 | 691.60 | -0.84 | 0.401 | 0.513 | 281 | 413 |
| **E. Male Military** | | | | | | | | |
| | beta | SE | df | t | p | Q | N, Depression | N, No Depression |
| CA1 | -0.22 | 0.08 | 507.84 | -2.70 | 0.007** | 0.077^ | 217 | 291 |
| CA4 | -0.07 | 0.08 | 507.59 | -0.96 | 0.339 | 0.452 | 217 | 291 |
| CA3 | -0.06 | 0.09 | 505.06 | -0.68 | 0.498 | 0.548 | 217 | 291 |
| Tail | -0.13 | 0.09 | 460.24 | -1.40 | 0.162 | 0.356 | 217 | 291 |
| Subiculum | -0.20 | 0.09 | 507.23 | -2.21 | 0.027* | 0.099^ | 217 | 291 |
| Presubiculum | -0.15 | 0.08 | 489.51 | -1.83 | 0.068^ | 0.187 | 217 | 291 |
| Parasubiculum | -0.08 | 0.09 | 503.85 | -0.98 | 0.327 | 0.452 | 217 | 291 |
| HATA | -0.09 | 0.08 | 503.06 | -1.05 | 0.295 | 0.452 | 217 | 291 |
| Fimbria | -0.03 | 0.10 | 503.20 | -0.34 | 0.735 | 0.735 | 217 | 291 |
| Molecular Layer | -0.19 | 0.08 | 507.57 | -2.40 | 0.017* | 0.094^ | 217 | 291 |
| Dentate | -0.07 | 0.08 | 507.41 | -0.90 | 0.370 | 0.452 | 217 | 291 |
| **F. Female Civilians** | | | | | | | | |
| | beta | SE | df | t | p | Q | N, Depression | N, No Depression |

| | | | | | | | | |
|---|---|---|---|---|---|---|---|---|
| CA1 | -0.09 | 0.13 | 295.92 | -0.69 | 0.490 | 0.665 | 165 | 131 |
| CA4 | -0.17 | 0.12 | 296.00 | -1.42 | 0.156 | 0.572 | 165 | 131 |
| CA3 | -0.16 | 0.14 | 285.34 | -1.14 | 0.257 | 0.654 | 165 | 131 |
| Tail | 0.02 | 0.15 | 294.74 | 0.13 | 0.894 | 0.894 | 165 | 131 |
| Subiculum | -0.08 | 0.13 | 294.91 | -0.61 | 0.544 | 0.665 | 165 | 131 |
| Presubiculum | -0.03 | 0.12 | 292.98 | -0.28 | 0.778 | 0.856 | 165 | 131 |
| Parasubiculum | 0.28 | 0.15 | 295.99 | 1.89 | 0.060^ | 0.572 | 165 | 131 |
| HATA | -0.12 | 0.14 | 295.18 | -0.81 | 0.416 | 0.654 | 165 | 131 |
| Fimbria | -0.13 | 0.14 | 272.76 | -0.91 | 0.362 | 0.654 | 165 | 131 |
| Molecular Layer | -0.11 | 0.12 | 291.86 | -0.97 | 0.332 | 0.654 | 165 | 131 |
| Dentate | -0.18 | 0.12 | 296.00 | -1.57 | 0.118 | 0.572 | 165 | 131 |

*Note.* Analyses conducted separately in civilians and military included sex covariates (sex, age*sex, and $age^2$*sex), and analyses conducted separately in males in females included a covariate for civilian/military status. Q-values depict FDR-corrected significance, values in bold ink passed the FDR threshold. ^ = 0.05<q<0.10, * = 0.01<q<0.05, ** = 0.001<q<0.01, ***=q<0.001.

**ST29. Moderation of depression symptom severity by Current PTSD**

| | beta | SE | df | t | p | Q | N, PTSD | N, No PTSD |
|---|---|---|---|---|---|---|---|---|
| **A. Military** | | | | | | | | |
| CA1 | 0.003 | 0.101 | 273.988 | 0.028 | 0.977 | 0.977 | 137 | 210 |
| CA4 | 0.126 | 0.101 | 261.105 | 1.254 | 0.211 | 0.749 | 137 | 210 |
| CA3 | 0.082 | 0.113 | 261.157 | 0.728 | 0.467 | 0.749 | 137 | 210 |
| Tail | 0.205 | 0.113 | 207.874 | 1.821 | 0.070^ | 0.749 | 137 | 210 |
| Subiculum | 0.068 | 0.113 | 229.935 | 0.606 | 0.545 | 0.749 | 137 | 210 |
| Presubiculum | -0.021 | 0.107 | 217.939 | -0.196 | 0.844 | 0.928 | 137 | 210 |
| Parasubiculum | -0.081 | 0.108 | 211.409 | -0.757 | 0.450 | 0.749 | 137 | 210 |
| HATA | 0.051 | 0.107 | 320.069 | 0.479 | 0.632 | 0.772 | 137 | 210 |
| Fimbria | 0.132 | 0.125 | 292.390 | 1.057 | 0.291 | 0.749 | 137 | 210 |
| Molecular Layer | 0.062 | 0.099 | 278.561 | 0.626 | 0.532 | 0.749 | 137 | 210 |
| Dentate | 0.116 | 0.099 | 287.332 | 1.170 | 0.243 | 0.749 | 137 | 210 |
| **B. Civilians** | | | | | | | | |
| | beta | SE | df | t | p | Q | N, PTSD | N, No PTSD |
| CA1 | -0.01 | 0.15 | 387.73 | -0.05 | 0.960 | 0.960 | 256 | 132 |
| CA4 | -0.12 | 0.15 | 387.99 | -0.75 | 0.455 | 0.854 | 256 | 132 |
| CA3 | -0.20 | 0.17 | 387.41 | -1.23 | 0.218 | 0.809 | 256 | 132 |
| Tail | -0.19 | 0.18 | 385.97 | -1.07 | 0.285 | 0.809 | 256 | 132 |
| Subiculum | -0.04 | 0.14 | 387.81 | -0.26 | 0.798 | 0.878 | 256 | 132 |
| Presubiculum | 0.15 | 0.14 | 387.39 | 1.05 | 0.294 | 0.809 | 256 | 132 |
| Parasubiculum | 0.22 | 0.18 | 384.03 | 1.23 | 0.220 | 0.809 | 256 | 132 |
| HATA | 0.07 | 0.17 | 388.00 | 0.42 | 0.677 | 0.854 | 256 | 132 |
| Fimbria | 0.10 | 0.16 | 386.77 | 0.62 | 0.533 | 0.854 | 256 | 132 |
| Molecular Layer | -0.05 | 0.14 | 387.48 | -0.39 | 0.699 | 0.854 | 256 | 132 |
| Dentate | -0.09 | 0.15 | 387.89 | -0.58 | 0.561 | 0.854 | 256 | 132 |
| **C. Females** | | | | | | | | |
| | beta | SE | df | t | p | Q | N, PTSD | N, No PTSD |
| CA1 | -0.06 | 0.12 | 398.00 | -0.50 | 0.619 | 0.824 | 245 | 153 |
| CA4 | -0.15 | 0.13 | 397.87 | -1.20 | 0.232 | 0.660 | 245 | 153 |
| CA3 | -0.21 | 0.14 | 398.00 | -1.51 | 0.133 | 0.660 | 245 | 153 |
| Tail | -0.14 | 0.14 | 397.89 | -0.99 | 0.321 | 0.706 | 245 | 153 |
| Subiculum | -0.03 | 0.11 | 396.98 | -0.26 | 0.796 | 0.824 | 245 | 153 |
| Presubiculum | 0.03 | 0.11 | 396.04 | 0.22 | 0.824 | 0.824 | 245 | 153 |
| Parasubiculum | 0.09 | 0.14 | 396.36 | 0.63 | 0.531 | 0.824 | 245 | 153 |
| HATA | -0.04 | 0.15 | 397.79 | -0.29 | 0.772 | 0.824 | 245 | 153 |

| | beta | SE | df | t | p | Q | N, PTSD | N, No PTSD |
|---|---|---|---|---|---|---|---|---|
| Fimbria | 0.20 | 0.14 | 395.72 | 1.49 | 0.136 | 0.660 | 245 | 153 |
| Molecular Layer | -0.09 | 0.12 | 397.35 | -0.75 | 0.452 | 0.824 | 245 | 153 |
| Dentate | -0.15 | 0.12 | 397.83 | -1.18 | 0.240 | 0.660 | 245 | 153 |
| **D. Males** | | | | | | | | |
| | beta | SE | df | t | p | Q | N, PTSD | N, No PTSD |
| CA1 | -0.02 | 0.11 | 199.53 | -0.21 | 0.835 | 0.964 | 148 | 189 |
| CA4 | 0.17 | 0.11 | 187.39 | 1.54 | 0.126 | 0.528 | 148 | 189 |
| CA3 | 0.18 | 0.12 | 191.67 | 1.45 | 0.148 | 0.528 | 148 | 189 |
| Tail | 0.17 | 0.13 | 140.83 | 1.31 | 0.192 | 0.528 | 148 | 189 |
| Subiculum | 0.04 | 0.12 | 140.99 | 0.32 | 0.752 | 0.964 | 148 | 189 |
| Presubiculum | -0.02 | 0.11 | 337.00 | -0.16 | 0.876 | 0.964 | 148 | 189 |
| Parasubiculum | 0.00 | 0.13 | 200.28 | 0.02 | 0.986 | 0.986 | 148 | 189 |
| HATA | 0.02 | 0.12 | 262.15 | 0.20 | 0.843 | 0.964 | 148 | 189 |
| Fimbria | -0.07 | 0.13 | 240.42 | -0.49 | 0.625 | 0.964 | 148 | 189 |
| Molecular Layer | 0.07 | 0.11 | 209.46 | 0.64 | 0.526 | 0.964 | 148 | 189 |
| Dentate | 0.16 | 0.11 | 213.91 | 1.47 | 0.143 | 0.528 | 148 | 189 |
| **E. Male Military** | | | | | | | | |
| | beta | SE | df | t | p | Q | N, PTSD | N, No PTSD |
| CA1 | 0.08 | 0.12 | 144.97 | 0.65 | 0.516 | 0.631 | 122 | 178 |
| CA4 | 0.28 | 0.11 | 122.43 | 2.50 | 0.014* | 0.099^ | 122 | 178 |
| CA3 | 0.26 | 0.12 | 112.92 | 2.13 | 0.036* | 0.099^ | 122 | 178 |
| Tail | 0.27 | 0.13 | 128.76 | 2.12 | 0.036* | 0.099^ | 122 | 178 |
| Subiculum | 0.12 | 0.12 | 103.27 | 0.97 | 0.334 | 0.567 | 122 | 178 |
| Presubiculum | -0.01 | 0.11 | 300.00 | -0.10 | 0.917 | 0.917 | 122 | 178 |
| Parasubiculum | -0.11 | 0.12 | 101.51 | -0.89 | 0.375 | 0.567 | 122 | 178 |
| HATA | 0.10 | 0.12 | 220.99 | 0.82 | 0.412 | 0.567 | 122 | 178 |
| Fimbria | -0.04 | 0.15 | 191.20 | -0.26 | 0.796 | 0.876 | 122 | 178 |
| Molecular Layer | 0.16 | 0.11 | 147.34 | 1.45 | 0.150 | 0.330 | 122 | 178 |
| Dentate | 0.26 | 0.11 | 146.87 | 2.33 | 0.021* | 0.099^ | 122 | 178 |
| **F. Female Civilians** | | | | | | | | |
| | beta | SE | df | t | p | Q | N, PTSD | N, No PTSD |
| CA1 | -0.003 | 0.16 | 350.42 | -0.02 | 0.984 | 0.897 | 230 | 121 |
| CA4 | -0.12 | 0.17 | 350.58 | -0.74 | 0.462 | 0.897 | 230 | 121 |
| CA3 | -0.23 | 0.18 | 349.69 | -1.29 | 0.198 | 0.897 | 230 | 121 |
| Tail | -0.15 | 0.18 | 349.36 | -0.83 | 0.408 | 0.897 | 230 | 121 |
| Subiculum | -0.004 | 0.15 | 350.92 | -0.03 | 0.977 | 0.897 | 230 | 121 |
| Presubiculum | 0.17 | 0.15 | 350.55 | 1.16 | 0.246 | 0.897 | 230 | 121 |
| Parasubiculum | 0.19 | 0.19 | 346.97 | 1.00 | 0.320 | 0.897 | 230 | 121 |

| | | | | | | | | |
|---|---|---|---|---|---|---|---|---|
| HATA | 0.07 | 0.19 | 350.97 | 0.36 | 0.720 | 0.897 | 230 | 121 |
| Fimbria | 0.07 | 0.17 | 349.73 | 0.42 | 0.676 | 0.897 | 230 | 121 |
| Molecular Layer | -0.05 | 0.15 | 350.88 | -0.34 | 0.734 | 0.897 | 230 | 121 |
| Dentate | -0.10 | 0.16 | 350.92 | -0.63 | 0.532 | 0.897 | 230 | 121 |

*Note.* Analyses conducted separately in civilians and military included sex covariates (sex, age*sex, and $age^2$*sex), and analyses conducted separately in males in females included a covariate for civilian/military status. Q-values depict FDR-corrected significance, values in bold ink passed the FDR threshold. ^ = 0.05<q<0.10, * = 0.01<q<0.05, ** = 0.001<q<0.01, ***=q<0.001.

**ST30. Interactions between C-PTSD and depression in males and military in composite subregions**

| | Beta | SE | df | t | p | q |
|---|---|---|---|---|---|---|
| | | | **A. Males** | | | |
| CA1/sub | -0.33 | 0.10 | 1325.72 | -3.17 | 0.002** | **0.018*** |
| Complete dentate | -0.13 | 0.10 | 1317.26 | -1.28 | 0.202 | 0.202 |
| HP | -0.26 | 0.10 | 1321.16 | -2.54 | 0.011* | **0.025*** |
| CA-only | -0.23 | 0.10 | 1319.35 | -2.28 | 0.023* | **0.030*** |
| HP/DG | -0.21 | 0.10 | 1318.33 | -2.12 | 0.035* | **0.039*** |
| Sub-complex | -0.26 | 0.11 | 1335.30 | -2.36 | 0.018* | **0.027*** |
| CA/SubPresub/MOL | -0.28 | 0.10 | 1323.49 | -2.76 | 0.006** | **0.025*** |
| HF | -0.25 | 0.10 | 1323.61 | -2.46 | 0.014* | **0.025*** |
| Hippo extended | -0.26 | 0.10 | 1327.10 | -2.63 | 0.009** | **0.025*** |
| | | | **B. Military** | | | |
| | Beta | SE | df | t | p | q |
| CA1/sub | -0.35 | 0.12 | 1080.63 | -3.00 | 0.003** | **0.027*** |
| Complete dentate | -0.14 | 0.11 | 1075.18 | -1.25 | 0.211 | 0.211 |
| HP | -0.27 | 0.11 | 1078.14 | -2.38 | 0.017* | **0.042*** |
| CA-only | -0.24 | 0.11 | 1076.88 | -2.14 | 0.033* | **0.042*** |
| HP/DG | -0.22 | 0.11 | 1076.03 | -2.01 | 0.045* | 0.051^ |
| Sub-complex | -0.27 | 0.12 | 1079.67 | -2.20 | 0.028* | **0.042*** |
| CA/SubPresub/MOL | -0.29 | 0.11 | 1079.56 | -2.63 | 0.009** | **0.041*** |
| HF | -0.25 | 0.11 | 1080.95 | -2.23 | 0.026* | **0.042*** |
| Hippo extended | -0.26 | 0.11 | 1084.69 | -2.34 | 0.019* | **0.042*** |

*Note*. Q-values depict FDR-corrected significance, values in bold ink passed the FDR threshold. ^ = 0.05<q<0.10, * = 0.01<q<0.05, ** = 0.001<q<0.01, ***=q<0.001. Descriptions: CA1/sub=CA1 + subiculum, Complete dentate=CA4+DG, HP (hippocampus proper)=CA1+CA2/3+CA4, CA-only= CA1+CA2/3, HP/DG= CA1+CA2/3+DG, Sub-complex=subiculum + presubiculum + parasubiculum, CA/SubPreSub/MOL= CA1 + CA2/3 + subiculum + presubiculum + MOL, HF (hippocampal formation) = CA1 + CA2/3 + CA4 + DG + subiculum + tail, hippo extended= CA1 + CA2/3 + CA4 + DG + subiculum + presubiculum + parasubiculum + tail + MOL + HATA + fimbria.

**ST31. Pairwise comparisons of PTSD*Depression interactions after removing childhood trauma**

| ROI | Pairwise Groups | beta | SE | df | t | p | q | Group1 | Group2 |
|---|---|---|---|---|---|---|---|---|---|
| CA1 | C-PTSD+Depression - Depression-only | 0.35 | 0.25 | 373.00 | 1.42 | 0.157 | 0.314 | **32** | 12 |
| | C-PTSD+Depression - C-PTSD-only | -0.10 | 0.17 | 375.52 | -0.57 | 0.569 | 0.726 | 32 | 40 |
| | C-PTSD+Depression - Control | -0.05 | 0.14 | 372.65 | -0.35 | 0.726 | 0.726 | 32 | 293 |
| | Depression-only - C-PTSD-only | -0.45 | 0.24 | 366.86 | -1.90 | 0.059^ | 0.183 | 12 | 40 |
| | Depression-only - Control | -0.40 | 0.21 | 370.70 | -1.88 | 0.061^ | 0.183 | 12 | 293 |
| | C-PTSD-only - Control | 0.05 | 0.12 | 376.74 | 0.40 | 0.686 | 0.726 | 40 | 293 |
| CA1/Sub | C-PTSD+Depression - Depression-only | 0.40 | 0.24 | 374.67 | 1.65 | 0.099^ | 0.198 | **32** | 12 |
| | C-PTSD+Depression - C-PTSD-only | -0.02 | 0.17 | 376.37 | -0.11 | 0.915 | 0.940 | 32 | 40 |
| | C-PTSD+Depression - Control | -0.03 | 0.13 | 376.49 | -0.20 | 0.841 | 0.940 | 32 | 293 |
| | Depression-only - C-PTSD-only | -0.42 | 0.23 | 369.34 | -1.80 | 0.072^ | 0.198 | 12 | 40 |
| | Depression-only - Control | -0.43 | 0.21 | 372.66 | -2.06 | 0.041* | 0.198 | 12 | 293 |
| | C-PTSD-only - Control | -0.01 | 0.12 | 376.89 | -0.08 | 0.940 | 0.940 | 40 | 293 |
| HP | C-PTSD+Depression - Depression-only | 0.33 | 0.25 | 370.74 | 1.32 | 0.189 | 0.378 | **32** | 12 |
| | C-PTSD+Depression - C-PTSD-only | -0.07 | 0.17 | 374.32 | -0.39 | 0.694 | 0.833 | 32 | 40 |
| | C-PTSD+Depression - Control | -0.01 | 0.14 | 357.50 | -0.07 | 0.941 | 0.941 | 32 | 293 |
| | Depression-only - C-PTSD-only | -0.40 | 0.24 | 365.29 | -1.67 | 0.096^ | 0.342 | 12 | 40 |
| | Depression-only - Control | -0.34 | 0.21 | 365.41 | -1.58 | 0.114 | 0.342 | 12 | 293 |
| | C-PTSD-only - Control | 0.06 | 0.13 | 350.33 | 0.46 | 0.644 | 0.833 | 40 | 293 |
| CA-only | C-PTSD+Depression - Depression-only | 0.32 | 0.25 | 371.16 | 1.27 | 0.205 | 0.410 | **32** | 12 |
| | C-PTSD+Depression - C-PTSD-only | -0.09 | 0.17 | 374.57 | -0.49 | 0.625 | 0.798 | 32 | 40 |
| | C-PTSD+Depression - Control | -0.03 | 0.14 | 352.50 | -0.22 | 0.826 | 0.826 | 32 | 293 |
| | Depression-only - C-PTSD-only | -0.40 | 0.24 | 366.09 | -1.69 | 0.092^ | 0.315 | 12 | 40 |
| | Depression-only - Control | -0.35 | 0.21 | 366.70 | -1.63 | 0.105 | 0.315 | 12 | 293 |
| | C-PTSD-only - Control | 0.05 | 0.13 | 335.93 | 0.43 | 0.665 | 0.798 | 40 | 293 |
| HP/DG | C-PTSD+Depression - Depression-only | 0.35 | 0.25 | 370.39 | 1.38 | 0.167 | 0.334 | **32** | 12 |
| | C-PTSD+Depression - C-PTSD-only | -0.05 | 0.17 | 374.18 | -0.31 | 0.754 | 0.905 | 32 | 40 |
| | C-PTSD+Depression - Control | 0.01 | 0.14 | 359.92 | 0.05 | 0.958 | 0.958 | 32 | 293 |
| | Depression-only - C-PTSD-only | -0.40 | 0.24 | 364.70 | -1.68 | 0.093^ | 0.334 | 12 | 40 |
| | Depression-only - Control | -0.34 | 0.21 | 364.52 | -1.58 | 0.114 | 0.334 | 12 | 293 |
| | C-PTSD-only - Control | 0.06 | 0.13 | 357.46 | 0.49 | 0.621 | 0.905 | 40 | 293 |
| CA/SubPresub/MOL | C-PTSD+Depression - Depression-only | 0.38 | 0.24 | 372.07 | 1.60 | 0.111 | 0.222 | **32** | 12 |
| | C-PTSD+Depression - C-PTSD-only | 0.00 | 0.16 | 375.49 | -0.01 | 0.991 | 0.991 | 32 | 40 |
| | C-PTSD+Depression - Control | -0.01 | 0.13 | 371.54 | -0.08 | 0.936 | 0.991 | 32 | 293 |
| | Depression-only - C-PTSD-only | -0.38 | 0.23 | 364.32 | -1.68 | 0.094^ | 0.222 | 12 | 40 |
| | Depression-only - Control | -0.39 | 0.20 | 369.14 | -1.91 | 0.057^ | 0.222 | 12 | 293 |
| | C-PTSD-only - Control | -0.01 | 0.12 | 376.78 | -0.08 | 0.940 | 0.991 | 40 | 293 |

| | | | | | | | | | |
|---|---|---|---|---|---|---|---|---|---|
| HF | C-PTSD+Depression - Depression-only | 0.39 | 0.25 | 370.80 | 1.56 | 0.120 | 0.240 | **32** | 12 |
| | C-PTSD+Depression - C-PTSD-only | -0.03 | 0.17 | 374.40 | -0.19 | 0.850 | 0.915 | 32 | 40 |
| | C-PTSD+Depression - Control | 0.01 | 0.14 | 369.71 | 0.11 | 0.915 | 0.915 | 32 | 293 |
| | Depression-only - C-PTSD-only | -0.42 | 0.24 | 367.32 | -1.77 | 0.078^ | 0.240 | 12 | 40 |
| | Depression-only - Control | -0.37 | 0.21 | 367.30 | -1.75 | 0.082^ | 0.240 | 12 | 293 |
| | C-PTSD-only - Control | 0.05 | 0.13 | 371.99 | 0.38 | 0.704 | 0.915 | 40 | 293 |
| Hippo Extended | C-PTSD+Depression - Depression-only | 0.43 | 0.24 | 370.78 | 1.75 | 0.080^ | 0.160 | **32** | 12 |
| | C-PTSD+Depression - C-PTSD-only | -0.02 | 0.17 | 374.70 | -0.11 | 0.910 | 0.916 | 32 | 40 |
| | C-PTSD+Depression - Control | 0.01 | 0.14 | 370.52 | 0.11 | 0.916 | 0.916 | 32 | 293 |
| | Depression-only - C-PTSD-only | -0.45 | 0.23 | 367.68 | -1.92 | 0.056^ | 0.160 | 12 | 40 |
| | Depression-only - Control | -0.41 | 0.21 | 368.05 | -1.97 | 0.049* | 0.160 | 12 | 293 |
| | C-PTSD-only - Control | 0.03 | 0.12 | 371.44 | 0.27 | 0.784 | 0.160 | 40 | 293 |

*Note*. Q-values depict FDR-corrected significance, values in bold ink passed the FDR threshold. ^ = 0.05<q<0.10, * = 0.01<q<0.05, ** = 0.001<q<0.01, ***=q<0.001. Text in gray ink denotes pairwise comparisons in composite regions that did not survive FDR at the omnibus level. Descriptions: CA1/sub=CA1 + subiculum, Complete dentate=CA4+DG, HP (hippocampus proper)=CA1+CA2/3+CA4, CA-only= CA1+CA2/3, HP/DG= CA1+CA2/3+DG, Sub-complex=subiculum + presubiculum + parasubiculum, CA/SubPreSub/MOL= CA1 + CA2/3 + subiculum + presubiculum + MOL, HF (hippocampal formation) = CA1 + CA2/3 + CA4 + DG + subiculum + tail, hippo extended= CA1 + CA2/3 + CA4 + DG + subiculum + presubiculum + parasubiculum + tail + MOL + HATA + fimbria. C-PTSD+Depression, n=621, C-PTSD-only, n=384, Depression-only, n=138, healthy control, n=1120

| ST32. Meta-analysis of C-PTSD, depression, and C-PTSD interactions | | | | |
|---|---|---|---|---|
| **A. C-PTSD (n=989) vs. Controls (n=1361)** | | | | |
| | Cohen's d | SE | p | q |
| Tail | -0.10 | 0.05 | 0.063^ | 0.231 |
| Subiculum | -0.06 | 0.05 | 0.178 | 0.392 |
| CA1 | -0.12 | 0.06 | 0.056^ | 0.231 |
| Presubiculum | -0.07 | 0.05 | 0.098^ | 0.270 |
| Parasubiculum | -0.02 | 0.05 | 0.726 | 0.799 |
| Molecular layer | -0.08 | 0.05 | 0.059^ | 0.231 |
| Dentate | -0.06 | 0.05 | 0.215 | 0.394 |
| CA3 | -0.002 | 0.05 | 0.967 | 0.967 |
| CA4 | -0.05 | 0.05 | 0.298 | 0.468 |
| Fimbria | -0.02 | 0.05 | 0.655 | 0.799 |
| HATA | -0.03 | 0.05 | 0.519 | 0.714 |
| **B. Depression (n=760) vs. Control (n=1369)** | | | | |
| | Cohen's d | SE | p.value | q.value |
| Tail | -0.12 | 0.05 | 0.013* | 0.083^ |
| Subiculum | -0.01 | 0.05 | 0.786 | 0.786 |
| CA1 | -0.12 | 0.05 | 0.015* | 0.083^ |
| Presubiculum | -0.02 | 0.05 | 0.729 | 0.786 |
| Parasubiculum | 0.03 | 0.06 | 0.627 | 0.766 |
| Molecular layer | -0.09 | 0.05 | 0.071^ | 0.204 |
| Dentate | -0.08 | 0.05 | 0.094^ | 0.204 |
| CA3 | -0.08 | 0.05 | 0.111 | 0.204 |
| CA4 | -0.08 | 0.05 | 0.092^ | 0.204 |
| Fimbria | 0.04 | 0.06 | 0.448 | 0.616 |
| HATA | -0.04 | 0.05 | 0.391 | 0.614 |
| **C. C-PTSD x Depression (n= 324) vs. control (n=1154)** | | | | |
| | Cohen's d | SE | p.value | q.value |
| Tail | -0.13 | 0.06 | 0.038* | 0.139 |
| Subiculum | -0.08 | 0.06 | 0.202 | 0.370 |
| CA1 | -0.15 | 0.06 | 0.007** | 0.077^ |
| Presubiculum | -0.07 | 0.07 | 0.362 | 0.498 |
| Parasubiculum | -0.01 | 0.06 | 0.869 | 0.869 |
| Molecular layer | -0.13 | 0.06 | 0.020* | 0.110 |
| Dentate | -0.10 | 0.06 | 0.071^ | 0.178 |
| CA3 | -0.08 | 0.07 | 0.245 | 0.385 |
| CA4 | -0.10 | 0.06 | 0.081^ | 0.178 |
| Fimbria | 0.02 | 0.07 | 0.812 | 0.869 |
| HATA | -0.05 | 0.07 | 0.412 | 0.504 |

*Note*. Q-values depict FDR-corrected significance, values in bold ink passed the FDR threshold. ^ = 0.05<q<0.10, * = 0.01<q<0.05, ** = 0.001<q<0.01, ***=q<0.001.

**ST33. Meta-analysis of pairwise comparisons of depression by C-PTSD interactions**

| | Cohen's d | SE | p | q | N cases | N controls | Total N |
|---|---|---|---|---|---|---|---|
| **A. C-PTSD + Depression vs. Depression-only** | | | | | | | |
| Tail | -0.20 | 0.11 | 0.072^ | 0.158 | 339 | 137 | 476 |
| Subiculum | -0.24 | 0.11 | 0.031* | 0.114 | 339 | 137 | 476 |
| CA1 | -0.32 | 0.14 | 0.025* | 0.114 | 339 | 137 | 476 |
| Presubiculum | -0.21 | 0.11 | 0.056^ | 0.154 | 339 | 137 | 476 |
| Parasubiculum | -0.17 | 0.11 | 0.122 | 0.192 | 339 | 137 | 476 |
| Molecular layer | -0.32 | 0.13 | 0.016* | 0.114 | 339 | 137 | 476 |
| Dentate | -0.22 | 0.13 | 0.092^ | 0.169 | 339 | 137 | 476 |
| CA3 | -0.15 | 0.16 | 0.321 | 0.321 | 339 | 137 | 476 |
| CA4 | -0.20 | 0.14 | 0.148 | 0.192 | 339 | 137 | 476 |
| Fimbria | -0.15 | 0.11 | 0.157 | 0.192 | 339 | 137 | 476 |
| HATA | -0.12 | 0.11 | 0.280 | 0.308 | 339 | 137 | 476 |
| **B. C-PTSD + Depression vs. C-PTSD-only** | | | | | | | |
| | Cohen's d | SE | p | q | N cases | N controls | Total N |
| Tail | -0.12 | 0.08 | 0.110 | 0.673 | 506 | 298 | 804 |
| Subiculum | 0.02 | 0.08 | 0.781 | 0.781 | 506 | 298 | 804 |
| CA1 | -0.10 | 0.08 | 0.173 | 0.673 | 506 | 298 | 804 |
| Presubiculum | 0.04 | 0.08 | 0.642 | 0.781 | 506 | 298 | 804 |
| Parasubiculum | 0.05 | 0.09 | 0.580 | 0.781 | 506 | 298 | 804 |
| Molecular layer | -0.07 | 0.08 | 0.356 | 0.673 | 506 | 298 | 804 |
| Dentate | -0.07 | 0.08 | 0.367 | 0.673 | 506 | 298 | 804 |
| CA3 | -0.09 | 0.09 | 0.336 | 0.673 | 506 | 298 | 804 |
| CA4 | -0.07 | 0.08 | 0.333 | 0.673 | 506 | 298 | 804 |
| Fimbria | 0.05 | 0.09 | 0.584 | 0.781 | 506 | 298 | 804 |
| HATA | -0.03 | 0.09 | 0.720 | 0.781 | 506 | 298 | 804 |
| **C. C-PTSD + Depression vs. Control** | | | | | | | |
| | Cohen's d | SE | p | q | N cases | N controls | Total N |
| Tail | -0.13 | 0.06 | 0.038* | 0.139 | 573 | 1044 | 1617 |
| Subiculum | -0.08 | 0.06 | 0.202 | 0.370 | 573 | 1044 | 1617 |
| CA1 | -0.15 | 0.06 | 0.007** | 0.077^ | 573 | 1044 | 1617 |
| Presubiculum | -0.07 | 0.07 | 0.362 | 0.498 | 573 | 1044 | 1617 |
| Parasubiculum | -0.01 | 0.06 | 0.869 | 0.869 | 573 | 1044 | 1617 |
| Molecular layer | -0.13 | 0.06 | 0.020* | 0.110 | 573 | 1044 | 1617 |

| | Cohen's d | SE | p | q | N cases | N controls | Total N |
|---|---|---|---|---|---|---|---|
| Dentate | -0.10 | 0.06 | 0.071^ | 0.178 | 573 | 1044 | 1617 |
| CA3 | -0.08 | 0.07 | 0.245 | 0.385 | 573 | 1044 | 1617 |
| CA4 | -0.10 | 0.06 | 0.081^ | 0.178 | 573 | 1044 | 1617 |
| Fimbria | 0.02 | 0.07 | 0.812 | 0.869 | 572 | 1044 | 1616 |
| HATA | -0.05 | 0.07 | 0.412 | 0.504 | 573 | 1044 | 1617 |
| | **D. C-PTSD-only vs. Depression-only** | | | | | | |
| | Cohen's d | SE | p | q | N cases | N controls | Total N |
| Tail | 0.13 | 0.12 | 0.303 | 0.978 | 202 | 114 | 316 |
| Subiculum | -0.01 | 0.15 | 0.933 | 0.978 | 202 | 114 | 316 |
| CA1 | 0.02 | 0.14 | 0.872 | 0.978 | 202 | 114 | 316 |
| Presubiculum | -0.07 | 0.12 | 0.599 | 0.978 | 202 | 114 | 316 |
| Parasubiculum | -0.17 | 0.12 | 0.174 | 0.978 | 202 | 114 | 316 |
| Molecular layer | -0.01 | 0.13 | 0.919 | 0.978 | 202 | 114 | 316 |
| Dentate | 0.00 | 0.12 | 0.978 | 0.978 | 202 | 114 | 316 |
| CA3 | -0.07 | 0.12 | 0.590 | 0.978 | 202 | 114 | 316 |
| CA4 | 0.01 | 0.13 | 0.929 | 0.978 | 202 | 114 | 316 |
| Fimbria | -0.09 | 0.13 | 0.480 | 0.978 | 202 | 114 | 316 |
| HATA | -0.06 | 0.12 | 0.609 | 0.978 | 202 | 114 | 316 |
| | **E. Depression-only vs. Control** | | | | | | |
| | Cohen's d | SE | p | q | N cases | N controls | Total N |
| Tail | 0.03 | 0.10 | 0.793 | 0.965 | 137 | 852 | 989 |
| Subiculum | 0.13 | 0.10 | 0.194 | 0.580 | 137 | 852 | 989 |
| CA1 | 0.07 | 0.14 | 0.638 | 0.965 | 137 | 852 | 989 |
| Presubiculum | 0.14 | 0.11 | 0.211 | 0.580 | 137 | 852 | 989 |
| Parasubiculum | 0.19 | 0.10 | 0.059* | 0.429 | 137 | 852 | 989 |
| Molecular layer | 0.09 | 0.12 | 0.435 | 0.957 | 137 | 852 | 989 |
| Dentate | 0.04 | 0.10 | 0.714 | 0.965 | 137 | 852 | 989 |
| CA3 | 0.00 | 0.10 | 0.999 | 0.999 | 137 | 852 | 989 |
| CA4 | 0.02 | 0.10 | 0.877 | 0.965 | 137 | 852 | 989 |
| Fimbria | 0.17 | 0.10 | 0.078 | 0.429 | 137 | 852 | 989 |
| HATA | -0.02 | 0.10 | 0.821 | 0.965 | 137 | 852 | 989 |
| | **F. C-PTSD-only vs. control** | | | | | | |
| | Cohen's d | SE | p | q | N cases | N controls | Total N |
| Tail | 0.05 | 0.07 | 0.429 | 0.771 | 325 | 1016 | 1341 |
| Subiculum | 0.04 | 0.07 | 0.597 | 0.771 | 325 | 1016 | 1341 |
| CA1 | 0.05 | 0.10 | 0.620 | 0.771 | 325 | 1016 | 1341 |
| Presubiculum | -0.01 | 0.07 | 0.885 | 0.771 | 325 | 1016 | 1341 |
| Parasubiculum | 0.03 | 0.07 | 0.631 | 0.771 | 325 | 1016 | 1341 |
| Molecular layer | 0.05 | 0.08 | 0.500 | 0.771 | 325 | 1016 | 1341 |

| | | | | | | | |
|---|---|---|---|---|---|---|---|
| Dentate | 0.08 | 0.07 | 0.257 | 0.771 | 325 | 1016 | 1341 |
| CA3 | 0.14 | 0.07 | 0.042* | 0.771 | 325 | 1016 | 1341 |
| CA4 | 0.09 | 0.07 | 0.206 | 0.771 | 325 | 1016 | 1341 |
| Fimbria | 0.01 | 0.07 | 0.910 | 0.771 | 325 | 1016 | 1341 |
| HATA | 0.05 | 0.08 | 0.552 | 0.771 | 325 | 1016 | 1341 |

*Note*. Q-values depict FDR-corrected significance. ^ = 0.05<q<0.10, * = 0.01<q<0.05, ** = 0.001<q<0.01, ***=q<0.001.

Main Effect of Depression in the CA1

| | Study | D | SE |
|---|---|---|---|
| A | AMC | 0.25 | 0.25 |
| | Duke | -0.06 | 0.16 |
| | Durham | 0.15 | 0.47 |
| | Emory | -0.31 | 0.22 |
| | McLean | -0.7 | 0.3 |
| | NSW | -0.06 | 0.18 |
| | Umichigan | -0.04 | 0.33 |
| | Yale | 0.09 | 0.32 |
| | **Subtotal** | **-0.09** | **0.08** |
| B | Chicago | -0.53 | 0.48 |
| | Columbia | -0.33 | 0.23 |
| | Groningen | -0.37 | 0.39 |
| | INTRUST | -0.24 | 0.14 |
| | MPLS | -0.11 | 0.17 |
| | Munster | -0.08 | 0.32 |
| | SouthDakota1 | 0.16 | 0.23 |
| | SouthDakota2 | 0.02 | 0.39 |
| | Stellenbosch | -0.06 | 0.13 |
| | UMSL | -0.13 | 0.25 |
| | VA_AnnArbor | -0.88 | 0.29 |
| | VETSA | 0.02 | 0.16 |
| | Wisconsin | 0.22 | 0.31 |
| | **Subtotal** | **-0.13** | **0.06** |
| | **Total** | **-0.12** | **0.05** |

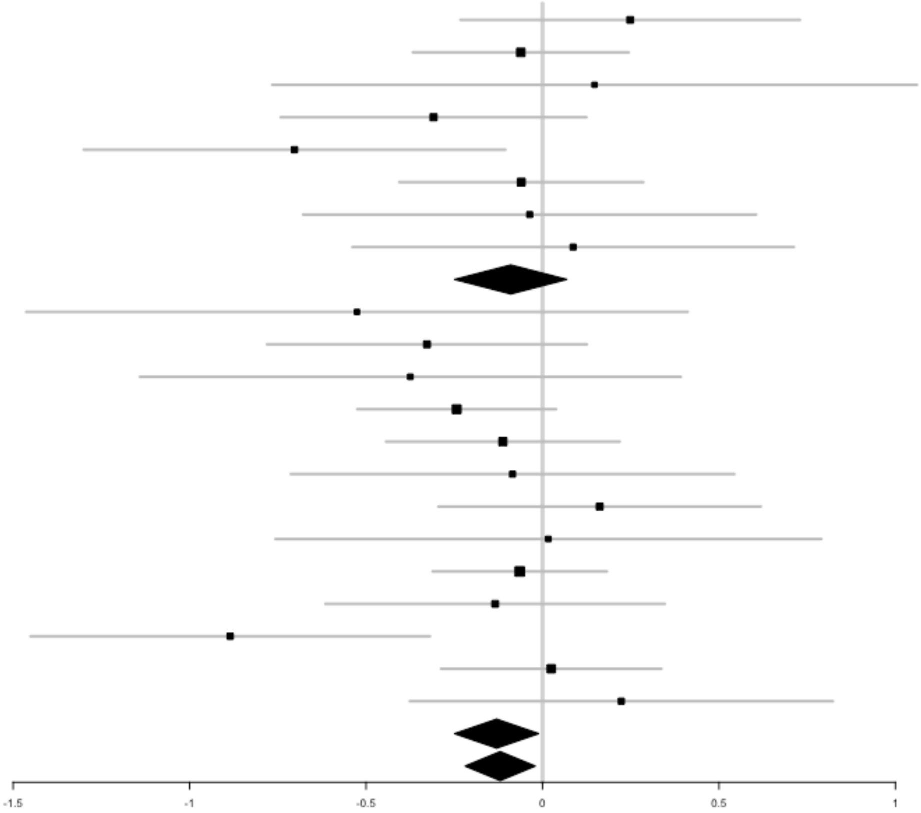


**SF3. Forest Plots of Depression in the CA1.** Forest plots show the heterogeneity in Cohen's D effect sizes across cohorts. Sub-sample meta-analyses of Cohen's d were computed for cohorts participating in our earlier meta-analysis by Logue et al. (2018; panel A), and new cohorts that were only included in the present study (panel B).

Main Effect of Depression in the Tail

| | Study | D | Tail SE |
|---|---|---|---|
| A | AMC | 0.33 | 0.25 |
| | Duke | 0.02 | 0.16 |
| | Durham | -0.99 | 0.49 |
| | Emory | -0.36 | 0.22 |
| | McLean | -0.59 | 0.3 |
| | NSW | -0.05 | 0.18 |
| | Umichigan | 0.08 | 0.33 |
| | Yale | 0.13 | 0.32 |
| | **Subtotal** | **-0.1** | **0.11** |
| B | Chicago | 0.22 | 0.47 |
| | Columbia | -0.28 | 0.23 |
| | Groningen | -0.75 | 0.4 |
| | INTRUST | -0.05 | 0.14 |
| | MPLS | -0.12 | 0.17 |
| | Munster | -0.27 | 0.32 |
| | SouthDakota1 | 0.18 | 0.23 |
| | SouthDakota2 | -0.44 | 0.4 |
| | Stellenbosch | -0.06 | 0.13 |
| | UMSL | -0.46 | 0.25 |
| | VA_AnnArbor | -0.75 | 0.29 |
| | VETSA | -0.06 | 0.16 |
| | Wisconsin | -0.05 | 0.3 |
| | **Subtotal** | **-0.14** | **0.06** |
| | **Total** | **-0.12** | **0.05** |

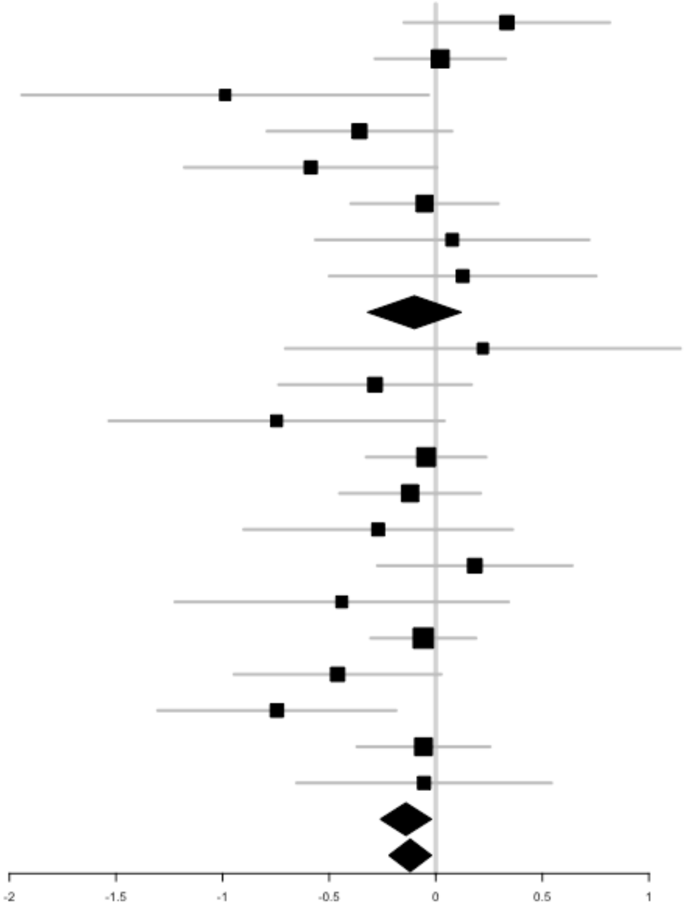


**SF4. Forest Plots of Depression in the hippocampal tail.** Forest plots show the heterogeneity in Cohen's D effect sizes across cohorts. Sub-sample meta-analyses of Cohen's d were computed for cohorts participating in our earlier meta-analysis by Logue et al. (2018; panel A), and new cohorts that were only included in the present study (panel B).

Subgroup Effect of C-PTSD in the CA1 Among Individuals with Depression

| | Study | D | SE |
|---|---|---|---|
| | AMC | -0.15 | 0.4 |
| A | Duke | -1.02 | 0.39 |
| | Emory | -0.27 | 0.41 |
| | NSW | -0.82 | 0.36 |
| | **Subtotal** | **-0.59** | **0.21** |
| B | INTRUST | -0.47 | 0.32 |
| | MPLS | 0.25 | 0.29 |
| | Stellenbosch | 0.15 | 0.27 |
| | VETSA | -0.31 | 0.28 |
| | Wisconsin | -0.55 | 0.34 |
| | **Subtotal** | **-0.16** | **0.16** |
| | **Total** | **-0.32** | **0.14** |

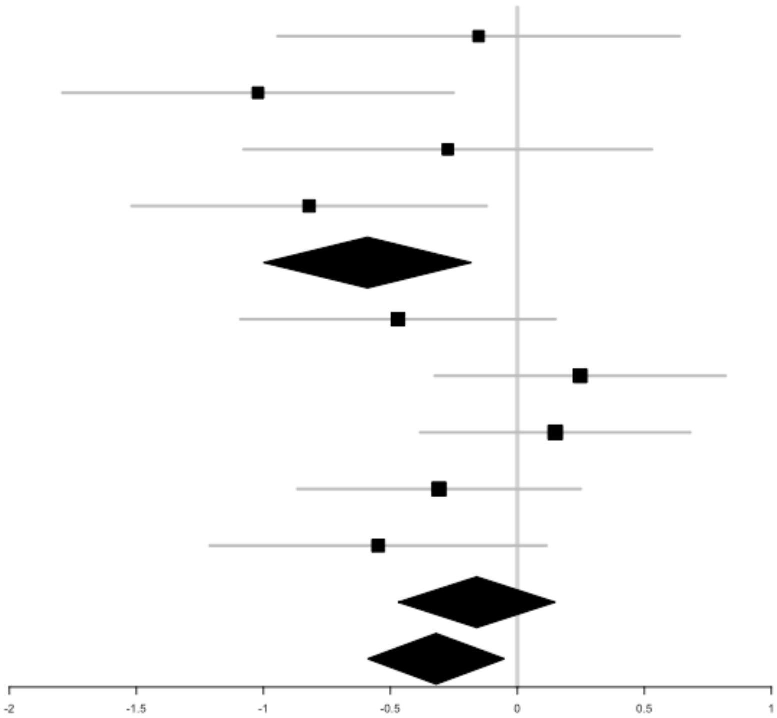


**SF5. Forest Plots of C-PTSD in the CA1 of depressed individuals.** Forest plots show the heterogeneity in Cohen's D effect sizes across cohorts. Sub-sample meta-analyses of Cohen's d were computed for cohorts participating in our earlier meta-analysis by Logue et al. (2018; panel A), and new cohorts that were only included in the present study (panel B).

Subgroup Effect of Depression in the CA1 Among Individuals with C-PTSD

| | Study | D | SE |
|---|---|---|---|
| A | AMC | 0.07 | 0.33 |
| | Duke | -0.26 | 0.24 |
| | Emory | -0.3 | 0.31 |
| | NSW | -0.31 | 0.24 |
| | Yale | 1.18 | 0.52 |
| | **Subtotal** | **-0.13** | **0.14** |
| B | Chicago | -0.53 | 0.48 |
| | Columbia | 0.06 | 0.41 |
| | Groningen | -0.37 | 0.39 |
| | INTRUST | -0.26 | 0.25 |
| | MPLS | 0.18 | 0.25 |
| | Munster | 0.14 | 0.46 |
| | SouthDakota1 | -0.23 | 0.27 |
| | SouthDakota2 | 0.58 | 0.46 |
| | Stellenbosch | -0.1 | 0.21 |
| | UMSL | -0.07 | 0.3 |
| | VETSA | -0.13 | 0.35 |
| | **Subtotal** | **-0.09** | **0.09** |
| | **Total** | **-0.1** | **0.08** |

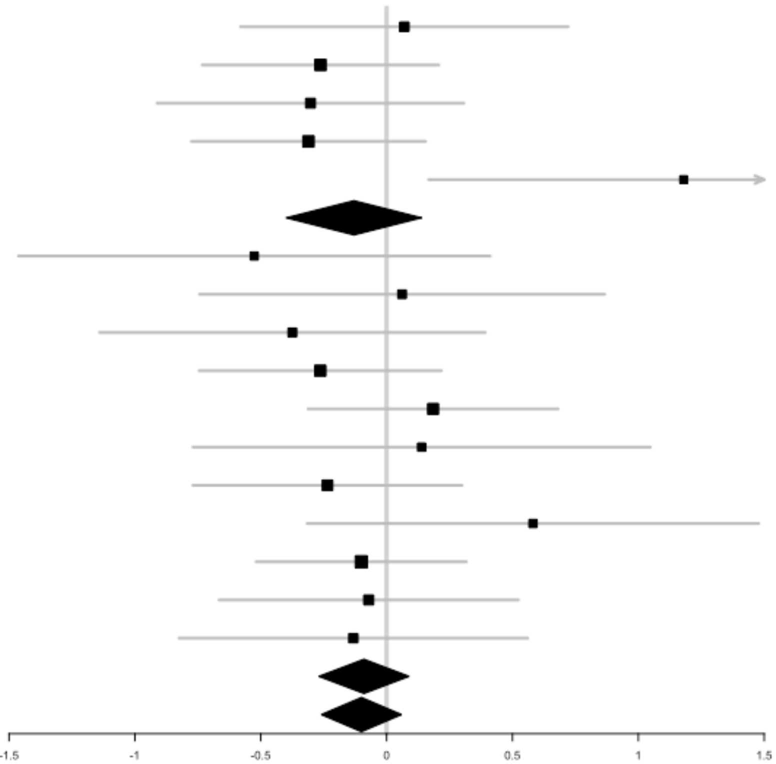


**SF6. Forest Plots of Depression in the CA1 in individuals with C-PTSD.** Forest plots show the heterogeneity in Cohen's D effect sizes across cohorts. Sub-sample meta-analyses of Cohen's d were computed for cohorts participating in our earlier meta-analysis by Logue et al. (2018; panel A), and new cohorts that were only included in the present study (panel B).
